# Genomics and adaptive divergence of the allopolyploid grass *Brachypodium hybridum* in a pangenotypic framework

**DOI:** 10.64898/2026.08.21.746290

**Authors:** Miguel Campos, Guohong Albert Wu, Li Lei, Rubén Sancho, Bruno Contreras-Moreira, Ernesto Perez-Collazos, John P. Vogel, Pilar Catalán

**Author notes:** Corresponding author: Pilar Catalán Rodríguez. Departamento de Ciencias Agrarias y del Medio Natural. Escuela Politécnica Superior de Huesca. Universidad de Zaragoza. C/ Carretera de Cuarte Km 1. E-22071 Huesca. Spain.

## Abstract

- Natural allopolyploids with multiple origins are powerful systems for analyzing genome evolution; however, population-level whole-genome studies of wild-type recurrent polyploids remain scarce.
- We investigated the origins and evolutionary dynamics of the allotetraploid grass *Brachypodium hybridum* and its diploid progenitors (*B. distachyon*, *B. stacei*) by combining whole-genome sequencing of 307 accessions from across the circum-Mediterranean region with phylogenomics, population genomics, and comparative subgenomic analyses.
- Nuclear and plastome phylogenies reveal three independent allopolyploidization events: an ancient Iberian origin (∼1.78 Ma) and two more recent origins in the western (∼0.56 Ma) and eastern (∼0.24 Ma) Mediterranean. Each subgenome (D and S) evolved independently with minimal recombination. All *B. hybridum* lineages carry higher deleterious loads than their diploid progenitors, and the Ancient lineage carries a disproportionately heavy burden, particularly in the S subgenome. Population structure identifies three genetic groups; while the Ancient lineage remained isolated, recent western and eastern lineages exchanged migrants in the eastern Mediterranean contact zone.
- *Brachypodium hybridum* exemplifies how recurrent allopolyploidization, minimal subgenomic recombination, and environmental filtering generate and maintain genetic diversity, establishing it as a model for polyploid evolution and ecological adaptation in grasses worldwide.

## 1. Introduction

Hybridization and whole-genome duplication are among the most important forces shaping plant diversification, speciation, and adaptation (Ramsey & Schemske, 1998, 2002; Soltis & Soltis, 1999; Soltis *et al*., 2015; Wendel *et al*., 2018; Mason & Wendel, 2020). In allopolyploids, two divergent parental genomes are brought together in a single nucleus, creating immediate opportunities for reproductive isolation, ecological expansion, subgenomic interaction, and novel evolutionary trajectories (Levy & Feldman, 2002; Soltis *et al*., 2016; Wendel *et al*., 2018). Yet allopolyploid species follow strikingly different paths after formation. Some undergo rapid genomic restructuring, biased fractionation, transposable element activation, or subgenome dominance, whereas others retain remarkably stable parental genome architecture over millions of years (Bird *et al*., 2018; Mason & Wendel, 2020; Scarlett *et al*., 2023). Understanding why some allopolyploids become dynamic genomic mosaics while others remain structurally stable is therefore a central question in plant evolutionary genomics.

A second unresolved question concerns the evolutionary consequences of recurrent allopolyploid formation. Recurrent origins are now recognized as common in plants and can generate substantial standing variation by sampling different parental populations (Soltis & Soltis, 1999; Soltis *et al*., 2015; Wendel *et al*., 2018). Independent origins may produce distinct lineage-specific combinations of parental variation, increase ecological amplitude, and create repeated opportunities for adaptation (Symonds *et al*., 2010; Te Beest *et al*., 2012; Mason & Wendel, 2020). However, recurrently formed lineages may differ strongly in age, geographic range, demographic history, progenitor identity, and post-origin gene flow. In young systems such as *Tragopogon miscellus*, recurrent polyploid formation has been directly documented from extant progenitor species (Soltis & Soltis, 1999; Chester *et al*., 2012). In selfing allopolyploids such as *Arabidopsis suecica*, whole-genome resequencing rejected a strict single-origin model and demonstrated that multiple founding individuals contributed to the species (Novikova *et al*., 2017; Burns *et al*., 2021). In older allopolyploids, however, one or more parental lineages may be extinct, unsampled, or genetically diverged from extant relatives, making it difficult to distinguish inherited parental legacy from post-polyploid evolution (Wendel *et al*., 2018; Mason & Wendel, 2020).

A third major challenge is to understand how deleterious variation evolves after allopolyploidization. Whole-genome duplication can mask recessive deleterious mutations, reduce the efficiency of purifying selection, and allow mildly deleterious variants to persist, particularly in selfing or demographically restricted lineages (Wright *et al*., 2013; Paape *et al*., 2018; Kryvokhyzha *et al*., 2019; Conover & Wendel, 2022). Deleterious-load dynamics may differ between subgenomes depending on the genetic load inherited from each progenitor, subsequent effective population size, recombination, selection, and drift (Conover & Wendel, 2022). Few natural systems permit deleterious variants to be compared simultaneously among diploid progenitors and multiple independently derived allopolyploid lineages differing in age. Such comparisons are essential for determining whether subgenomic mutation load reflects steady post-polyploid accumulation, inherited “legacy load” from parental lineages, or lineage-specific demographic histories.

Addressing these three questions together requires a combination of features that is rarely available in a single natural system: multiple independent origins spanning deep and recent timescales, well-characterized extant diploid progenitors, and population-level whole-genome data. Crop allopolyploids such as bread wheat and *Brassica napus* offer deep genomic resources but represent single-origin or domestication-filtered systems (Pont *et al*., 2019; Avni *et al*., 2022). Synthetic polyploids and young natural systems such as *Tragopogon* provide temporal resolution but lack the long-term evolutionary trajectories needed to assess genome stability and mutation-load dynamics (Chester *et al*., 2012). Conversely, older allopolyploids often lack identifiable extant progenitors, precluding direct progenitor–derivative comparisons (Wendel *et al*., 2018; Mason & Wendel, 2020). Population genomic analyses based on whole-genome sequencing have dramatically improved our ability to infer life-history traits, demographic history, adaptation, and genome evolution at unprecedented resolution (Exposito-Alonso *et al*., 2019; Alonge *et al*., 2020; Varshney *et al*., 2021). These approaches have been widely applied in major crop systems, including maize, wheat, barley, and wheat relatives, revealing extensive introgression, local adaptation, and haplotype differentiation across environmental gradients (Hufford *et al*., 2012, 2021; Takuno *et al*., 2015; Zhou *et al*., 2021; Avni *et al*., 2022; Guo *et al*., 2025). However, comparable population-level genomic studies remain limited in wild allopolyploid systems, especially those involving naturally occurring, recurrently formed lineages with different ages and degrees of progenitor resolution (Baduel *et al*., 2018; Wendel *et al*., 2018; Song *et al*., 2023). This gap limits our ability to generalize from crop systems and synthetic polyploids to the evolutionary dynamics of wild allopolyploids in natural landscapes.

The annual *Brachypodium distachyon*–*B. stacei*–*B. hybridum* complex offers an exceptional natural system for addressing these questions. The diploids *B. distachyon* and *B. stacei* are the known progenitor species of the allotetraploid *B. hybridum*, and the complex occurs across the circum-Mediterranean region, where strong geographic, climatic, and edaphic gradients provide opportunities for local adaptation (Catalán *et al*., 2012, 2016a,b; López-Alvarez *et al*., 2015; Sancho *et al*., 2018; Campos *et al*., 2024, 2026). Previous genomic studies have shown that *B. hybridum* originated recurrently from distinct hybridization and polyploidization events, including one ancient lineage and two more recent lineages (Gordon *et al*., 2020; Scarlett *et al*., 2023; Mu *et al*., 2023a). They have also shown that *B. hybridum* exhibits low subgenome dominance, limited homeologous exchange, and high structural stability, suggesting an unusually stable allopolyploid genome (Scarlett *et al*., 2023; Mu *et al*., 2023a). However, previous whole-genome studies were based on more limited sampling and therefore could not fully resolve the population-level consequences of recurrent origin, lineage divergence, chromosomal admixture, deleterious-load evolution, or isolation by geography and environment.

Here, we use whole-genome resequencing of 307 accessions from the annual *Brachypodium* complex to investigate how recurrent allopolyploid formation has shaped the evolutionary diversification of *B. hybridum*. Our study integrates nuclear syntenic SNPs, subgenome-specific phylogenomics, plastome comparisons, deleterious-variant prediction, chromosomal local ancestry inference, genomic differentiation scans, and environmental association analyses. This design allows us to compare the two diploid progenitor species with the D and S subgenomes of *B. hybridum,* and to evaluate three independently originated *B. hybridum* lineages spanning different temporal and geographic scales. Specifically, we aim to: (i) resolve the number of independent origins of *B. hybridum* at population-genomic scale, and characterize their age, geographic distribution, and progenitor affinity; (ii) assess whether independently originated allopolyploid lineages remain genomically isolated or exchange nuclear and plastid genomes upon secondary contact; (iii) quantify del-eterious-variant accumulation across the D and S subgenomes relative to their diploid progenitors, and contrast genetic load between older and younger lineages; (iv) test whether genomic islands of differentiation are enriched for functional categories associated with development, stress response, phenology, and environmental adaptation; and (v) disentangle the relative contributions of geography and environment to genetic differentiation within and among *B. hybridum* lineages.

By addressing these questions, this study uses the *Brachypodium* complex as a model for understanding the broader evolutionary consequences of recurrent allopolyploidization in wild plants. We show that *B. hybridum* combines three features rarely examined together in a single natural system: multiple origins across deep and recent timescales, strong nuclear subgenomic stability despite cytonuclear discordance and limited secondary admixture and contrasting subgenomic trajectories of deleterious-variant accumulation. These features make *B. hybridum* a powerful system for testing how recurrent origin, parental legacy, selfing, drift, selection, and environmental filtering interact during the diversification of wild allopolyploids.

## 2. Material & Methods

### 2.1. Plant Material

A total of 307 *B. distachyon - B. stacei - B. hybridum* (Bd-Bs-Bh) complex accessions distributed across its native circum-Mediterranean region and some invaded areas were selected for this study (Table **S1**, Fig. **1A**). The sampling included 113 individuals of *B. distachyon*, 65 of *B. stacei* and 129 of *B. hybridum*. Seeds were obtained from 307 natural populations (Table **S1**). Plants were grown under standard greenhouse conditions with a 16-hour light period at 22°C, 50-60% relative humidity, and natural light supplemented with artificial lighting when necessary in different institutions [High Polytechnic School of Huesca - University of Zaragoza (Spain), Joint Genome Institute (USA)] at different years, to avoid any potential outbreeding of samples, following the procedures indicated in Catalán et al. (2012).

**Figure 1.**
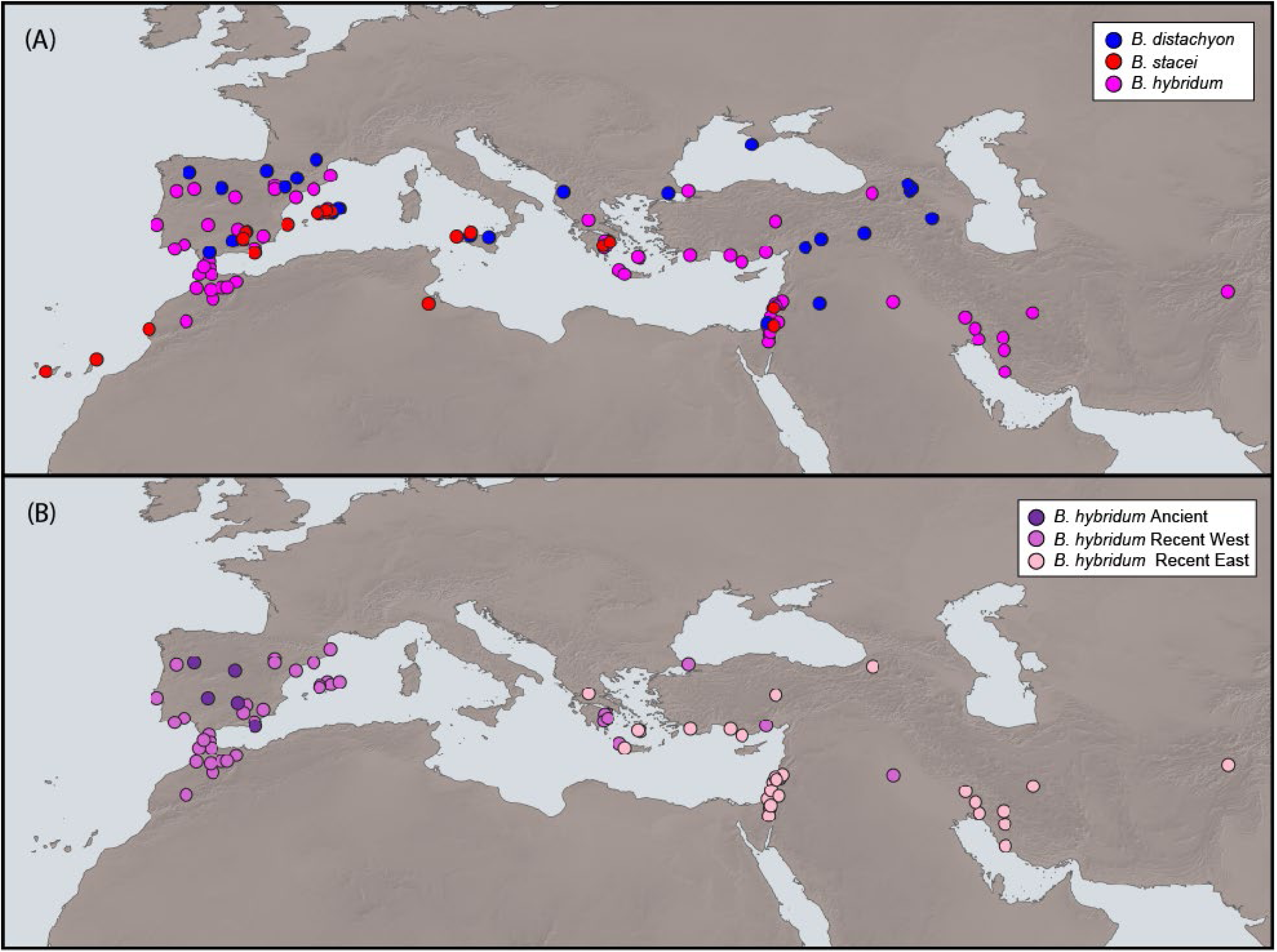
Circum-Mediterranean geographic distribution of population samples of the three annual Brachypodium species under study **(A)**: B. distachyon (n = 113; blue), B. stacei (n = 65; red) and B. hybridum (n = 125; pink). **(B)** Distribution of the three independent B. hybridum lineages: Ancient (n = 5; mauve), Recent West (n = 59; wine) and Recent East (n = 61; light pink). (see Table **S1**). Of the total of 129 individuals of B. hybridum, four individuals from non-native areas (1 Australia, 2 South Africa, 1 Uruguay; see Table **S1**) were excluded for this figure, all four belonging to the Recent West group (n=63) (see Results).

### 2.2. Whole-genome sequencing and syntenic SNP (Single Homeologous Polymorphisms) calling

Total genomic DNA was obtained from leaf tissue using CTAB protocol with modifications (Doyle & Doyle, 1987). Whole-genome sequence data were generated at the DOE Joint Genome Institute (JGI) for 146 samples using Illumina technology (4 *B. distachyon*, 29 *B. stacei*, 113 *B. hybridum*; Table **S1**).

An Illumina library per sample was constructed and sequenced using the Illumina NovaSeq S4 platform (2×151) and duplicate reads were removed based on paired sequence matching using Clumpify in BBMap v39.33 (Bushnell, 2014). BBDuk v38.96 (BBMap suite) was used to trim adapter sequences and terminal G-homopolymers (≥5), perform right-end quality trimming (Q < 6), and remove reads that contained ’N’ bases, had an average quality score < 6, or were shorter than 49 bp (33% of the full read length).

Paired-end Illumina read sequences of all *Brachypodium* samples under study (Table **S1**) were mapped on a multiple progenitor-reference genome generated by concatenating the chromosomes of the reference genomes of *B. distachyon* (Bd21 v3_2; https://phytozome-next.jgi.doe.gov/info/Bdistachyon_v3_2) and *B. stacei* (ABR114 v2_1; https://phytozome-next.jgi.doe.gov/info/Bstacei_v2_1) using minimap2 v2.28 (Li, 2018). Individuals with over 90% of reads mapped to *B. distachyon* chromosomes were assigned to this species, and same for *B. stacei*. Those displaying approximately 50% values to each progenitor genome were classified as *B. hybridum* (Table **S2**). One sample of *Oryza sativa* (GCA_034140825.1) was added to the study and used as outgroup in phylogenomic analysis. The variant calling process was conducted for each individual and the data were subsequently merged into a unified variant calling format (VCF) file of 308 samples (307 *Brachypodium* accessions plus *Oryza sativa*) using bcftools v1.18 (Danecek *et al*., 2021). This VCF file containing information from whole genomes’ SNPs was employed to filter single homeologous polymorphisms (SHP), equivalent to syntenic SNPs, using the AlloSHP pipeline (https://github.com/eead-csic-compbio/AlloSHP) developed by Sancho et al. (2025). Briefly, it combines the strategies of simultaneous mapping against multiple reference genomes and syntenic alignment of these genomes to call SHPs (∼SNPs), using as input data a single VCF file and the concatenated reference genomes of the progenitor species. The pipeline generated two sequence data sets for each individual, one containing the *B. distachyon* genome syntenic SNP positions and the other the *B. stacei* positions. For *B. hybridum*, both data sets were retained (two sequences per accession, labelled, respectively, D and S), while for the progenitor species samples the data set from the absent genome was discarded (a single sequence per accession). The initial alignment contained an average of 496,556,626 positions for *B. hybridum*, 264,826,759 positions for *B. distachyon*, 252,427,179 positions for *B. stacei*, and 12,331,508 positions for *Oryza sativa*.

The full syntenic SNP alignment data matrix was filtered using trimAl v1.5 (Capella-Gutiérrez *et al*., 2009) to remove low-quality SNPs or those with gaps, resulting in a final matrix of 7,964,944 informative SNP positions without gaps (Bd-Bs-Bh complex dataset #1; Table **S2**). Next, to assess subgenomic differences within *B. hybridum*, the SNP matrix obtained before the filtering step was subdivided into two datasets, one containing SNP data from *B. hybridum* subgenome D samples (*B. hybridum*-D dataset #2; 528,470 SNPs after filtering), and the other SNP data from *B. hybridum* subgenome S samples (*B. hybridum*-S dataset #3; 455,524 SNPs after filtering). First, these subgenomic VCFs were filtered for data quality using a MAF = 0.05 threshold and no missing data allowed. Then, they were subsequently filtered to avoid LD using the SNPRelate v1.38.1 package in R (Zheng *et al*., 2012) using a 10 kb window and a r² threshold of 0.2. Furthermore, a total *B. hybridum* dataset was created using the full genome data of *B. hybridum* through the concatenation of datasets #2 and #3 (*B. hybridum* dataset #4; 983,994 SNPs). The four separate datasets were employed for different downstream analyses.

### 2.3. *Brachypodium* complex (Bd-Bs-Bh) evolutionary analyses

#### 2.3.1. Phylogenomic trees

The full data matrix obtained from syntenic SNPs retrieved from all studied individuals (ca. 8M SNPs; Bd-Bs-Bh dataset #1) was used to reconstruct a whole nuclear-based phylogeny of *B. distachyon* and *B. stacei* genomic samples and *B. hybridum* subgenomic samples (D and S subgenomes). A complete maximum-likelihood (ML) phylogenomic tree was constructed using IQtree2 v2.0.7 (Nguyen *et al*., 2015) and 1000 ultrafast bootstrap (UFBoot) replicates. Furthermore, a multispecies coalescence (MSC) tree was constructed by dividing this syntenic SNP matrix into 10,000 bp windows, building ML trees for each window, and then using Weighted-ASTRAL included in ASTER package v.1-3.5 (Zhang et al., 2025) which was fed with the individual ML gene trees to reconstruct the species tree. The topology of this MSC tree was compared to that of the ML tree to detect potential incongruences and to assess the potential influence of ILS.

#### 2.3.2. Genomic diversity

Genomic diversity metrics (heterozygosity, Fis, selfing rate) were calculated for each species, and for the three independent *B. hybridum* lineages separately, using plink2 v2.00a5.12LM (Chang *et al*., 2015) in the initial VCF file. We applied a genotype-filtering criteria discarding sites with read depth outside 0.5xMode–2xMode, and any heterozygous call (0/1 or 0|1) with an alternative-allele fraction outside 30–70% was recoded as missing. Using this filtering, we then estimated genetic-diversity parameters via two approaches. First, we estimated per-SNP heterozygosity, defined as the proportion of heterozygous genotypes (0/1) at variable sites across the syntenic SNP matrix, averaged per individual. Second, to mitigate bias from uneven SNP discovery, we calculated genome-wide heterozygosity across all callable genomic sites (both variant and invariant positions). In highly inbred diploid samples, this genome-wide heterozygosity approaches zero, whereas per-SNP heterozygosity may still yield higher values because it is restricted to polymorphic loci. In addition, we calculated nucleotide diversity (π) and Tajima’s D for each species and lineage using VCFtools v0.1.16 (Danecek *et al*., 2011), based on non-overlapping 10kb genomic windows. Statistical differences among groups were assessed through Kruskal–Wallis and pairwise Wilcoxon tests with Benjamini–Hochberg correction. Finally, mean heterozygosity, π, Tajima’ D, Fis, and selfing rate were compared among species and lineages by one-way Kruskal-Wallis with Wilcoxon pairwise post hoc tests to identify statistically significant differences. To avoid overinterpretation due to the large number of windows and potential pseudoreplication in π and Tajima’s D, we quantified the effect size (η² for global tests and r for pairwise comparisons) to determine whether statistically significant differences were biologically relevant. To infer recent gene flow among lineages of *B. hybridum* with different origins, we estimated pairwise Wright’s Fst and Slatkin’s linearized Fst (Fst/(1–Fst) in Arlequin v3.5.2.2 (Excoffier & Lischer, 2010), and converted Fst to the number of migrants per generation (Nm) via the formula Fst≈1/(4 Nm + 1).

#### 2.3.3. Potentially deleterious variants

To identify potentially deleterious variants in the nuclear genomes from the three species, we combined the variant annotation tool SnpEff v5.2.f (Cingolani, 2012) with the likelihood-based predictor BAD_Mutations (Kono *et al*., 2018). SnpEff was used to annotate genetic variants as single nucleotide polymorphisms (SNPs), insertions, deletions, and multiple nucleotide polymorphisms (MNPs). These were then analyzed by transcript and amino acid position to obtain substitution data, which served as input for BAD_Mutations. This program applies a likelihood ratio test (dN < dS) to predict whether amino acid substitutions are likely to be deleterious, based on phylogenetically informed site models. First, SnpEff was used to characterize the genomic distribution and functional categories of all detected variants (intronic, intergenic, upstream, downstream, missense, synonymous, and UTR). Variants located in upstream, downstream, and intergenic regions were excluded, and only those within CDS regions were retained as potentially deleterious. These gene variants were quantified per individual and averaged at the species and *B. hybridum*-lineage levels to avoid bias caused by unequal sample sizes. To account for differences in genome size between subgenomes, total counts of potential and confirmed deleterious variants were further normalized as variants per megabase of coding sequence (variants/Mb CDS), thereby enabling direct comparisons of potentially deleterious variant load across subgenomes and species.

All gene variants previously annotated as potentially deleterious by SnpEff were subsequently analyzed with BAD_Mutations to determine which were significantly constrained (p < 0.05), thus identifying the subset of truly deleterious amino acid changes. The BAD_Mutations workflow was set up using *B. distachyon* and *B. stacei* as reference species for allotetraploid and progenitor species analysis, respectively, with homologous sequences recovered and aligned between related Poaceae species to provide evolutionary context.

### 2.4. *Brachypodium hybridum* population genomics and phylogenomics

#### 2.4.1. Phylogenomic analyses

Phylogenomic analyses of *B. hybridum* were carried out in three complementary steps. First, we reconstructed maximum-likelihood (ML) trees for each subgenome separately (D and S) using IQtree2 v2.1.3 (Nguyen *et al*., 2015) on datasets #2 (D subgenome) and #3 (S subgenome). Branch support was assessed with 1,000 ultrafast bootstrap replicates (-bb 1000). To formally test whether the two subgenome topologies were significantly different, we performed generalized Robinson–Foulds (gRF) metric (Smith, 2020) to test tree-wide distances. The two resulting subgenome trees were then compared in a tanglegram using a custom R script (CoPhylo.R), which mirrors tip labels and highlights topological concordance. Second, to analyze potential cytonuclear discordances between nuclear-versus plastome-based phylogenies, we built an ML tree from the concatenated genome-wide SNP matrix (dataset #4) under the same IQ-TREE settings and contrasted it with the *B. hybridum* plastome tree (Campos *et al*., 2026). To quantify and visualize topological incongruence between nuclear and plastome trees, we implemented the Procrustean Approach to Cophylogeny (PACo) in R (Balbuena *et al*., 2013). We generated 1,000 bootstrap trees for both nuclear and plastome alignments, then used PACo to superimpose the two sets of terminal positions via Procrustean superposition and computed the sum of squared residuals for each tip, serving as a concordance score. Residuals were binned into quartiles to gauge the magnitude of discordance (quartiles 3 and 4 indicating the strongest conflict) and plotted it using ggplot2 in R (Wickham, 2016). Pairwise generalized Robinson–Foulds (gRF) metric (Smith, 2020) was also computed to compare the topologies of the nuclear and plastome ML trees.

#### 2.4.2. Dating analysis

We also computed a Multi Species Coalescent (MSC) phylogeny of *B. hybridum* lineages using the combined (D+S) dataset #4 and the SVDQuartets method implemented in PAUP* (Swofford, 2003; Chifman & Kubatko, 2014), evaluating all possible quartets and performing 1,000 standard bootstrap replicates to assess node support. To estimate absolute divergence times of the *B. hybridum* splits, we applied TreePL (Smith & O’Meara, 2012), which implements a penalized-likelihood framework to accommodate rate heterogeneity among branches by optimizing a smoothing parameter via cross-validation. We imposed calibrations for the crown Ancient clade (2.16 Ma ± 0.5 Ma), the crown Recent West clade (0.72 Ma ± 0.1 Ma) and the crown Recent East clade (0.13 Ma ± 0.1 Ma) according to Mu et al. (2023a). To establish 95% confidence intervals around our age estimates, we ran TreePL separately on each of the 1,000 bootstrap replicate trees generated for the *B. hybridum* dataset. Continuous rate smoothing and calibration priors together yielded an ultrametric chronogram for each tree. All resulting dated trees were then summarized with TreeAnnotator implemented in BEAST2 v2.7.8 (Bouckaert *et al*., 2014) to calculate the 95% highest-posterior-density intervals for each node.

#### 2.4.3. Genomic structure and differentiation

We inferred *B. hybridum* population structure using the maximum-likelihood, Bayesian-inspired clustering algorithm implemented in ADMIXTURE v1.3.0 (Alexander & Lange, 2011) on our combined SNP dataset (dataset #4). ADMIXTURE jointly estimates individual ancestry proportions and allele frequencies via block-relaxation optimization across multilocus genotypes. We ran ten independent replicates for each K = 1–10, then identified the optimal number of clusters by minimizing the cross-validation error. Individual ancestry coefficients for the best K were summarized and visualized with Pophelper v2.3.1 in R (Francis, 2017), and spatial pie-chart maps were generated to display mean cluster membership at each sampling locality using custom script in R (AdmixturePieChart.R).

Furthermore, we used RFMix v2 (Maples *et al*., 2013) to delve into chromosomal-level positions and assess potential chromosomal admixture predominance from ancestral populations in individuals with high admixture levels. This software utilizes genomic data to infer local segmental ancestry in individuals, identifying regions of genetic mixing at the chromosomal level and assigning proportions of ancestry from distinct ancestral populations.

We quantified genome-wide patterns of absolute (Dxy) and relative (Fst) genomic differentiation using Pixy v1.2.6 (Korunes & Samuk, 2021) on dataset #4. Non-overlapping 10 kb windows were used to (i) compute pairwise Dxy as the absolute nucleotide divergence between each pair of main *B. hybridum* lineages and (ii) calculate Weir and Cockerham’s Fst as a relative measure of population differentiation among those lineages. The window estimates were then plotted chromosome-by-chromosome to identify regions of high divergence or low diversity. We applied a 95th-percentile cutoff to identify outlier windows in the Fst and Dxy analyses. Windows above this threshold were considered as candidate regions under selection, as they likely reflect non-neutral divergence between lineages. These SNPs were then mapped to the annotated progenitor-reference genome, retaining only those located in coding regions, as these are more likely to have functional impacts. We then searched for the Gene Ontology (GO) terms associated with these genes using the Ensembl dataset “bdistachyon_eg_gene” with the biomart v2.60.1 package in R (Durinck *et al*., 2005).

We conducted a principal coordinate analysis (PCoA) on 1,000 kb windows on dataset #4, to examine how genomic differences are distributed along the chromosomes among the main lineages of *B. hybridum*. We used a custom R script (WindowedPCoA.R) to perform this analysis, which allowed us to visualize the genomic variation in the principal axes, offering insights into the genomic structure and differentiation across the lineages.

#### 2.4.4. Impact of spatial isolation and environment

To explore patterns of isolation by distance (IBD) and isolation by environment (IBE) in *B. hybridum* populations, we used the dataset #4 and geographic coordinates and climate variables. First, we classified individuals into their corresponding lineages (Ancient, Recent West, Recent East) using the dartR v2.9.7 library (Mijangos *et al*., 2022). We then computed geographic distances between individuals using the Haversine method with the geosphere v1.5-20 package (Hijmans, 2026), and bioclimatic distances based on a set of 19 bioclimatic variables retrieved from WorldClim using stats R package (R Core Team, 2026). Genetic distances were estimated using pairwise Manhattan distances between SNPs using vegan R package (Oksanen *et al*., 2026).

To quantify the impact of geographic distance versus environmental variation on genetic differentiation in *B. hybridum* lineages, we performed distance-based redundancy analysis (dbRDA) using the capscale function in the vegan R package. We performed Principal Coordinates Analysis on each distance matrix to extract orthogonal axes for use as explanatory variables. We then fitted three dbRDA models: (1) a comprehensive model containing both sets of predictors, (2) an environment-only model testing isolation by environment (IBE), and (3) a space-only model testing isolation by distance (IBD). Model significance and the effect of individual PCoA axes were evaluated through permutation tests (10,000 permutations). To partition the distinct contributions of geography and climate, we conducted partial dbRDA, conditioning environmental predictors on geographic axes (and vice versa), thereby isolating IBE and IBD effects. Finally, variance partitioning and pairwise ANOVA comparisons were used to quantify the unique and shared proportions of genetic variation explained by spatial and environmental factors.

## 3. Results

### 3.1. *Brachypodium distachyon* – *B. stacei* – *B. hybridum* complex (Bd-Bs-Bh) evolutionary analyses

#### 3.1.1. Genome statistics, syntenic variant positions and linkage disequilibrium (LD) filtering

The mapping results revealed a high self-mapping percentage for *B. distachyon* individuals, with an average genomic coverage of 97.4% of the *B. distachyon* reference genome, and less than 1% mapped to *B. stacei.* The *B. stacei* individuals showed an average genomic coverage of 99.4% of their reference genome, and less than 1% mapped to *B. distachyon*. In contrast, the *B. hybridum* individuals exhibited nearly equal mapping coverage percentages to both genomes, with 96.6% of the *B. distachyon* genome covered and 98.2% of the *B. stacei* genome covered (Table **S2**).

After filtering out monomorphic positions and retaining only syntenic positions, the final alignment consisted of 15,505,655 SNPs for *B. hybridum* (7,718,627 for the subgenome D and 7,787,028 for the subgenome S), 7,811,129 for *B. distachyon*, 7,896,801 for *B. stacei*, and 367,184 for *Oryza sativa*. Thus, the resulting Bd-Bs-Bh complex dataset (dataset #1) consisted of ca. 8M syntenic SNPs. For the exclusive subgenomic-level analyses of *B. hybridum*, syntenic SNP filtering based on linkage disequilibrium (LD) was performed separately for the D and S subgenomes, retaining a total of 528,470 SNPs (*B. hybridum* subgenome D; dataset #2) and 455,524 SNPs (*B. hybridum* subgenome S, dataset #3), respectively. Overall, the combined VCF for both subgenomes (*B. hybridum* subgenomes D + S; dataset #4) had a total of 983,994 polymorphic positions for downstream analyses.

#### 3.1.2. Species tree, introgression and dating analysis

The MSC tree constructed using the syntenic SNP data matrix and *O. sativa* as outgroup revealed two major clades; the first clade (S-genome) comprised *B. stacei* individuals and the *B. hybridum*-S (subgenome S) samples, and the second clade (D-genome) *B. distachyon* individuals and the *B. hybridum*-D (subgenome D) samples (Figs. **2**, **S1A**). The quartet support values were generally high for the deep and shallow nodes of the best topology (q1) but showed similar values for the species tree and the two alternative topologies (q1, q2, q3) for the intermediate divergent nodes (Fig. **S1A**). Regarding the species tree phylogeny of *B. stacei* (Figs. **2**, **S1A**), two main sister lineages were identified; the first included ancestral populations from the Canary Islands, the Balearic Islands, and Sicily, and the second included other Mediterranean populations, the most recent being those from the Eastern Mediterranean, in line with previous findings (Campos *et al*., 2024). With respect to the phylogeny of *B. distachyon*, the initial divergence of the Sicilian lineage was followed by that of the EDF+ clade, separated into two subclades (Western Mediterranean, and predominantly Eastern Mediterranean), and that of the expanded S+T+ lineage, divided in turn into the S+ (Western Mediterranean, Spain mainly) and T+ (Eastern Mediterranean, Turkey mainly) subclades, also consistent with previous studies (Figs. **2**, **S1A**; Gordon *et al*., 2017; Sancho *et al*., 2018; Stritt *et al*., 2022).

**Figure 2.**
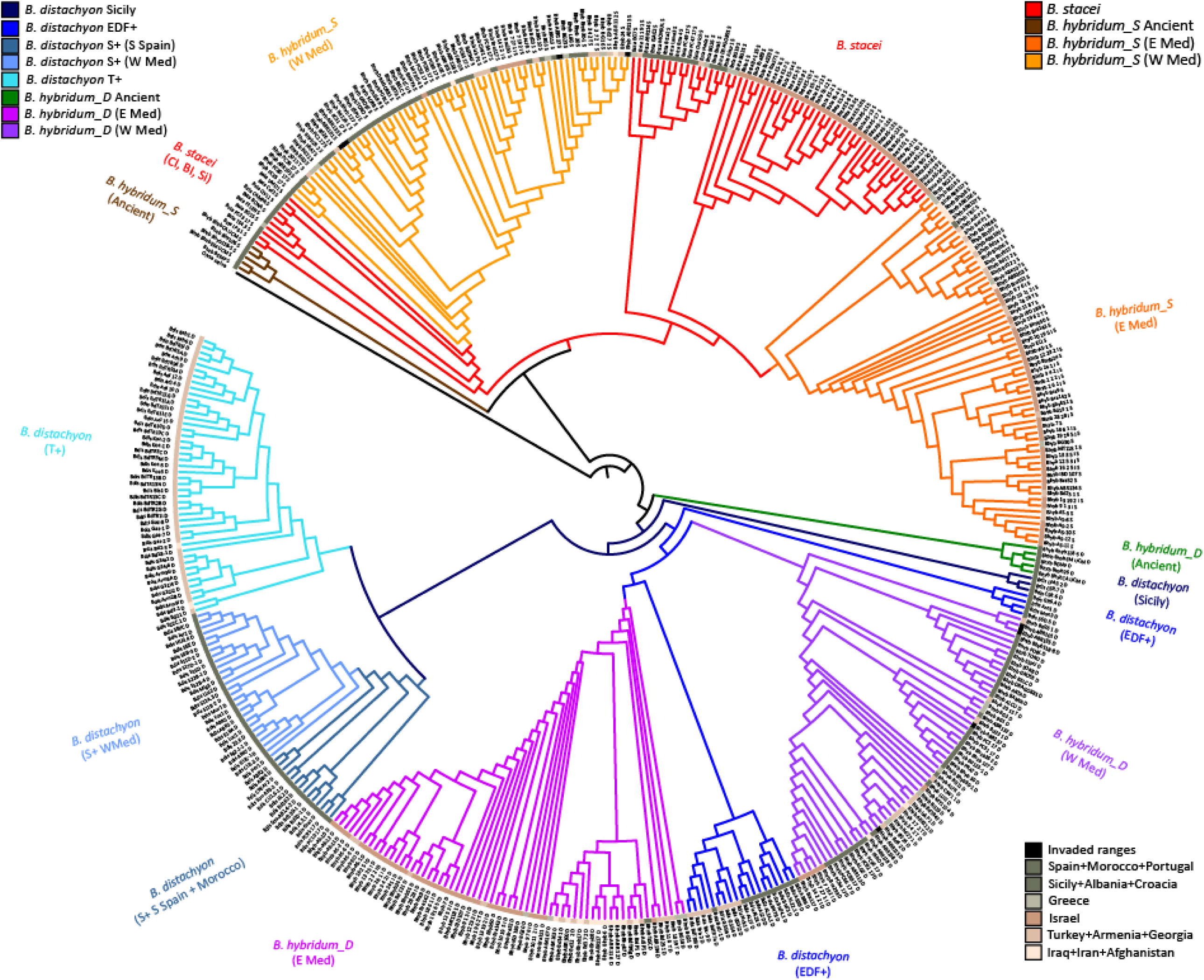
Multispecies coalescent tree of the Brachypodium annual complex samples inferred with Weighted-ASTRAL. Maximum-likelihood trees were estimated for each non-overlapping 10,000 bp windows of the full syntenic SNP matrix (∼8 million SNPs), and the resulting trees were combined under the ASTER package v.1-3.5 ASTRAL (Zhang et al. 2025) to reconstruct the MSC tree. Tips represent individual sequences: B. distachyon (n = 113), B. stacei (n = 65), and for B. hybridum two subgenomic sequences per accession (D and S subgenomes; n = 129 accessions). Branch colors denote genome/species/subgenome lineage identity and tip-colored squares correspond to geographic regions as indicated in the respective charts. Support values of nodal quartets are indicated in Fig. **S1A**.

The *B. hybridum* subgenomic clades exhibited evolutionary patterns reflecting their hybrid complexity, supporting the three independent origins of the allotetraploid (Figs. **2**, **S1A**; Mu *et al*., 2023a). In both subgenomic subtrees (S and D), the ancient lineage of *B. hybridum* diverged earlier than its corresponding progenitor species’ lineages, whereas the two most recently evolved *B. hybridum* lineages derived from their respective local progenitor lineages. Within these, the *B. hybridum* Western Mediterranean lineage originated from an old *B. stacei* lineage (Canary Islands, Sicily and Balearic Islands) (subgenome S) and the *B. distachyon* Western EDF+ lineage (subgenome D), whereas the *B. hybridum* Eastern Mediterranean lineage originated from the *B. stacei* Recent Eastern Mediterranean lineage (subgenome S) and the *B. distachyon* Eastern EDF+ lineage (subgenome D) (Figs. **2**, **S1A**). Four *B. hybridum* individuals from non-native areas (1 Australia, 2 South Africa, 1 Uruguay: Table **S1**) fall within the Recent West S and D lineages (Figs. **2**, **S1A**), supporting the recent western Mediterranean origins of these invaders.

Topological congruence between the MSC and the concatenated maximum likelihood (ML) trees was generally high (Figs. **2**, **S1**). The main difference concerned the relationships of the Western Mediterranean *B. hybridum*-D lineage, resolved as sister to the western *B. distachyon* EDF+ lineage in the MSC tree (Fig. **S1A**) but as a basal paraphyletic grade of the *B. distachyon* S+T+ clade in the ML tree (Fig. **S1B**).

The ancient allotetraploidization of *Brachypodium hybridum* was dated by TreePL to ∼1.78 Ma (Fig. **S2**), although no extant diploid progenitors have been identified for this earlier hybridization (Fig. **2**). In contrast, the estimated ages of the Recent West and Recent East lineages of *B. hybridum* were dated to ∼0.56 Ma and ∼0.24 Ma, respectively (Fig. **S2**). The two recent polyploidization events appear to have involved distinct geographic pools of the same genomic lineages of the D subgenome; both Recent West and Recent East *B. hybridum* groups trace their origin to the *B. distachyon* EDF+ clade, but to its western and eastern Mediterranean representatives, respectively (Fig. **2**). Regarding the S subgenome, the Recent West lineage derives from the old *B. stacei* populations of the western islands (Canary Islands, Balearic Islands and Sicily; Campos et al., 2024), while the Recent East lineage is linked to the more recently diverged *B. stacei* populations of the eastern Mediterranean (Fig. **2**).

#### 3.1.3. Diversity and inbreeding

Our analysis revealed that *B. hybridum* exhibited levels of SNP heterozygosity (0.0085 ± 0.0069) that were statistically comparable to those of its diploid progenitor species, *B. distachyon* (0.0093 ± 0.0033) and *B. stacei* (0.0090 ± 0.0012) (Table **1**). A similar pattern was observed for genome-wide (WG) heterozygosity, where *B. hybridum* (0.0004 ± 0.0000) was grouped with *B. distachyon* and showed higher values than *B*. *stacei*. Consistently, nucleotide diversity (π) and Tajima’s D followed parallel trends; *B. distachyon* displayed the highest π (0.0054 ± 0.0012), *B. stacei* the lowest (0.0031 ± 0.0013), and those of *B. hybridum* (0.0037 ± 0.0019) were intermediate between both progenitor species (Table **1**). Tajima’s D values were negative for *B. distachyon* (−0.686 ± 0.366) and *B. stacei* (−0.711 ± 0.547), but positive in *B. hybridum* (0.240 ± 0.770). As anticipated for highly autogamous species, all three taxa exhibited high inbreeding coefficients (Fis > 0.93), leading to a significant excess of homozygotes. Consequently, all three species were in significant Hardy-Weinberg disequilibrium (p < 0.05), a finding consistent with their predominant selfing reproductive strategies, which were reflected in their high selfing rates (>0.96) (Table **1**).

**Table 1.** Genomic diversity parameters (heterozygosity, nucleotide diversity (π), Tajima’s D, inbreeding coefficient (Fis), deviations from Hardy-Weinberg equilibrium (HWE), and selfing rate) of the Brachypodium distachyon-B. stacei-B. hybridum (Bd-Bs-Bh) complex at the species level, and separately for each subgenome (subgenome S, subgenome D) within B. hybridum and the three independent lineages of B. hybridum (Ancient, Recent West, Recent East). P-values from Hardy-Weinberg equilibrium tests are provided to assess deviations from equilibrium in each group and subgenome. Post-hoc Wilcoxon pairwise test letter assignments are provided for each index, indicating significant differences with p-values < 0.05. All pairwise comparisons for π and Tajima’s D were statistically significant; however, the assigned letters were determined according to both statistical significance and effect size (r for pairwise contrasts). Comparisons were made between the three species, between the progenitor species and the two subgenomes of B. hybridum, and between the three lineages of B. hybridum.

| Groups | N | SNPs Heterozygosity | WG Heterozygosity | $\pi$ | Tajima's D | Fis | HWE<br>p-Values | Selfing |
| --- | --- | --- | --- | --- | --- | --- | --- | --- |
| *B. distachyon* | 113 | 0.0093 ± 0.0033 <sup>AC</sup> | 0.0004 ± 0.0001 <sup>A</sup> | 0.0054 ± 0.0012 <sup>A</sup> | -0.686 ± 0.366 <sup>A</sup> | 0.939 ± 0.0213 <sup>AB</sup> | P < 0.05 | 0.968 ± 0.0114 <sup>AB</sup> |
| *B. stacei* | 65 | 0.0090 ± 0.0012 <sup>A</sup> | 0.0002 ± 0.0000 <sup>B</sup> | 0.0031 ± 0.0013 <sup>B</sup> | -0.711 ± 0.547 <sup>A</sup> | 0.941 ± 0.0761 <sup>B</sup> | P < 0.05 | 0.970 ± 0.0041 <sup>B</sup> |
| *B. hybridum* | 129 | 0.0085 ± 0.0069 <sup>A</sup> | 0.0004 ± 0.0000 <sup>A</sup> | 0.0037 ± 0.0019 <sup>BC</sup> | 0.240 ± 0.770 <sup>B</sup> | 0.937 ± 0.0052 <sup>A</sup> | P < 0.05 | 0.967 ± 0.0028 <sup>A</sup> |
| Subgenome S | 129 | 0.0061 ± 0.0071 <sup>B</sup> | 0.0003 ± 0.0070 <sup>C</sup> | 0.0027 ± 0.0010 <sup>B</sup> | 0.165 ± 0.842 <sup>B</sup> | 0.950 ± 0.0060 <sup>B</sup> | P < 0.05 | 0.974 ± 0.0031 <sup>C</sup> |
| Subgenome D | 129 | 0.0102 ± 0.0011 <sup>C</sup> | 0.0005 ± 0.0010 <sup>D</sup> | 0.0046 ± 0.0015 <sup>AC</sup> | 0.310 ± 0.688 <sup>B</sup> | 0.929 ± 0.0073 <sup>C</sup> | P < 0.05 | 0.963 ± 0.0041 <sup>D</sup> |
| Ancient | 5 | 0.0149 ± 0.0010 <sup>A</sup> | 0.0018 ± 0.0000 <sup>A</sup> | 0.0039 ± 0.0015 <sup>A</sup> | 0.805 ± 0.182 <sup>A</sup> | 0.8162 ± 0.0079 <sup>A</sup> | P < 0.05 | 0.893 ± 0.0054 <sup>A</sup> |
| Recent West | 63 | 0.0087 ± 0.0007 <sup>B</sup> | 0.0004 ± 0.0000 <sup>B</sup> | 0.0037 ± 0.0020 <sup>A</sup> | 0.506 ± 1.100 <sup>A</sup> | 0.935 ± 0.0052 <sup>B</sup> | P < 0.05 | 0.967 ± 0.0028 <sup>B</sup> |
| Recent East | 61 | 0.0081 ± 0.0005 <sup>C</sup> | 0.0004 ± 0.0000 <sup>C</sup> | 0.0033 ± 0.0013 <sup>A</sup> | -1.470 ± 0.720 <sup>B</sup> | 0.939 ± 0.0038 <sup>C</sup> | P < 0.05 | 0.969 ± 0.0020 <sup>C</sup> |

A more detailed analysis at the subgenomic level in *B. hybridum* revealed a more complex picture than a simple additive effect of its parental genomes. Our results showed an asymmetric pattern of diversity, with its S subgenome exhibiting a significant reduction in SNP heterozygosity (0.0061 ± 0.0071) compared to that of its progenitor species *B. stacei* (Table **1**). In contrast, the D subgenome showed a significant increase in diversity (0.0102 ± 0.0011) relative to that of its progenitor species *B. distachyon*. A similar asymmetric pattern was detected for nucleotide diversity (π) and Tajima’s D; π was lowest in the S subgenome (0.0027 ± 0.0010) and highest in the D subgenome (0.0046 ± 0.0015), while Tajima’s D values were positive in both subgenomes (0.165 ± 0.842 in S and 0.310 ± 0.688 in D) (Table **1**).

Within *B. hybridum*, the three independent lineages showed marked differences in their genetic variation (Table **1**). The Ancient lineage exhibited significantly higher genetic diversity across all metrics when compared to the two Recent lineages. Specifically, its SNP heterozygosity was 0.0149 ± 0.0010 and its WG heterozygosity 0.0018 ± 0.0000, values substantially greater than those of the Recent West (0.0087 ± 0.0007; 0.0004 ± 0.0000) and Recent East (0.0081 ± 0.0005; 0.0004 ±0.0000) lineages, which showed similar or equal values (Table **1**). Consistent with this pattern, nucleotide diversity (π = 0.0039 ± 0.0015) was also highest in the Ancient lineage, whereas the Recent West and Recent East groups showed slightly lower values (π = 0.0037 ± 0.0020 and 0.0033 ± 0.0013, respectively). Tajima’s D values were positive in the Ancient (0.805 ± 0.182) and Recent West (0.506 ± 1.100) lineages, but strongly negative in the Recent East group (−1.470 ± 0.720). Furthermore, the Ancient lineage was characterized by a significantly lower inbreeding coefficient (Fis = 0.8162 ± 0.0079) and a correspondingly lower selfing rate (0.893 ± 0.0054) than both the Recent West (0.935 ± 0.0052; 0.967 ± 0.0028) and Recent East (0.939 ± 0.0038; 0.969 ± 0.0020) groups. Despite these differences in diversity and inbreeding levels, results from the Hardy-Weinberg equilibrium test indicate that all three lineages showed a significant deviation from HWE (p < 0.05) (Table **1**).

#### 3.1.4. Deleterious polymorphisms

We defined deleterious polymorphisms as the variants annotated as stop_gained, frameshift_variant, splice_acceptor_variant, splice_donor_variant, start_lost, stop_lost, and transcript_ablation / feature_ablation by SNPeff, and the nonsynonymous variants predicted as deleterious mutation by BAD_Mutation with potentially disruptive or functionally constrained effects on gene function. The combined analysis allowed us to distinguish between presumably deleterious variants widely distributed across genomic regions (according to SnpEff) and those with effects restricted to the protein level (according to BAD_Mutations).

Based on the functional annotation by SnpEff, most polymorphic variants were located outside coding regions, specifically in upstream gene regions (28.02–31.69%), downstream gene regions (26.83–28.53%), and intergenic regions (20.32–29.90%) (Table **S3A**). Intronic variants accounted for between 7.68% and 13.96%, while synonymous variants constituted only between 3.00% and 1.42% and missense mutations between 1.89% and 2.83%. Splice-site disruptions (acceptor, donor, region) and in UTRs were infrequent (<1% each), and very few loss-of-function alleles (stop codon gain, start codon loss) were detected (<0.05%).

To focus on polymorphic variants that potentially affect gene functions, we excluded intergenic and regulatory (upstream/downstream) variants and retained only sites in genes for subsequent analysis using BAD_Mutations. This likelihood-based framework identified which of these amino acid substitutions were significantly constrained (p < 0.05), representing truly deleterious changes.

Analysis of potentially deleterious polymorphic variants, normalized per megabase of coding sequence (Mb CDS), revealed contrasting patterns between the two subgenomes of *B. hybridum* and their diploid progenitors (Fig. **3**; Table **S3B**). In D-type genomes, the Ancient lineage showed the highest load (1,898.32 potentially deleterious variants / 797.11 confirmed deleterious per Mb CDS), followed by Recent West (803.03 / 328.75) and Recent East (736.43 / 271.37). The diploid progenitor *B. distachyon* exhibited the lowest values (598.87 / 235.18). In S-type genomes, the Ancient lineage again carried the highest load (2,635.30 / 1,048.58), far exceeding that of the Recent West (713.09 / 283.67) and Recent East (728.08 / 291.09) lineages. The diploid progenitor *B. stacei* showed the lowest S-genome load (624.86 / 248.95). Therefore, both subgenomes thus follow the same ranking: Ancient > Recent > diploid progenitor (Fig. **3**; Table **S3B**). At the genomic level, type D genomes exhibited a greater number of deleterious polymorphisms detected by SnpEff compared to type S genomes, except for the *B. hybridum* Ancient lineage, which showed considerably more mutations in its S subgenome than in its D subgenome, while the deleterious variants detected by BAD_Mutations were always higher in type S (sub)genomes than in type D (sub)genomes, and particularly in the *B. hybridum* Ancient lineage (Fig. **3**; Table **S3B**).

**Figure 3.**
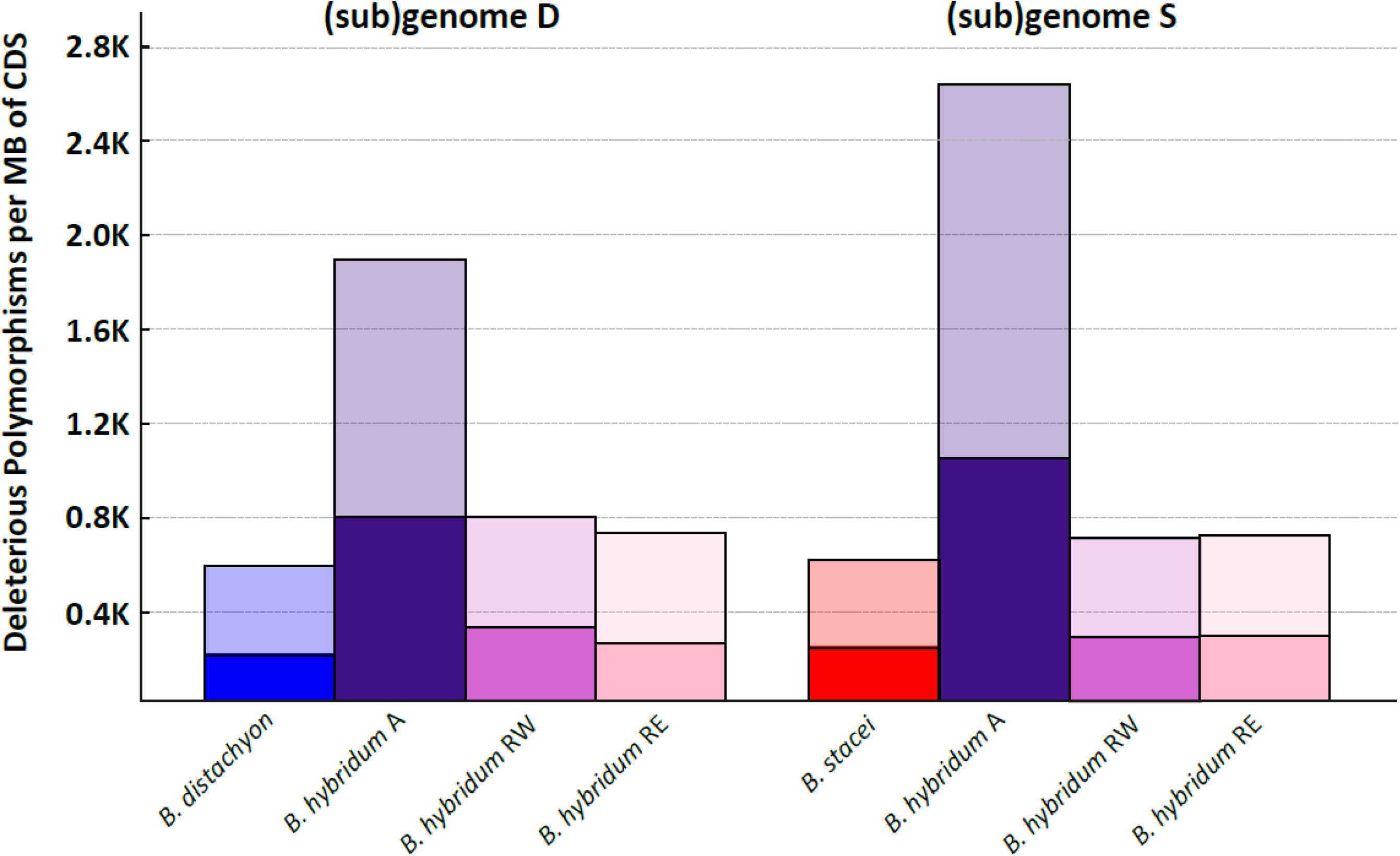
Genome-type level distribution of putatively deleterious polymorphic variants in Brachypodium hybridum allotetraploids and its diploid progenitor species. Colors denote species and B. hybridum lineages (B. distachyon: blue; B. stacei: red; B. hybridum Ancient: mauve; B. hybridum Recent West: wine; B. hybridum Recent East: light pink). Bars represent counts of potential deleterious load detected by SnpEff (with transparency) and those mutations inferred as deleterious (p < 0.05) from BAD_Mutations (solid color) per megabase (Mb) of CDS.

### 3.2. *Brachypodium hybridum* population genomics

#### 3.2.1. Nuclear genomic structure and cytonuclear discordances

Admixture analysis recovered an optimal K = 3 for the three *B. hybridum* lineages (Ancient, Recent West, and Recent East; Figs. **4A**, **S3**, Table **S4**) and quantified individual-level admixture proportions. Individuals from the Ancient group (n = 5) showed no detectable admixture (all Q > 0.99 to the Ancient cluster). In the Recent West group (n = 63), five accessions exhibited mixed ancestry: three from Greece, one from Israel, and one from Turkey. In the Recent East group (n = 61), 18 accessions were admixed, including 11 from Israel, three from Greece, and four from Turkey. These admixed individuals predominately occur in the eastern Mediterranean geographic contact zone between the two recently evolved *B. hybridum* groups where individuals from both lineages were found (Figs. **4A**, **S3**).

**Figure 4.**
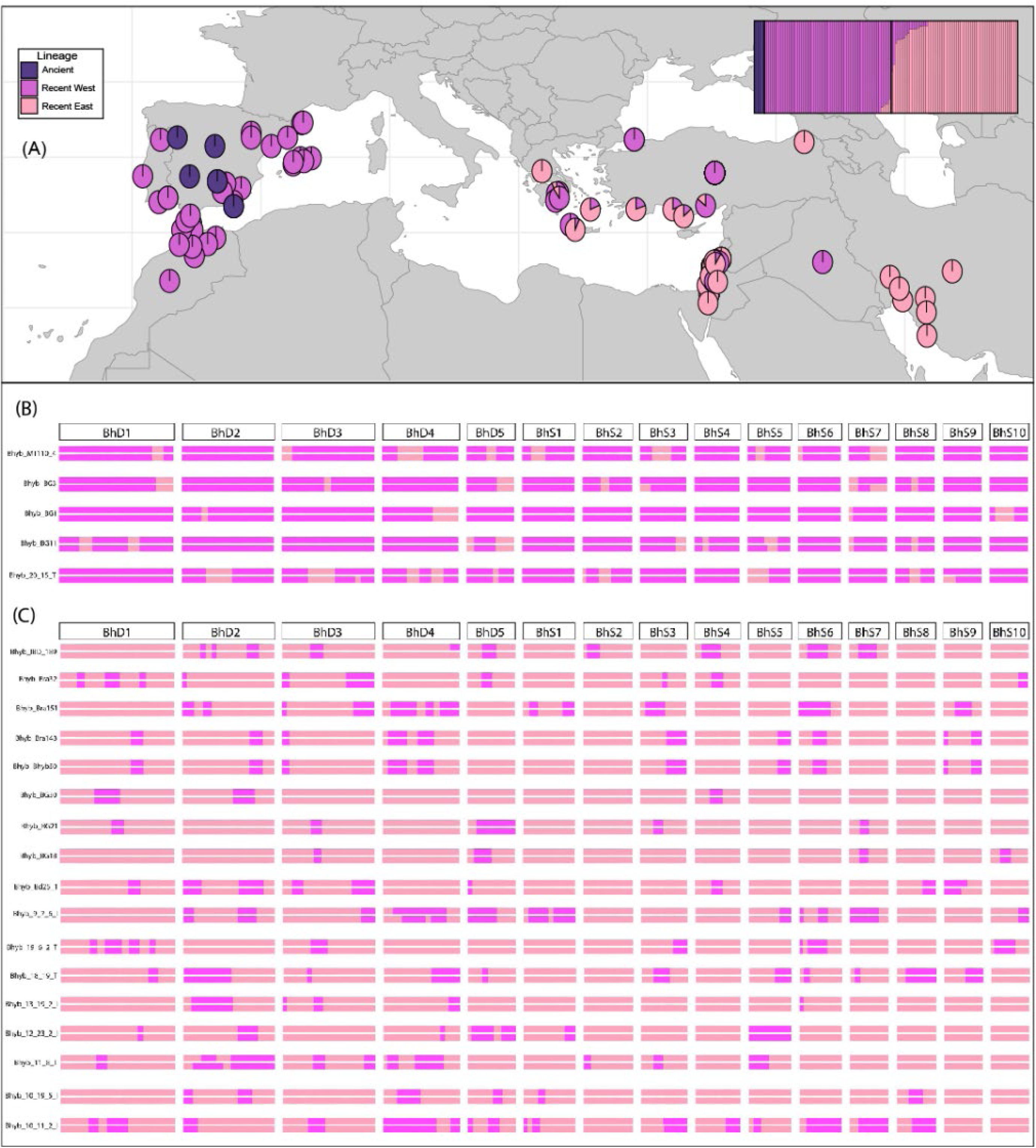
**(A)** Population structure of Brachypodium hybridum individuals inferred by Admixture (K = 3) and their circum-Mediterranean distribution. The individual-level barplot displays ancestry proportions for all 129 accessions, grouped by lineage (Ancient, Recent West, Recent East), with three colors corresponding to the genetic clusters identified. K = 3 was chosen as optimal based on the cross-validation error method (see Table **S4**). At each sampling locality, a pie chart shows the mean cluster membership for that individual. Slice angles are proportional to average ancestry proportions. Mixed plots correspond to 22 admixed accessions (see more details in Fig. **S3**). **(B, C)** Chromosomal-level admixture profiles of the 22 genomically admixed B. hybridum individuals identified by RF-mix. Panels show the mosaic ancestry composition along each of the 15 chromosome types for admixed accessions from the Recent West (n = 5) (B) and Recent East (n = 17) (C) lineages. Each horizontal track represents one individual, with chromosomes ordered from 1 to 15 left to right (BhD1-BhD5: B. distachyon type chromosomes; BhS1-BhS10: B. stacei-type chromosomes). Ancestry tracks were called using the three Admixture K = 3 reference clusters and were colored accordingly (Cluster 1 “Ancient-like”: mauve; Cluster 2 “Recent West–like”: wine; Cluster 3 “Recent East–like”: light pink). Note that in these recently admixed genomes only Cluster 2 (wine) and Cluster 3 (light pink) tracks are detected, whereas Cluster 1 (mauve) ancestry is absent.

Fine-scale local ancestry inference with RFmix v2.03 on the five admixed Recent West accessions confirmed admixture across all 15 chromosomes (Fig. **4B**). Rather than a chromosome-specific bias, each homologous pair displayed nearly identical ancestry proportions, reflecting highly homozygous introgressed tracks. In the Recent East group, 17 of the 18 accessions pre-identified as admixed retained detectable local ancestry segments (Fig. **4C**) while one individual did not show them. As in the Recent West cases, admixture patterns varied in length and position among chromosomes but were predominantly homozygous within each individual, indicating rapid fixation of foreign genomic markers across both subgenomes.

Contrasting topological analyses of the S and D subgenomic trees in *B. hybridum* recovered the same overarching structure, the Ancient lineage diverging earlier, followed by the split of the Recent West and Recent East lineages (Fig. **5**). Moreover, generalized Robinson–Foulds distance analysis yielded a gRF value of 0.4378 between the S and D subgenomic topologies, further highlighting relative topological congruence. Several Recent West and Recent East accessions shifted their relative positions within their respective clades in the S versus D trees, reflecting some internal population-level subgenomic restructuring (Fig. **5**).

**Figure 5.**
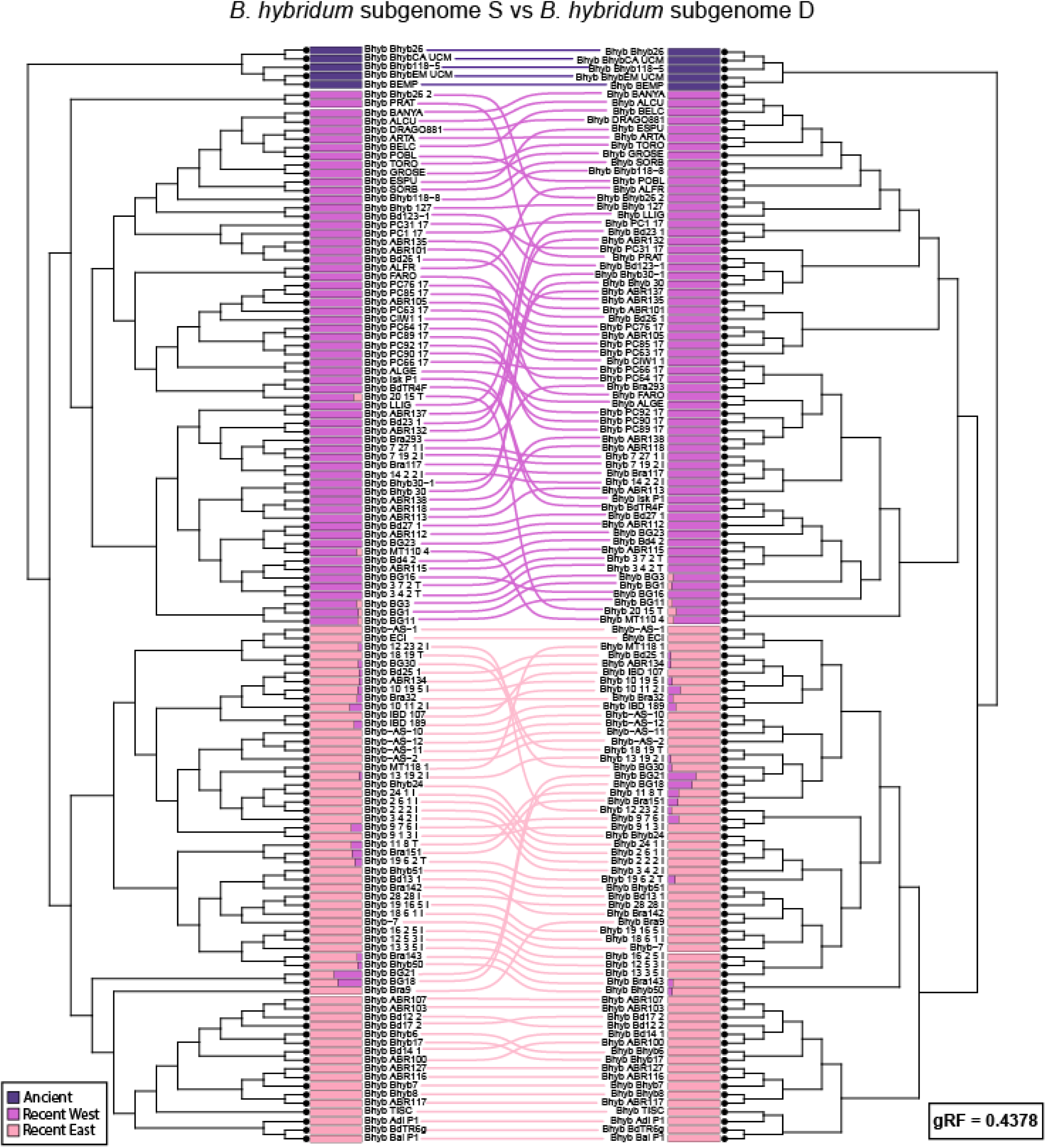
Cophylogenetic comparison of Brachypodium hybridum subgenome S- and subgenome D-based phylogenetic trees inferred by maximum-likelihood. Independent ML trees were reconstructed for each subgenome. Next to every tip on both the S- (left) and D-(right) subgenome trees, the corresponding Admixture K = 3 ancestry bar (as in Figs. **4A**, **S3**) is plotted, allowing direct comparison of nuclear admixture profiles across subgenomes. Colored lines between labels denote the three independent B. hybridum lineages (Ancient: mauve; Recent West: wine; Recent East: light pink). The generalized Robinson–Foulds (gRF) distance (0.4378) between the two topologies indicated relative congruence between both subgenomic trees.

Contrasting topological PACo analyses of nuclear versus plastome phylogenies yielded extensive topological discordance at both intra- and inter-clade levels for the recently diverged western and eastern Mediterranean *B. hybridum* lineages, while the Ancient lineage accessions showed no discordance (Figs. **6**, **S4**). Fifty-two accessions (40.3 %) showed significant residuals (upper two quartiles) indicating plastome–nuclear conflict; of them 27 corresponded to the Recent West clade (3 admixed) and 25 to the Recent East clade (all admixed). Complementary generalized Robinson–Foulds analysis returned a gRF distance of 0.8267 between the nuclear and plastome ML topologies, further underscoring the pronounced cytonuclear phylogenetic incongruence in these recently evolved lineages.

**Figure 6.**
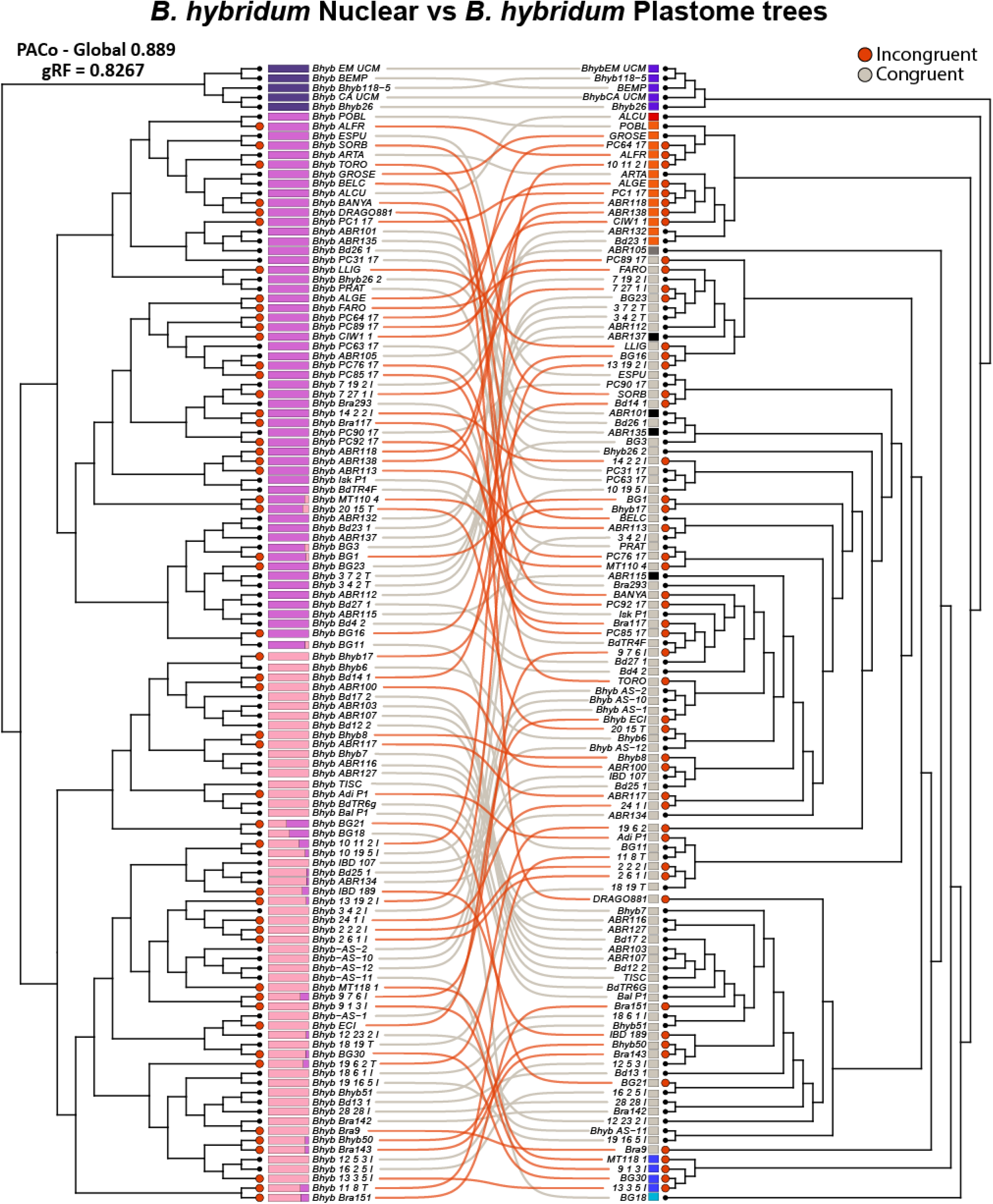
Comparison of Brachypodium hybridum nuclear versus plastome topological concordances based on Procrustean Approach to Cophylogeny (PACo) matching the nuclear ML tree (left) and the plastome ML tree (right). Tips showing congruent PACo residuals (lower two quartiles) are highlighted with gray line connectors, and those showing incongruent significant PACo residuals (upper two quartiles) are highlighted with orange circles and orange lines connectors. Bars next to the tips correspond to population groups representing the three independent lineages of B. hybridum according to nuclear SNPs Admixture analysis (left; see Figs. **4A**, **S3**) and plastome groups (right) based in Discriminant Analysis of Principal Components (DAPC) obtained from Campos et al. (2026). The PACo analysis detected cytonuclear concordance for the Ancient lineage, and intra- and inter-clade discordances for the Recent West and Recent East lineages.

#### 3.2.2. Genomic differentiation of *B. hybridum* lineages: genomic islands of divergence

Windowed Fst analyses revealed a pronounced gradient of differentiation across the *B. hybridum* genomes. In the Ancient vs. Recent lineages’ comparison mean Fst was 0.699 ± 0.179 (Fig. **S5A**), whereas Recent West vs. Recent East showed a substantially lower mean Fst of 0.312 ± 0.161 (Fig. **S5B**). Absolute divergence (Dxy) mirrored these trends (Fig. **S5C**, **S5D**); Ancient *vs* Recent lineages average Dxy was 0.019 ± 0.004 (Fig. **S5C**), while Recent West vs. Recent East comparison mean Dxy dropped to 0.006 ± 0.005 (Fig. **S5D**). Associations of significant outlier windows (95% cutoff and selection of common Fst and Dxy loci) with genomic regions revealed several overlapping windows of genomic islands of divergence, 199 islands for the Ancient - Recent comparison and 170 for Recent West - Recent East comparison. When we searched for genes within these islands, we detected 312 genes between Ancient and Recent lineages, and 288 genes between Recent West and Recent East groups.

The gene ontology (GO) enrichment analysis of the *B. hybridum* lineages revealed distinct sets of biological processes, molecular functions, and cellular components associated with each comparison (Table **2**). In the Ancient vs. Recent comparison, enriched biological processes included fructose metabolic process, regulation of mitosis, response to zinc ion, and brassinosteroid homeostasis, alongside broader categories like amino acid metabolic process and peptide modification. These genes were also associated with cellular components like retrotransposon nucleocapsid and cytoplasm-localized enzymes, suggesting metabolic reprogramming and stress-related responses in the Ancient lineage. Enriched molecular functions included carotenoid isomerase activity, enzyme regulator activity, oxidoreductase activity, iron-sulfur cluster binding, and heme binding, pointing to redox and regulatory adaptations linked to cellular metabolism. In the Recent West vs. Recent East comparison, enriched terms reflected divergence in developmental and physiological processes, including cytokinesis by cell plate formation, pollen germination, vernalisation response, and lignin biosynthetic process. Associated molecular functions were diverse, spanning DNA-binding transcription factor activity, oxidoreductase activity, enzyme regulator activity, and iron-sulfur cluster binding, among others. Notably, amino acid metabolic process was shared between the two comparisons, pointing to common functional pathways under selection across multiple lineages, together with four shared molecular functions: enzyme regulator activity, heme binding, iron-sulfur cluster binding, and oxidoreductase activity (Table **2**).

**Table 2.**
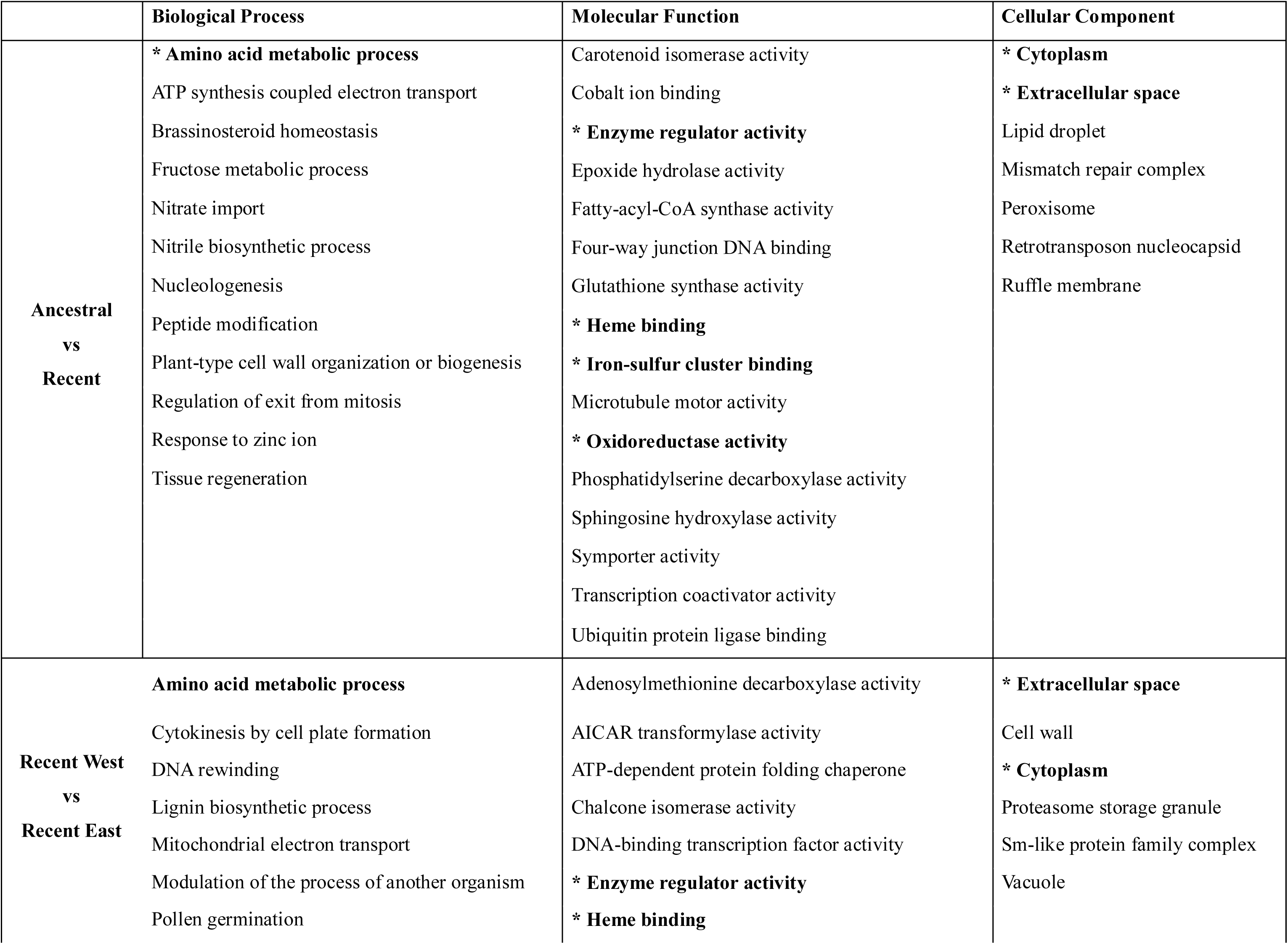
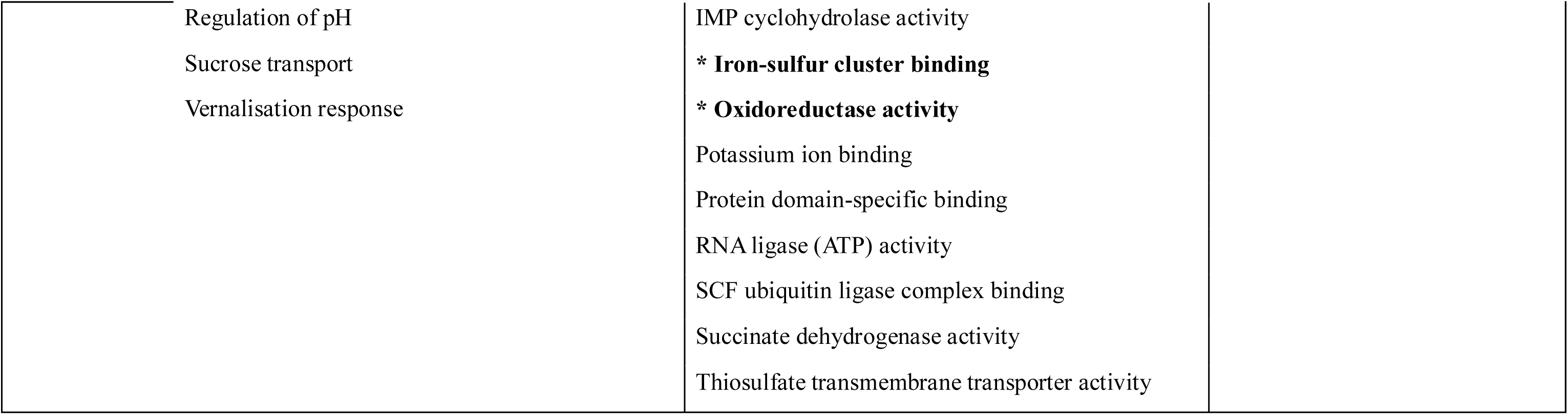
Summary of Gene Ontology (GO) terms categorized by biological process, molecular function, and cellular component for the comparison of genomic islands of divergence (Dxy, Fst) between the Ancient vs Recent lineages, and Recent West vs Recent East lineages of Brachypodium hybridum. The GO terms were obtained by first detecting outlier SNPs, identified as likely non-neutral based on Dxy and Fst estimates, and then mapping these SNPs to the annotated progenitor-reference genome. Functional annotations for the genes affected by these outliers SNPs were retrieved and associated with their corresponding GO terms. Shared GO terms between comparisons are highlighted in bold and marked with an asterisk.

|  | Biological Process | Molecular Function | Cellular Component |
| --- | --- | --- | --- |
| Ancestral<br>vs<br>Recent | * Amino acid metabolic process | Carotenoid isomerase activity | * Cytoplasm |
|  | ATP synthesis coupled electron transport | Cobalt ion binding | * Extracellular space |
|  | Brassinosteroid homeostasis | * Enzyme regulator activity | Lipid droplet |
|  | Fructose metabolic process | Epoxide hydrolase activity | Mismatch repair complex |
|  | Nitrate import | Fatty-acyl-CoA synthase activity | Peroxisome |
|  | Nitrile biosynthetic process | Four-way junction DNA binding | Retrotransposon nucleocapsid |
|  | Nucleologenesis | Glutathione synthase activity | Ruffle membrane |
|  | Peptide modification | * Heme binding |  |
|  | Plant-type cell wall organization or biogenesis | * Iron-sulfur cluster binding |  |
|  | Regulation of exit from mitosis | Microtubule motor activity |  |
|  | Response to zinc ion | * Oxidoreductase activity |  |
|  | Tissue regeneration | Phosphatidylserine decarboxylase activity |  |
|  |  | Sphingosine hydroxylase activity |  |
|  |  | Symporter activity |  |
|  |  | Transcription coactivator activity |  |
|  |  | Ubiquitin protein ligase binding |  |
| Recent West<br>vs<br>Recent East | Amino acid metabolic process | Adenosylmethionine decarboxylase activity | * Extracellular space |
|  | Cytokinesis by cell plate formation | AICAR transformylase activity | Cell wall |
|  | DNA rewinding | ATP-dependent protein folding chaperone | * Cytoplasm |
|  | Lignin biosynthetic process | Chalcone isomerase activity | Proteasome storage granule |
|  | Mitochondrial electron transport | DNA-binding transcription factor activity | Sm-like protein family complex |
|  | Modulation of the process of another organism | * Enzyme regulator activity | Vacuole |
|  | Pollen germination | * Heme binding |  |

|  |  |
| --- | --- |
| Regulation of pH | IMP cyclohydrolase activity |
| Sucrose transport | * **Iron-sulfur cluster binding** |
| Vernalisation response | * **Oxidoreductase activity** |
|  | Potassium ion binding |
|  | Protein domain-specific binding |
|  | RNA ligase (ATP) activity |
|  | SCF ubiquitin ligase complex binding |
|  | Succinate dehydrogenase activity |
|  | Thiosulfate transmembrane transporter activity |

The results of the PCoA analysis by genomic windows revealed a clear separation along PC1, which on average explains 4.25% of the genomic variance, between the Ancient and the Recently evolved *B. hybridum* lineages (Fig. **S6A**). The individuals from the ancient group exhibited a distinct clustering, indicating a strong genetic divergence from the recent groups. Among the Recent West and Recent East *B. hybridum* individuals, no significant differentiation was observed along PC1, suggesting a relatively uniform genetic pattern across these populations. However, PC2, accounting for 3.14% of variance, highlighted a slight separation between the two recent lineages of *B. hybridum* (Fig. **S6B**), distinguishing the Recent West and Recent East groups, suggesting a common genetic background with subtle differences corresponding to their geographic and evolutionary origins.

#### 3.2.3. Migration rates, Isolation by distance and environment

Pairwise genetic differentiation and inferred migration rates highlighted contrasting levels of gene flow among the three *B. hybridum* lineages (Table **3**). Wright’s Fst values were highest between the Ancient and both Recent West and Recent East lineages (0.61 and 0.65, respectively), reflecting strong divergence, whereas differentiation between Recent West and Recent East was more moderate (Fst = 0.34; Table **3****A**). Slatkin’s linearized Fst values reinforced these patterns, ranging from 0.52 (Recent West–Recent East) to 1.53 (Ancient–Recent West) and 1.87 (Ancient–Recent East) (Table **3****A**). Corresponding estimates of migrants per generation (Nm), were all below 1 for comparisons involving the Ancient lineage with the Recent East and Recent West groups (Nm = 0.327 and 0.268), indicating no effective gene flow, whereas Recent West and Recent East exchanged nearly one migrant per generation (Nm = 0.962), suggesting limited but appreciable connectivity (Table **3****B**).

**Table 3.** Pairwise genetic differentiation and inferred migration among Brachypodium hybridum lineages. (A) Genetic differentiation between the Ancient, Recent West, and Recent East lineages, measured by Wright’s Fst (range 0–1, where 0 denotes no differentiation and 1 indicates complete separation) and Slatkin’s linearized Fst/(1–Fst), which scales differentiation for comparison across loci. (B) Estimated number of migrants per generation (Nm) between each pair of lineages, derived from the equation Fst≈1/(4 Nm + 1) using Arlequin v3.5.2.2. Values of Nm < 1 suggest no gene flow, whereas Nm ≥ 1 indicates sufficient migration to counteract genetic drift.

| Fst / SlatkinFst | Ancient | Recent West | Recent East |
| --- | --- | --- | --- |
| **Ancient** | 0 | - | - |
| **Recent West** | 0.61 / 1.53 | 0 | - |
| **Recent East** | 0.65 / 1.87 | 0.34 / 0.52 | 0 |

(B)
| No. migrants (nm) | Ancient | Recent West | Recent East |
| --- | --- | --- | --- |
| **Ancient** | 1 | - | - |
| **Recent West** | 0.327 | 1 | - |
| **Recent East** | 0.268 | 0.962 | 1 |

The distance-based redundancy analysis of isolation by environment (IBE) and isolation by distance (IBD) models revealed significant effects of both environmental and geographic distances on genetic variation among populations (Table **4**). When tested independently at species level, environmental distance (IBE) explained 28.11 % of genetic variance, marginally more than geographic distance (IBD), which accounted for 26.17 %. After partially excluding the other predictor, IBE retained 13.58 % of variance (conditional IBE) and IBD retained 11.64 % (conditional IBD). Together, both factors explained 39.74 % of the total variance, with their shared contribution amounting to 14.53 %. At lineage level, the Ancient group yielded a saturated model but in both the Recent West and Recent East groups, IBE and IBD explain a larger proportion of genetic variance than at the species level, and their joint explanatory power is even greater. In the Ancient lineage, although the full model explained 100% of the genetic variance, neither IBE nor IBD were statistically significant when tested independently (p > 0.05), suggesting that neither predictor alone robustly explains the genetic variation in this group, likely due to the small sample size (n = 5) or the high collinearity between environmental and geographic distances, which had a shared contribution of 49%. In the Recent West lineage, IBE and IBD independently accounted for 30.11 % and 31.70 % of genetic variance, respectively; conditional effects were 23.10 % (IBE) and 24.69 % (IBD), and combined they explained 54.82 % (with 7.01 % overlap). Similarly, in the Recent East lineage, IBE explained 35.35 % and IBD 34.00 % independently; conditional variances were 31.20 % (IBE) and 29.85 % (IBD), and the full model accounted for 65.20 % (with 4.15 % overlap) (Table **4**).

**Table 4.** Summary of distance-based redundancy analysis (dbRDA) results assessing the effects of geographic isolation (IBD) and environmental isolation (IBE) distances on genomic differentiation within Brachypodium hybridum at species level (All) and at group level (Ancient, Recent West and Recent East lineages). Genetic, geographic, and environmental distance matrices were subjected to Principal Coordinates Analysis (PCoA), and dbRDA models were fitted using the resulting axes as predictors. Separate analyses were conducted for the Ancestral, Recent West and Recent East groups. P-values were obtained from permutation tests (n = 10,000). The marginal variance represents the percentage of genetic variation explained by each predictor when tested independently. The conditional variance corresponds to the variation explained by a given predictor after accounting for the effect of the other (i.e., partial dbRDA). The shared variance represents the proportion of variance jointly explained by both geography and environment in the full model. Asterisks indicate significant p-values (p < 0.05), suggesting that the corresponding predictor explains a significant portion of the genetic structure.

|  | Model | % Expl. Var | % Expl. Var. Conditional | % Overlapping Var. |
| --- | --- | --- | --- | --- |
| All | IBE | 28.11%* | 13.58%* | - |
|  | IBD | 26.17%* | 11.64%* | - |
|  | Combined | 39.74%* | - | 14.53% |
| Ancient | IBE | 75.59% | 26.59% | - |
|  | IBD | 73.42% | 24.41% | - |
|  | Combined | 100% | - | 48.99% |
| Recent West | IBE | 30.11%* | 23.10%* | - |
|  | IBD | 31.70%* | 24.69%* | - |
|  | Combined | 54.82%* | - | 7.01% |
| Recent East | IBE | 35.35%* | 31.20%* | - |
|  | **IBD** | 34.00%* | 29.85%* | - |
|  | **Combined** | 65.20%* | - | 4.15% |

## 4. Discussion

### 4.1. Multiple origins of *B. hybridum* and their diverse population genomics

This study demonstrates that a wild allopolyploid can simultaneously exhibit recurrent origins spanning nearly two million years, long-term nuclear subgenomic stability, extensive cytonuclear discordance, and contrasting trajectories of deleterious-variant accumulation; all within a circum-Mediterranean geographic framework. These features are rarely captured together in a single natural system, making *B. hybridum* a powerful model for testing how the fundamental processes of polyploid formation, genome stabilization, mutation-load dynamics, and environmental filtering interact over evolutionary time.

Our population genomic analysis of 307 accessions confirms three independent allopolyploid origins for *B. hybridum*, an Ancient lineage and two more recent lineages (Recent West and Recent East) (Gordon *et al*., 2020; Scarlett *et al*., 2023; Mu *et al*., 2023a), and significantly expands their known geographic distributions (Figs. **1****, 2**). The penalized likelihood divergence times estimated for the crown nodes of these three lineages (Ancient, ∼1.78 Ma; Recent West, ∼0.56 Ma; Recent East, ∼0.24 Ma; Fig. **S2**) are consistent with previous cross-bracing estimations (Gordon et al., 2020; Mu et al., 2023a), providing a robust temporal framework for investigating the evolutionary trajectory of each lineage. Each independent allopolyploidization event has left a distinct genomic footprint shaped by its respective progenitor lineages, geographic origin, and demographic history.

A key finding of our study is the juxtaposition of a highly stable and three-times independently originated nuclear genome with a remarkably discordant plastome history. Comparison of the S and D nuclear subgenomic phylogenies within *B. hybridum* revealed high topological congruence (generalized Robinson–Foulds distance of 0.4378; Fig. **5**), indicating a stable, shared phylogenetic backbone in which the Ancient lineage diverged (or was formed) first, followed by the more closely related Recent West and Recent East lineages. This nuclear consistency agrees with previous comparative genomic analyses of *B. hybridum* reporting extremely rare homeologous exchange events between subgenomes in all three independent lineages (Mu et al. 2023a). Even the evolutionary close Recent West and Recent East lineages show low admixture and limited chromosomal introgression in both subgenomes (Figs. **4**, **S3**), reinforcing their independent evolutionary trajectories. This pattern of stability and post-polyploidization independence, which is also found in some allopolyploids such as *Trifolium repens* (Griffiths et al., 2019) and *Arabidopsis suecica* (Burns *et al*., 2021), contrasts with more dynamic allopolyploid systems in which subgenome interactions are accompanied by stronger expression asymmetries, biased fractionation, or extensive genomic restructuring, such as those observed in *Tragopogon miscellus* (Chester et al., 2012), *Gossypium* allotetraploids (Flagell & Wendel, 2010; Guo et al., 2014), *Brassica napus* (Hurgobin *et al*., 2018), and *Fragaria x ananassa* (Edger *et al*., 2019). In maize, for example, population genomic analyses have revealed pervasive signatures of local adaptation and introgression shaping genomic diversity at large geographic scales (Hufford *et al*., 2021). Similarly, in bread wheat, the allopolyploid genome has been strongly influenced by introgression from wild relatives such as *Aegilops* species, resulting in a more dynamic and reticulate evolutionary history (Zhou *et al*., 2021; Avni *et al*., 2022). The stability of *B. hybridum* suggests that selfing and the absence of backcrossing to diploid progenitors can lock in the initial genomic configuration of an allopolyploid and preserve it over millions of years.

Our analyses revealed profound discordance between the nuclear and plastome phylogenies of *B. hybridum*, particularly in the recent lineages, which share an S-type plastome, but not in the Ancient lineage, whose samples show a D-type plastome (Table **1**; Gordon *et al*., 2020; Mu *et al*., 2023a). PACo analysis identified significant topological conflict in ∼40% of accessions, and quantified a high gRF distance of 0.8267 (Figs. **6**, **S4**). This pattern is likely attributable to recurrent chloroplast capture between the Recent West and Recent East lineages (Figs. **6**, **S4**), a phenomenon previously documented in the progenitor species *B. distachyon* and *B. stacei* (Campos et al., 2024, 2026; Sancho et al., 2018). These findings demonstrate that differential evolutionary pressures and introgressive hybridization can profoundly shape phylogenomic patterns even in highly selfing systems, leading to complex cytonuclear relationships (Toews & Brelsford, 2012; Sloan *et al*., 2017). Cytonuclear incompatibilities have been documented in plant hybrids and nascent allopolyploids, where mismatches between divergent nuclear and organellar genomes can reduce fertility or viability (Burke & Arnold, 2001; Fishman & Willis, 2006; Postel & Touzet, 2020). However, evidence from *Brassica* (Ferreira de Carvalho *et al*., 2019), synthesis studies across diverse angiosperm lineages (Sloan *et al*., 2024), and a synthetic allotetraploid *Brachypodium hybridum* (Dinh Thi *et al*., 2016), indicates that many polyploid plants buffer these perturbations, attaining reproductive stability and speciation. While nuclear subgenomic histories remain well-aligned at deep nodes, the profound plastome–nuclear discordance in the Recent West and Recent East lineages reveals that chloroplast capture continues to reshape cytoplasmatic evolutionary trajectories long after polyploid formation.

### 4.2. Subgenomic divergence and deleterious variant load

Our analyses reveal contrasting trajectories of deleterious variant accumulation between the diploid progenitors’ genomes and the corresponding *B. hybridum* subgenomes (Fig. **3**, Table **S3**). The *B. hybridum* subgenomes exhibit a higher deleterious variant burden than their respective progenitor genomes. This could reflect duplicated genetic content in allotetraploids, although expression studies have shown that most S and D genes are co-expressed in *B. hybridum* and subgenomic dominance is very rare (Scarlett *et al*., 2023; Mu *et al*., 2023a). In the D-type and S-type genomes, the Ancient lineage shows the highest number of deleterious variants, followed by the Recent West and Recent East lineages, while the diploid progenitor species *B. stacei* and *B. distachyon* exhibit the lowest (Table **S3B**). These results support an initial increase in mildly deleterious variants linked to older hybridization, followed by subsequent elimination and/or reduced influx in more recent origins. However, the pattern may also reflect different burdens of deleterious variants in the parental genomes that contributed to each allotetraploid lineage, which are unknown for the ancient lineage of *B. hybridum* (Catalán *et al*., 2016a; Mu *et al*., 2023a).

From a mechanistic perspective, this is consistent with strong genetic drift and a reduction in the effectiveness of purifying selection, mediated by self-fertilization, during or just after the formation of the earliest allopolyploid (Wright *et al*., 2013), which increases the initial genetic load, and with more effective purifying selection or larger effective sizes in later-origin lineages that limited or eliminated some of that load (Paape *et al*., 2018; Kryvokhyzha *et al*., 2019). Subgenomic scans reveal that both subgenomes follow the same ranking, yet the S subgenome of the Ancient lineage carries a particularly heavy burden (1,048.58 confirmed deleterious variants/Mb CDS, versus 797.11 for the D subgenome), which may reflect a high legacy load inherited from its extinct S-progenitor. Subtle lineage differences persist between recent lineages; in subgenome D, the recent western lineage predominates over the eastern one, while in subgenome S the two recent lineages are nearly equal. This likely reflects distinct demographic histories and balances between selection and genetic drift, rather than intrinsic properties of the subgenomes (Conover & Wendel, 2022). Alternatively, it is possible that the ancient population is the only remaining group of a lineage prone to the accumulation of deleterious mutations, given that its presumed diploid progenitors likely became extinct (Mu *et al*., 2023a).

The disomic inheritance of *B. hybridum* has likely restricted homeologous exchanges, preserving subgenome identity and allowing for largely independent mutation accumulation. Homeologous exchange events are rare in all three lineages (Mu *et al*., 2023a), indicating effective amphidiploidy with minimal intersubgenomic recombination. This genomic stability, along with the absence of the strong subgenome dominance frequently reported in other systems, fits a relatively symmetric polyploid model (Bird *et al*., 2018), but also demonstrates that expression symmetry does not preclude asymmetry in mutational burden (Mason & Wendel, 2020).

### 4.3. Population structure, admixture, and gene flow

Our population genomic analyses have revealed a well-defined genetic structure within *B. hybridum*, shaped by its multiple independent origins and a predominantly selfing mating system that restricts gene flow. Admixture analyses robustly identify three major genetic lineages: Ancient, Recent West, and Recent East (Fig. **4A**); that align with the three independent polyploidization events confirmed by our phylogenomic reconstructions (Figs. **2**, **S1**, **S2**) and previous studies (Gordon *et al*., 2020; Scarlett *et al*., 2023; Mu *et al*., 2023a). The strong genomic differentiation observed among these lineages (Fst ≥ 0.61; Table **3**) matches their clear temporal and geographic separation, reinforcing the view that recurrent formation is a key mechanism for generating diversity in polyploids, creating distinct gene pools from the outset (Soltis & Soltis, 2000; Te Beest *et al*., 2012).

The genetic diversity patterns differ markedly among these lineages, reflecting their distinct demographic and evolutionary trajectories (Table **1**). The Ancient lineage displays the highest genetic diversity (SNP Heterozygosity = 0.0149), a value significantly greater than those of the Recent West and Recent East lineages. This greater diversity, observed despite a smaller sample size, is likely a "legacy" of high variation inherited from its unknown diploid progenitors (Mu *et al*., 2023a). This aligns with the concept that subgenomes are not evolutionary blank slates but carry the indelible imprint of their parental history (Wendel *et al*., 2016). In contrast, the two Recent lineages exhibit significantly lower diversity and higher inbreeding (Fis ≈ 0.94), a pattern consistent with more recent origins potentially shaped by founder effects during colonization (Te Beest *et al*., 2012).

Complementary analyses based on nucleotide diversity (π) and Tajima’s D supported these patterns (Table **1**). π differed significantly among species, being highest in *B. distachyon*, intermediate in *B. hybridum*, and lowest in *B. stacei*, consistent with contrasts in long-term effective population size (Akhunov *et al*., 2010). Within *B. hybridum*, the S subgenome resembled *B. stacei* and the D subgenome resembled *B. distachyon*, both showing slightly reduced π values relative to their progenitors, suggesting partial erosion of ancestral polymorphism after polyploid formation (Page *et al*., 2013). Tajima’s D was negative in both diploids but positive in *B. hybridum*, indicating an excess of intermediate-frequency variants that may result from population structure or demographic stability (Carlson *et al*., 2005; Korneliussen *et al*., 2013). Across *B. hybridum* lineages, π showed similar values, with the Ancient lineage slightly higher than the Recent West and Recent East groups, whereas Tajima’s D was most positive in the Ancient lineage and strongly negative in the Recent East. This pattern is consistent with a scenario of long-term demographic stability or balancing selection in the Ancient lineage (positive D), and a recent population expansion or selective sweep in the Recent East lineage (negative D), while the negative D in both diploid progenitors probably reflect their historical demographic expansions (Vlček *et al*., 2025).

Despite high rates of selfing, our analyses reveal subtle secondary contact between the two recent lineages. The Ancient lineage remains effectively isolated, with no admixture detected (Figs. **4A**, **S3**) and negligible estimated gene flow (Nm < 0.33; Table **3**), suggesting that strong reproductive barriers have been maintained over its nearly two-million-year divergence from the other allotetraploids (Fig. **S2**). This agrees with the failure to cross recent and ancient lines in the laboratory (Gordon *et al*., 2020; Scarlett *et al*., 2023). In contrast, 23 admixed individuals were identified in the geographic contact zone between the Recent West and Recent East lineages (Fig. **4A**), with fine-scale analysis confirming mosaic ancestry along their chromosomes (Figs. **4B**, **4C**). Chromosomal-level ancestry resolution has been successfully employed in other complex plant systems, such as *Arabidopsis suecica* (Novikova *et al*., 2017) and bread wheat relatives (Zhou *et al*., 2021), where ’ancestry painting’ has allowed the detection of specific introgressed blocks that persist despite high selfing rates. The estimated gene flow (Nm ≈ 0.96; Table **3**) suggests that while rare outcrossing occurs, the dominant selfing mating system largely preserves the genetic integrity of each lineage, a common feature in colonizing species (Stebbins, 1985; Wright *et al*., 2013). Thus, the Recent West and Recent East lineages exhibit limited but detectable admixture restricted to their geographic contact zone, while the Ancient lineage remains effectively isolated.

At subgenomic level within *B. hybridum*, both subgenomes have experienced a loss of genetic diversity following polyploidization, consistent with founder effects or bottlenecks during allopolyploid formation (Mu *et al*., 2023a). The asymmetric evolution of subgenomes is a hallmark of polyploidy and can be driven by the distinct histories of the progenitor genomes (Wendel *et al*., 2016). *B. hybridum* provides an interesting contrast to that of bread wheat, where the more recently incorporated D subgenome is the least diverse (Pont *et al*., 2019), highlighting that subgenome evolutionary trajectories are system-specific. Given the selfing mating system of *B. hybridum*, the reduced effective recombination rate likely amplifies the footprint of linked selection, extending over larger genomic tracks and accentuating Fst peaks even without strong barriers to introgression (Nordborg *et al*., 1996; Hartfield & Glémin, 2016). This mechanism could underlie the concentration of divergence islands in particular genomic regions (Figs. **S5A**, **S5B**), mirroring observations in *B. distachyon*, where genomic outliers of differentiation have been linked to the polygenic architecture of adaptive traits such as flowering time (Minadakis *et al*., 2024) and are often associated with the non-random accumulation of transposable elements (TEs) in low-recombination regions (Stritt *et al*., 2019). The overlap of these ’islands’ across species suggests that certain genomic regions are inherently prone to accumulate divergence due to the interplay between structural variation and selection on life-history traits.

Overall, the genetic structure of *B. hybridum* illustrates how recurrent, independent origins can generate substantial diversity in a wild polyploid, even in the absence of significant subsequent gene flow. Unlike many crops where post-domestication introgression with diploid relatives is a key source of diversity, *B. hybridum* serves as a model for diversification driven primarily by its multiple formations and the absence of backcrossing to any of its progenitor species. The lack of backcrossing with its diploid progenitors is consistent with the strong reproductive barriers typically imposed by ploidy mismatches in wild allopolyploids (Ramsey & Schemske, 1998), underscoring the importance of polyploidy as a potent speciation mechanism that can lead to rapid reproductive isolation and independent evolutionary pathways (Abbott *et al*., 2013).

### 4.4. Functional categories under selection

The enrichment of functional categories within genomic islands (windows above the 95th percentile of Fst and/or Dxy; see Methods) provides additional insight into the evolutionary trajectories of *B. hybridum* lineages (Table **2**). In the Ancient vs. Recent comparison, processes related to brassinosteroid homeostasis, mitotic regulation and stress responses were prominent, whereas in the Recent West vs. East contrast, developmental and phenological pathways such as vernalization, pollen germination and lignin biosynthesis emerged as significant (Table **2**). These categories match ecological pressures expected in the circum-Mediterranean distribution, including adaptation to drought, ion imbalance and seasonal shifts. Brassinosteroid pathways are well known to mediate drought and ion-stress tolerance (Wei & Li, 2020; Gillani *et al*., 2022), flowering time regulation via vernalization genes has been extensively documented in *Brachypodium* (Colton-Gagnon *et al*., 2014; Ream *et al*., 2014; Woods *et al*., 2017), and lignin deposition is a recurrent mechanism of drought resilience in grasses (Moura et al., 2010; Song et al., 2022). The identification of these functional categories within genomic islands in *B. hybridum* mirrors the patterns observed in its progenitor *B. distachyon*, where outliers of differentiation are strongly associated with the polygenic architecture of flowering time and local adaptation to Mediterranean environmental gradients (Minadakis *et al*., 2024). The emergence of these islands may also be influenced by the ancestral genomic landscape; in *B. distachyon*, regions of high differentiation often overlap with low-recombination areas enriched in transposable elements (TEs), which can accentuate Fst peaks and preserve adaptive haplotypes (Stritt *et al*., 2019). Although enrichment does not prove causality, these pathways constitute promising candidates underlying lineage-specific adaptation.

### 4.5. Isolation by environment versus distance

Isolation-by-environment (IBE) emerged as a slightly stronger determinant of genetic differentiation than isolation-by-distance (IBD) across the *B. hybridum* range, indicating that ecological factors play an important role in shaping lineage divergence (Table **4**). At the species level, environmental distances accounted for 28.1% of genetic variance, marginally exceeding geographic distances (26.2%) and retained 13.6% of variance after conditioning on geography. This pattern intensified within the Recent West and Recent East clades: in Recent West, IBE and IBD independently explained ∼30.1% and ∼31.7% of variance, respectively, but conditional IBD (24.7%) remained slightly higher than conditional IBE (23.1%), and together they captured 54.8% of variance (with only 7.0% overlap). In Recent East, IBE (35.3%) and IBD (34.0%) were both strong drivers, yet conditional IBE (31.2%) marginally outpaced conditional IBD (29.9%), yielding a combined 65.2% of explained variance (4.2% shared). These results contrast with classic IBD-driven scenarios in selfing annual grasses where limited dispersal often makes geographic distance the dominant force (Bakker *et al*., 2009; Jenkins *et al*., 2010), but align with growing evidence from Mediterranean *Brachypodium* species showing that soil and microclimate differences generate significant genetic structure even when populations are geographically close (Marques *et al*., 2017; Shiposha *et al*., 2020; Campos *et al*., 2024). In *B. distachyon*, Marques et al. (2017) demonstrated that limestone versus acidic-soil populations in Iberian Peninsula diverged genetically despite close proximity. In *B. stacei*, Mu *et al*. (2023b) documented fine-scale IBE at “Evolution Canyon”, where opposing slopes support genetically distinct populations adapted to mesic versus xeric microclimates. By analogy, *B. hybridum*, an allotetraploid inhabiting intermediate habitat, likely experiences similar environmental filtering. Our partial dbRDA shows that environmental axes explain additional genetic variance after removing geographic effects, implying that selection on habitat-specific traits contributes to within-lineage divergence. These results collectively demonstrate that environmental differences frequently outweigh geographic distance in driving genetic differentiation within *B. hybridum*, underscoring the central role of local adaptation in shaping diversity across its circum-Mediterranean range.

By showing how recurrent polyploidization, cytonuclear reshuffling, subgenomic divergence, and environmental filtering combine to generate and maintain genetic variation, *B. hybridum* stands as a powerful model for unraveling the mechanisms of polyploid evolution and adaptation in wild grasses. The features documented here (independent origins spanning nearly two million years, long-term nuclear genome stability despite ongoing chloroplast capture, and contrasting subgenomic trajectories of deleterious-variant accumulation) are rarely captured together in a single study. This combination makes *B. hybridum* uniquely suited for testing how the fundamental processes of allopolyploid evolution unfold in natural populations, offering lessons that extend well beyond the *Brachypodium* genus to polyploid plant systems worldwide.

## Data Availability

Data generated for this project are available on Zenodo under the DOI 10.5281/zenodo.22042302. Custom scripts designed for the analyses performed in this study are available on Github (https://github.com/Bioflora/BhybridumPopGenomics).

## Author Contributions

PC, JV and EP obtained funding for the project. PC, MC and RS designed the study. MC, RS, AGW, LL and PC conducted and supervised the analyses. BCM and RS adapted the AlloSHP code to this study. MC and PC wrote the original draft, and all authors reviewed it.

## Acknowledgments

The authors wish to thank the staff at the DOE Joint Genome Institute (JGI) for whole-genome sequencing support, and all collaborators and institutions that contributed plant material and logistical assistance for this study.

## Figures & Tables

**Figure 7.** Circum-Mediterranean geographic distribution of population samples of the three annual *Brachypodium* species under study **(A)**: *B. distachyon* (n = 113; blue), *B. stacei* (n = 65; red) and *B. hybridum* (n = 125; pink). **(B)** Distribution of the three independent *B. hybridum* lineages: Ancient (n = 5; mauve), Recent West (n = 59; wine) and Recent East (n = 61; light pink). (see Table **S1**). Of the total of 129 individuals of *B. hybridum*, four individuals from non-native areas (1 Australia, 2 South Africa, 1 Uruguay; see Table **S1**) were excluded for this figure, all four belonging to the Recent West group (n=63) (see Results).

**Figure 8.** Multispecies coalescent tree of the *Brachypodium* annual complex samples inferred with *Weighted-*ASTRAL. Maximum-likelihood trees were estimated for each non-overlapping 10,000 bp windows of the full syntenic SNP matrix (∼8 million SNPs), and the resulting trees were combined under the ASTER package v.1-3.5 ASTRAL (Zhang et al. 2025) to reconstruct the MSC tree. Tips represent individual sequences: *B. distachyon* (n = 113), *B. stacei* (n = 65), and for *B. hybridum* two subgenomic sequences per accession (D and S subgenomes; n = 129 accessions). Branch colors denote genome/species/subgenome lineage identity and tip-colored squares correspond to geographic regions as indicated in the respective charts. Support values of nodal quartets are indicated in Fig. **S1A**.

**Figure 9.** Genome-type level distribution of putatively deleterious polymorphic variants in *Brachypodium hybridum* allotetraploids and its diploid progenitor species. Colors denote species and *B. hybridum* lineages (*B. distachyon*: blue; *B. stacei*: red; *B. hybridum* Ancient: mauve; *B. hybridum* Recent West: wine; *B. hybridum* Recent East: light pink). Bars represent counts of potential deleterious load detected by SnpEff (with transparency) and those mutations inferred as deleterious (p < 0.05) from BAD_Mutations (solid color) per megabase (Mb).

**Figure 10.** (A) Population structure of *Brachypodium hybridum* individuals inferred by Admixture (K = 3) and their circum-Mediterranean distribution. The individual-level barplot displays ancestry proportions for all 129 accessions, grouped by lineage (Ancient, Recent West, Recent East), with three colors corresponding to the genetic clusters identified. K = 3 was chosen as optimal based on the cross-validation error method (see Table **S4**). At each sampling locality, a pie chart shows the mean cluster membership for that individual. Slice angles are proportional to average ancestry proportions. Mixed plots correspond to 22 admixed accessions (see more details in Fig. **S3**). **(B, C)** Chromosomal-level admixture profiles of the 22 genomically admixed *B. hybridum* individuals identified by RF-mix. Panels show the mosaic ancestry composition along each of the 15 chromosome types for admixed accessions from the Recent West (n = 5) (B) and Recent East (n = 17) (C) lineages. Each horizontal track represents one individual, with chromosomes ordered from 1 to 15 left to right (BhD1-BhD5: *B. distachyon* type chromosomes; BhS1-Bh10: *B. stacei*-type chromosomes). Ancestry tracks were called using the three Admixture K = 3 reference clusters and were colored accordingly (Cluster 1 “Ancient-like”: mauve; Cluster 2 “Recent West–like”: wine; Cluster 3 “Recent East–like”: light pink). Note that in these recently admixed genomes only Cluster 2 (wine) and Cluster 3 (light pink) tracks are detected, whereas Cluster 1 (mauve) ancestry is absent.

**Figure 11.** Cophylogenetic comparison of *Brachypodium hybridum* subgenome S and subgenome D-based phylogenetic trees inferred by maximum-likelihood. Independent ML trees were reconstructed for each subgenome. Next to every tip on both the S- (left) and D-(right) subgenome trees, the corresponding Admixture K = 3 ancestry bar (as in Figs. **4A**, **S3**) is plotted, allowing direct comparison of nuclear admixture profiles across subgenomes. Colored lines between labels denote the three independent *B. hybridum* lineages (Ancient: mauve; Recent West: wine; Recent East: light pink). The generalized Robinson–Foulds (gRF) distance (0.4378) between the two topologies indicated relative congruence between both subgenomic trees.

**Figure 12.** Comparison of *Brachypodium hybridum* nuclear versus plastome topological concordances based on Procrustean Approach to Cophylogeny (PACo) matching the nuclear ML tree (left) and the plastome ML tree (right). Tips showing congruent PACo residuals (lower two quartiles) are highlighted with gray line connectors, and those showing incongruent significant PACo residuals (upper two quartiles) are highlighted with orange circles and orange lines connectors. Bars next to the tips correspond to population groups representing the three independent lineages of *B. hybridum* according to nuclear SNPs Admixture analysis (left; see Figs. **4A**, **S3**) and plastome groups (right) based in Discriminant Analysis of Principal Components (DAPC) obtained from Campos et al. (2026). The PACo analysis detected cytonuclear concordance for the Ancient lineage, and intra- and inter-clade discordances for the Recent West and Recent East lineages.

**Table 5.** Genomic diversity parameters (heterozygosity, nucleotide diversity (π), Tajima’s D, inbreeding coefficient (Fis), deviations from Hardy-Weinberg equilibrium (HWE), and selfing rate) of the *Brachypodium distachyon*-*B. stacei*-*B. hybridum* (Bd-Bs-Bh) complex at the species level, and separately for each subgenome (subgenome S, subgenome D) within *B. hybridum* and the three independent lineages of *B. hybridum* (Ancient, Recent West, Recent East). P-values from Hardy-Weinberg equilibrium tests are provided to assess deviations from equilibrium in each group and subgenome. Post-hoc Wilcoxon pairwise test letter assignments are provided for each index, indicating significant differences with p-values < 0.05. All pairwise comparisons for π and Tajima’s D were statistically significant; however, the assigned letters were determined according to both statistical significance and effect size (r for pairwise contrasts). Comparisons were made between the three species, between the progenitor species and the two subgenomes of *B. hybridum*, and between the three lineages of *B. hybridum*.

**Table 6.** Summary of Gene Ontology (GO) terms categorized by biological process, molecular function, and cellular component for the comparison of genomic islands of divergence (Dxy, Fst) between the Ancient vs recent lineages, and Recent West vs Recent East lineages of *Brachypodium hybridum*. The GO terms were obtained by first detecting outlier SNPs, identified as likely non-neutral based on Dxy and Fst, and then mapping these SNPs to the annotated progenitor-reference genome. Functional annotations for the genes affected by these outliers SNPs were retrieved and associated with their corresponding GO terms. Shared GO terms between comparisons are highlighted in bold and marked with an asterisk.

**Table 7.** Pairwise genetic differentiation and inferred migration among *Brachypodium hybridum* lineages. (A) Genetic differentiation between the Ancient, Recent West, and Recent East lineages, measured by Wright’s Fst (range 0–1, where 0 denotes no differentiation and 1 indicates complete separation) and Slatkin’s linearized Fst/(1–Fst), which scales differentiation for comparison across loci. (B) Estimated number of migrants per generation (Nm) between each pair of lineages, derived from the equation Fst≈1/(4 Nm + 1) using Arlequin v3.5.2.2. Values of Nm < 1 suggest no gene flow, whereas Nm ≥ 1 indicates sufficient migration to counteract genetic drift.

**Table 8.** Summary of distance-based redundancy analysis (dbRDA) results assessing the effects of geographic isolation (IBD) and environmental isolation (IBE) distances on genomic differentiation within *Brachypodium hybridum* at species level (All) and at group level (Ancient, Recent West and Recent East lineages). Genetic, geographic, and environmental distance matrices were subjected to Principal Coordinates Analysis (PCoA), and dbRDA models were fitted using the resulting axes as predictors. Separate analyses were conducted for the Ancient, Recent West and Recent East lineages. P-values were obtained from permutation tests (n = 10,000). The marginal variance represents the percentage of genetic variation explained by each predictor when tested independently. The conditional variance corresponds to the variation explained by a given predictor after accounting for the effect of the other (i.e., partial dbRDA). The shared variance represents the proportion of variance jointly explained by both geography and environment in the full model. Asterisks indicate significant p-values (p < 0.05), suggesting that the corresponding predictor explains a significant portion of the genetic structure.

**Figure S1.**
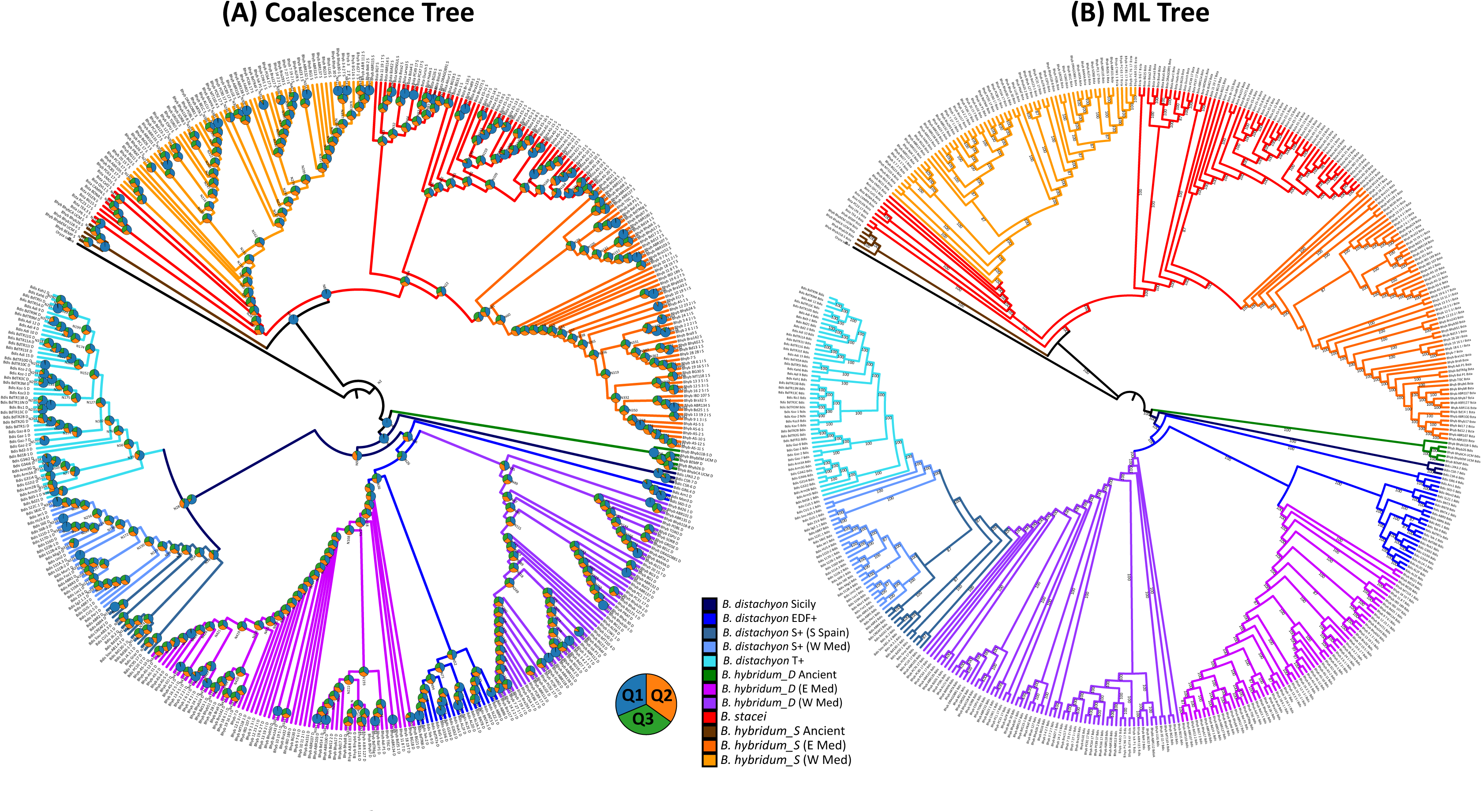
Comparison of Multi Species Coalescent (MSC) and Maximum-Likelihood (ML) phylogenies for the *Brachypodium distachyon – B. stacei – B. hybridum* complex. Left: MSC ASTRAL species tree inferred from windowed trees estimated across non-overlapping SNP-based 10 kb genomic windows; pie charts indicated the nodal quartet support for the optimal (q1) and the first (q2) and second (q3) alternative topologies. Right: IQTree2 ML tree inferred from the concatenated full-SNP alignment across all windows. Tip branches and lineages are colored by genomic group using the same scheme as in Fig. **2**: *B. distachyon* (blue), B. stacei (red), *B. hybridum* S-subgenome (orange) and *B. hybridum* D-subgenome (purple).

**Figure S2.**
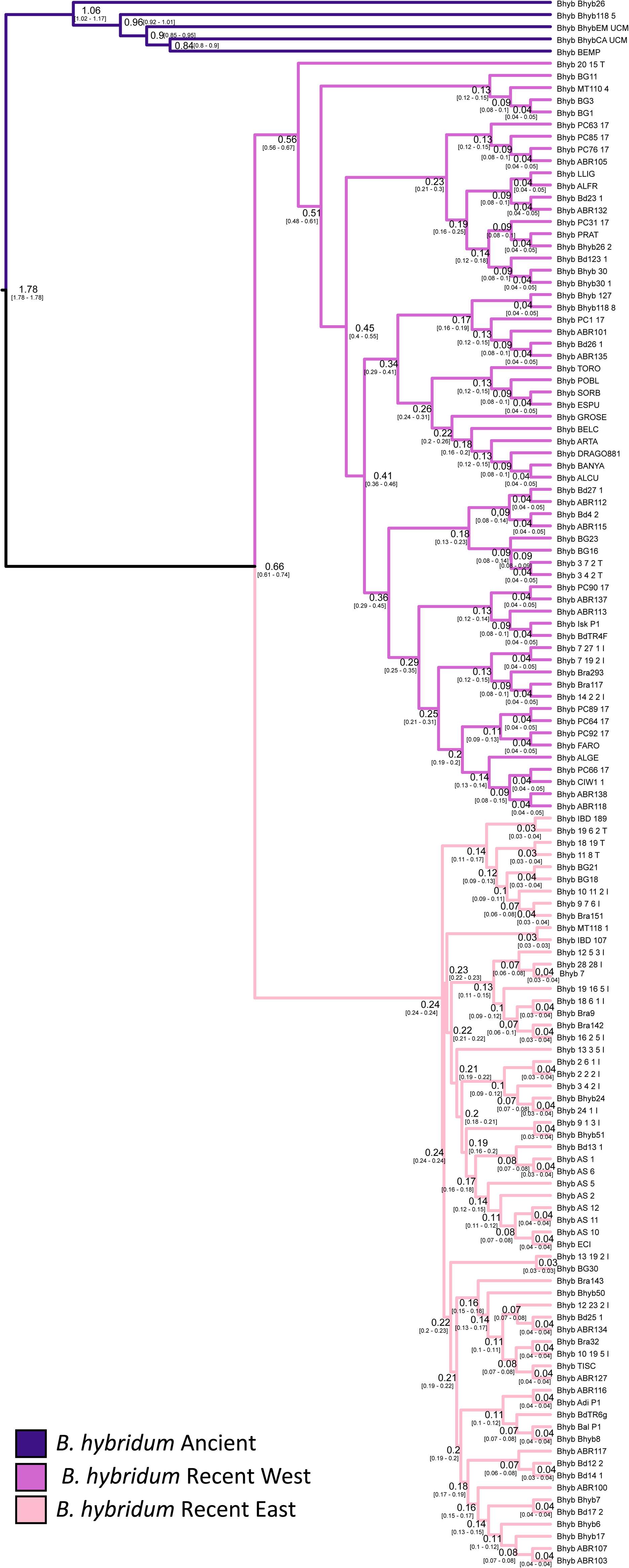
Divergence dating analysis of the nuclear *Brachypodium hybridum* phylogeny reconstructed with SVDQuartets and dataset #4 (983,994 SNPs) using a penalized likelihood method with TreePL. Secondary calibrations were imposed on key nodes [crown Ancient clade (2.16 Ma ± 0.5 Ma); crown Recent West clade (0.72 Ma ± 0.1 Ma); crown Recent East clade (0.13 Ma ± 0.1 Ma)] following Mu et al. (2023a). The optimal smoothing parameter was determined using cross-validation. Additionally, we generated 1,000 bootstrap trees from the SNP dataset and dated each with TreePL to assess temporal uncertainty. All resulting dated trees were then summarized with TreeAnnotator in BEAST2 to calculate the 95% highest-posterior-density intervals for each node.

**Figure S3.**
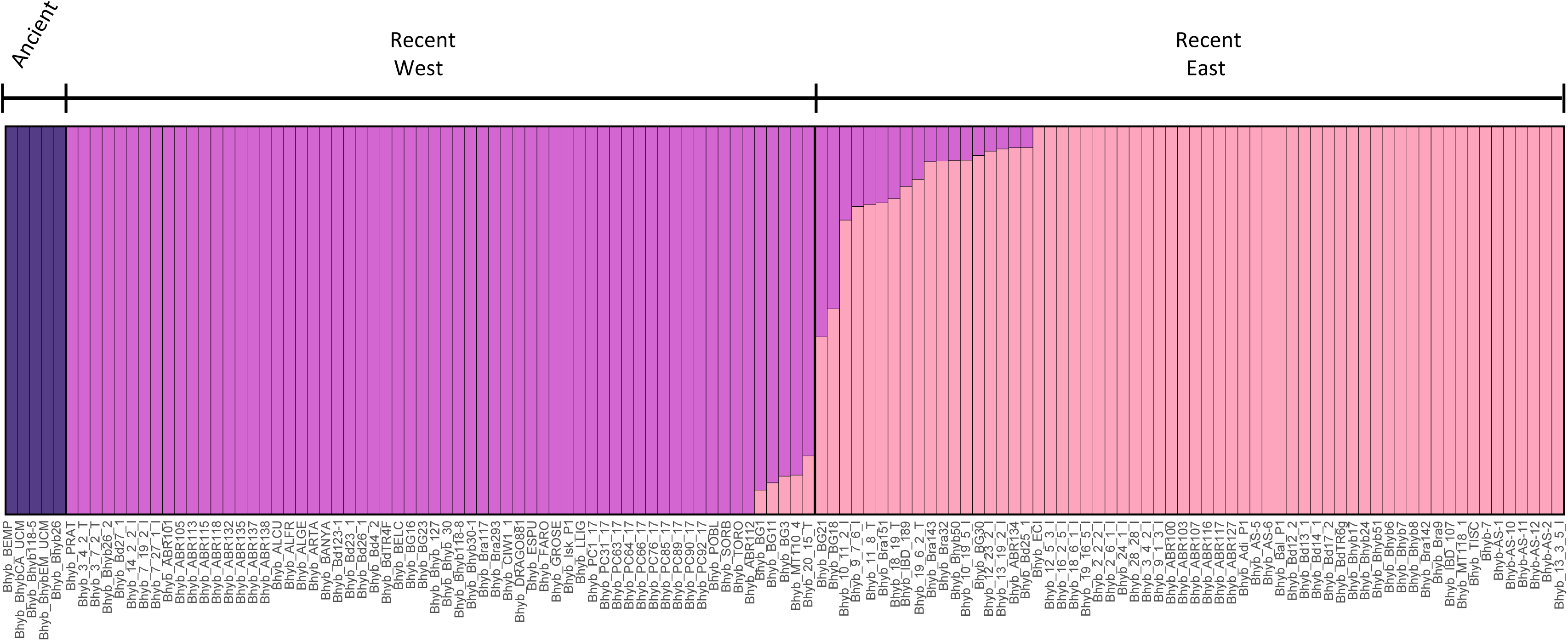
Individual-level Admixture barplot for *Brachypodium hybridum* at K = 3. Each vertical bar represents one of the 129 accessions, ordered by population origin (Ancient n = 5; Recent West n = 63; Recent East n = 61). Bar colors denote the three genetic clusters identified: Cluster 1 (Ancient–like) in mauve, Cluster 2 (Recent West–like) in wine, Cluster 3 (Recent East–like) in light pink. Membership coefficients (Q-values) are plotted as the proportional length of each color segment within a bar.

**Figure S4.**
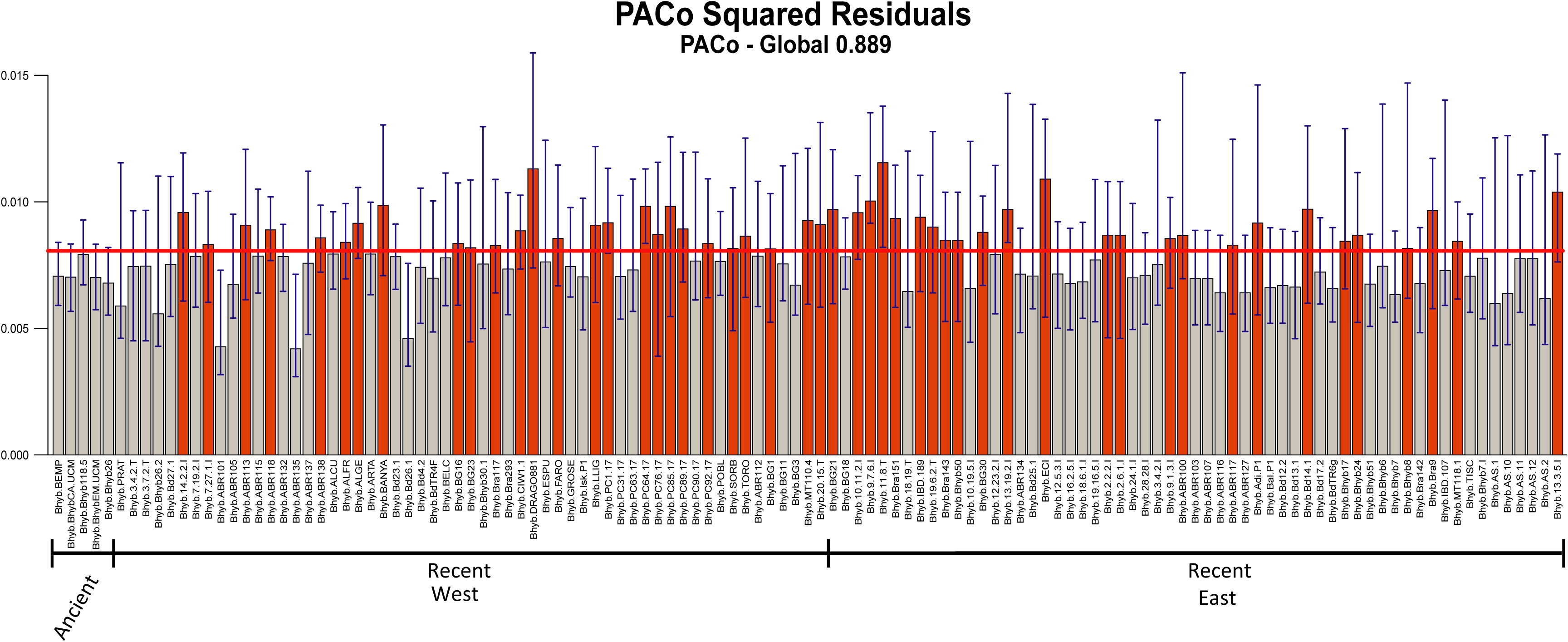
Bar plot illustrating normalized squared residuals from Procrustean Approach to Cophylogeny (PACo) analyses between phylogenetic trees of nuclear SNPs data and whole plastome data of *Brachypodium hybridum*. Each bar represents a specific nuclear-plastome association, showing the median of normalized squared residuals with 95% confidence intervals (blue lines). The red horizontal line indicates the incongruence threshold calculated with the Q3+Q4 quartile; bars exceeding this threshold represent incongruent associations (orange), suggesting independent evolutionary histories, while bars below denote congruent associations (grey), consistent with coevolutionary patterns.

**Figure S5.**
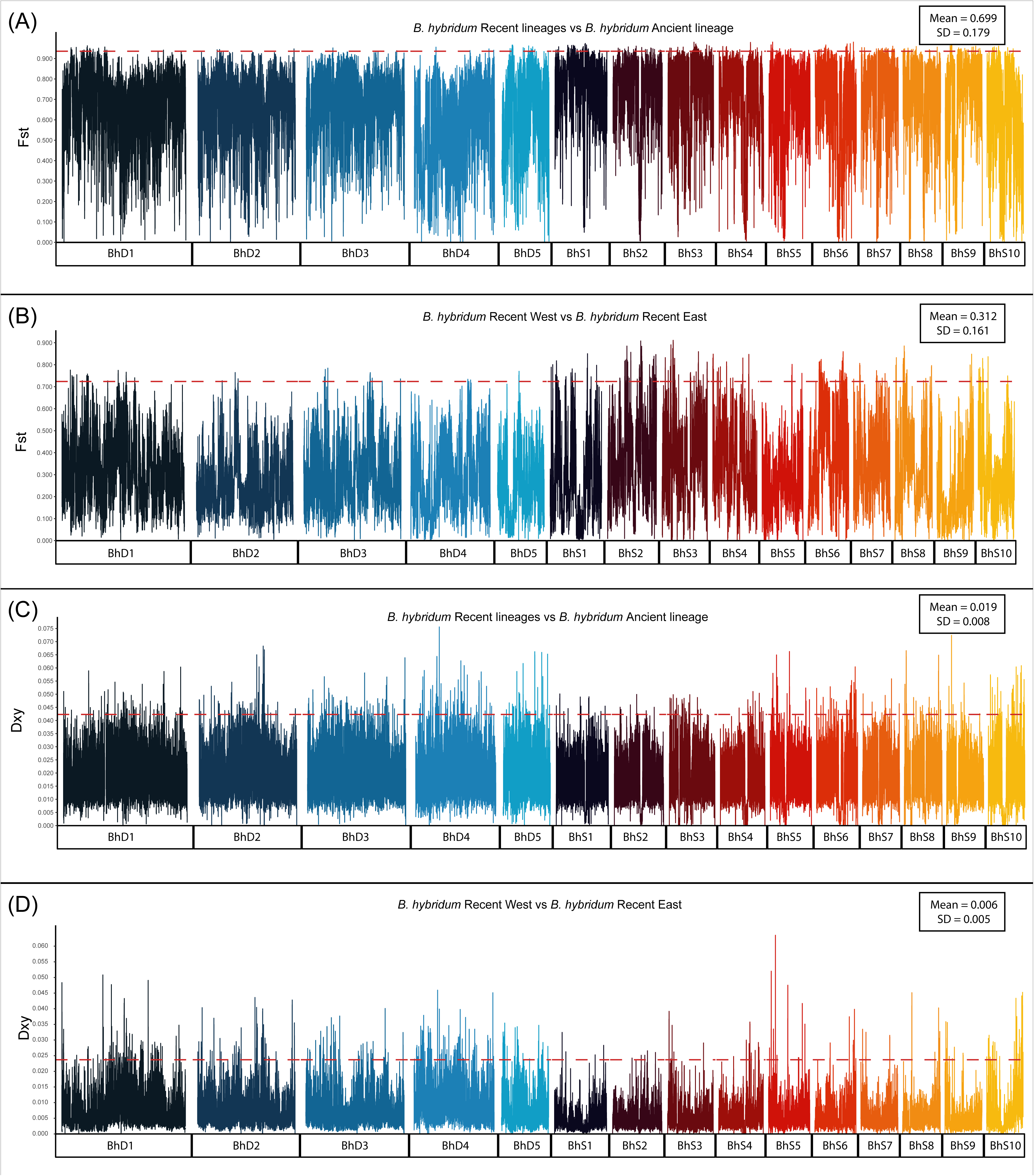
Chromosome-wide patterns of genetic differentiation and divergence among *Brachypodium hybridum* lineages. Pairwise Fst (panels A–B) and Dxy (panels C–D) were calculated in non-overlapping windows along chromosomes 1–15 (D chromosomes: BhD01-BhD05, plus S chromosomes: BhS1-BhS10). In panels A and C, comparisons are made between the Ancient lineage and the combined Recent lineages, whereas panels B and D show comparisons between Recent West and Recent East lineages. Each point corresponds to a windowed estimate (Fst in A–B; Dxy in C–D), and the horizontal dashed red line indicates the 95th-percentile genome-wide threshold for each statistic, above which windows are highlighted as outliers.

**Figure S6.**
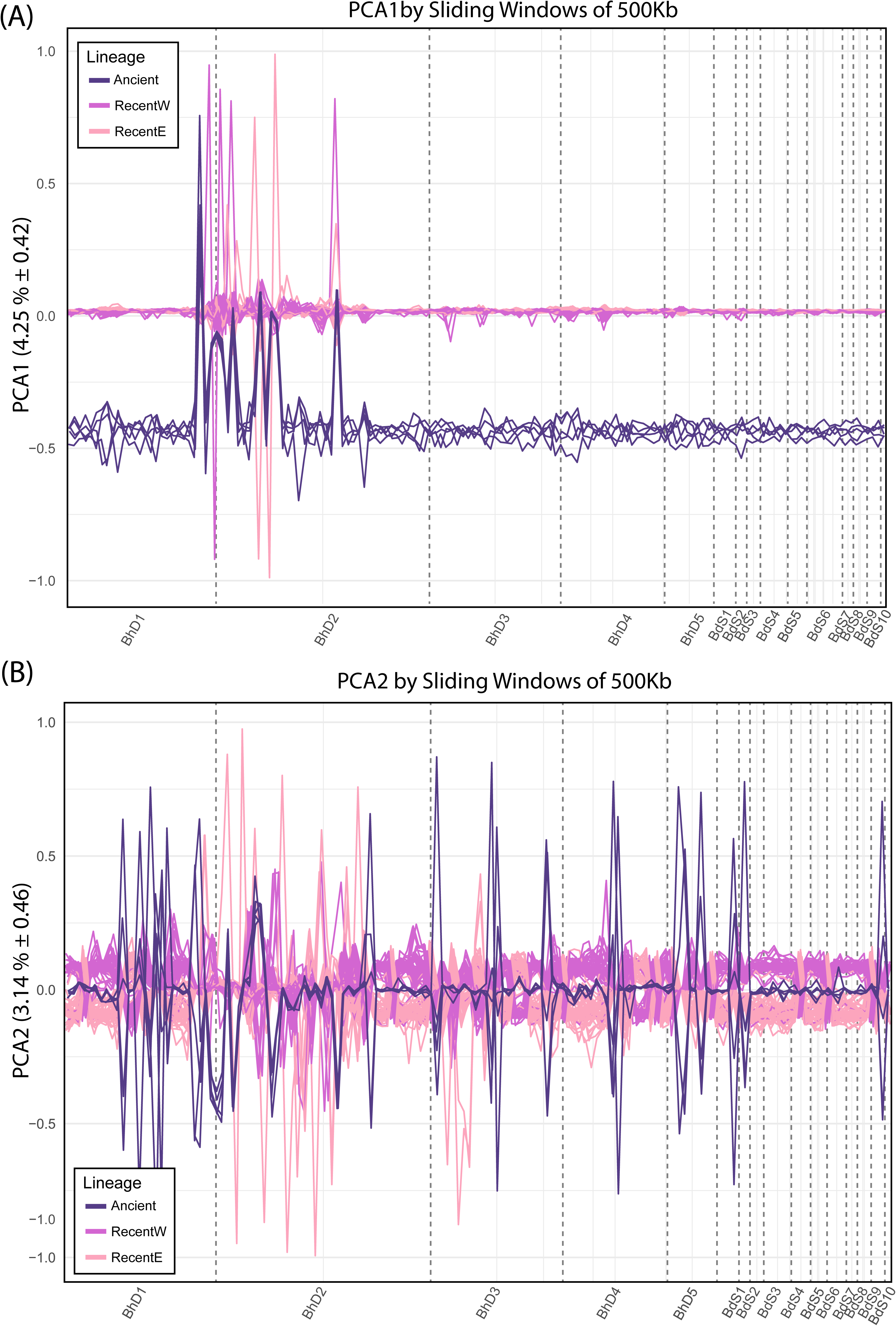
Principal Coordinates Analysis (PCoA) of *Brachypodium hybridum* per 1,000 Kb genomic windows, colored by Ancient, Recent West, and Recent East groups, showing PCA1 (A) and PCA2 (B).

**Table S1.** Samples of *Brachypodium distachyon*, *B. stacei* and *B. hybridum* from their native circum-Mediterranean region and some invaded areas used in the population genomic study, including species, accession, accession code (used in this study, includes species acronym and accession), plastome type (D: distachyon-type: S; stacei-type), location and geographic coordinates. Newly sequenced samples are highlighted in bold.

| Species | Accession | Accession Code<br>(used in this study) | Plastome type | Location | Longitude | Latitude |
| --- | --- | --- | --- | --- | --- | --- |
| *B. distachyon* | ABR2 | Bdis_ABR2 | D | France: Hérault | 43.591 | 3.258 |
| *B. distachyon* | ABR3 | Bdis_ABR3 | D | Spain: Huesca, Aísa | 42.695 | -0.682 |
| *B. distachyon* | ABR4 | Bdis_ABR4 | D | Spain: Huesca, Arén | 42.250 | 0.533 |
| *B. distachyon* | ABR5 | Bdis_ABR5 | D | Spain: Huesca, Jaca | 42.568 | -0.550 |
| *B. distachyon* | ABR6 | Bdis_ABR6 | D | Spain: Navarra, Los Arcos | 42.558 | -2.289 |
| *B. distachyon* | ABR7 | Bdis_ABR7 | D | Spain: Valladolid, Otero | 41.580 | -5.319 |
| *B. distachyon* | ABR9 | Bdis_ABR9 | D | Croatia: Ljubljana | 46.066 | 14.475 |
| *B. distachyon* | Adi_10 | Bdis_Adi_10 | D | Turkey: Adiyaman | 37.764 | 38.277 |
| *B. distachyon* | Adi_15 | Bdis_Adi_12 | D | Turkey: Adiyaman | 37.764 | 38.277 |
| *B. distachyon* | Adi_4 | Bdis_Adi_15 | D | Turkey: Adiyaman | 37.764 | 38.277 |
| *B. distachyon* | Adi_9 | Bdis_Adi_4 | D | Turkey: Adiyaman | 37.764 | 38.277 |
| *B. distachyon* | Adi_12 | Bdis_Adi_9 | D | Turkey: Adiyaman | 37.764 | 38.277 |
| *B. distachyon* | AL1A.1 | Bdis_AL1A.1 | D | Albania: Elbasan | 41.113 | 20.082 |
| *B. distachyon* | AL2D | Bdis_AL2D | D | Albania: Elbasan | 41.113 | 20.082 |
| *B. distachyon* | AL2E | Bdis_AL2E | D | Albania: Elbasan | 41.113 | 20.082 |
| *B. distachyon* | AL2F | Bdis_AL2F | D | Albania: Elbasan | 41.113 | 20.082 |
| *B. distachyon* | Arm2B | Bdis_Arm2B | D | Armenia: Vorotan | 39.429 | 46.025 |
| *B. distachyon* | Arm3A | Bdis_Arm3A | D | Armenia: Archi | 40.905 | 45.181 |
| *B. distachyon* | Arm3G | Bdis_Arm3G | D | Armenia: Archi | 40.905 | 45.181 |
| *B. distachyon* | Arm3i | Bdis_Arm3i | D | Armenia: Archi | 40.905 | 45.181 |
| *B. distachyon* | Arn1 | Bdis_Arn1 | D | Spain: Huesca, Arén | 42.250 | 0.533 |
| *B. distachyon* | Bd1-1 | Bdis_Bd1-1 | D | Turkey: Soma, Manisa | 39.187 | 27.608 |
| *B. distachyon* | Bd18-1 | Bdis_Bd18-1 | D | Turkey: Kaman, Kırşehir | 39.205 | 33.301 |
| *B. distachyon* | Bd2-3 | Bdis_Bd2-3 | D | Iraq | 33.223 | 43.679 |
| *B. distachyon* | Bd21 | Bdis_Bd21_reference | D | Iraq: near Salakudin | 33.223 | 43.679 |
| *B. distachyon* | Bd3-1 | Bdis_Bd3-1 | D | Iraq | 33.223 | 43.679 |
| *B. distachyon* | Bd30-1 | Bdis_Bd30-1 | D | Spain: Granada, Dilar | 37.042 | -3.545 |
| *B. distachyon* | BdTR10C | Bdis_BdTR10C | D | Turkey | 38.964 | 35.243 |
| *B. distachyon* | BdTR10D | Bdis_BdTR10D | D | Turkey | 38.964 | 35.243 |
| *B. distachyon* | BdTR11A | Bdis_BdTR11A | D | Turkey | 38.964 | 35.243 |
| *B. distachyon* | BdTR11E | Bdis_BdTR11E | D | Turkey | 38.964 | 35.243 |
| *B. distachyon* | BdTR11G | Bdis_BdTR11G | D | Turkey: Kırklareli | 41.733 | 27.217 |
| *B. distachyon* | BdTR11I | Bdis_BdTR11I | D | Turkey | 38.964 | 35.243 |
| *B. distachyon* | BdTR13B | Bdis_BdTR13B | D | Turkey: Ankara | 39.933 | 32.860 |
| *B. distachyon* | BdTR13C | Bdis_BdTR13C | D | Turkey: Ankara | 39.933 | 32.860 |
| *B. distachyon* | BdTR13N | Bdis_BdTR13N | D | Turkey: Ankara | 39.933 | 32.860 |
| *B. distachyon* | BdTR1i | Bdis_BdTR1i | D | Turkey: Aydın | 37.845 | 27.840 |
| *B. distachyon* | BdTR2B | Bdis_BdTR2B | D | Turkey | 38.964 | 35.243 |
| *B. distachyon* | BdTR2G | Bdis_BdTR2G | D | Turkey: Ankara | 39.933 | 32.860 |
| *B. distachyon* | BdTR3C | Bdis_BdTR3C | D | Turkey | 38.964 | 35.243 |
| *B. distachyon* | BdTR3M | Bdis_BdTR3M | D | Turkey | 38.964 | 35.243 |
| *B. distachyon* | BdTR5A | Bdis_BdTR5A | D | Turkey | 38.964 | 35.243 |
| *B. distachyon* | BdTR5I | Bdis_BdTR5I | D | Turkey | 38.964 | 35.243 |
| *B. distachyon* | BdTR7a | Bdis_BdTR7a | D | Turkey: Yozgat | 39.820 | 34.809 |
| *B. distachyon* | BdTR8i | Bdis_BdTR8i | D | Turkey | 38.964 | 35.243 |
| *B. distachyon* | BdTR9K | Bdis_BdTR9K | D | Turkey: Eskişehir | 39.767 | 30.526 |
| *B. distachyon* | BdTR9M | Bdis_BdTR9M | D | Turkey | 38.964 | 35.243 |
| *B. distachyon* | Bd130-1 | Bdis_Bd130-1 | D | Spain: Jaen: Fontanar: Guadiana Menor | 38.499 | -3.469 |
| *B. distachyon* | Bd162 | Bdis_Bd162 | D | Spain: Jaen: Srra Segura: La Vaquilla. | 38.148 | -3.105 |
| *B. distachyon* | Bis1 | Bdis_Bis1 | D | Turkey: Bismil | 37.844 | 40.789 |
| *B. distachyon* | BNT3 | Bdis_BNT3 | D | Italy: Sicily, Bronte | 37.790 | 14.836 |
| *B. distachyon* | BNT4.1 | Bdis_BNT4.1 | D | Italy: Sicily, Bronte | 37.790 | 14.836 |
| *B. distachyon* | BNT8 | Bdis_BNT8 | D | Italy: Sicily, Bronte | 37.790 | 14.836 |
| ***B. distachyon*** | **CREAF2** | **Bdis_CREAF2** | **D** | **Spain: Barcelona, Garraf** | **41.256** | **1.903** |
| *B. distachyon* | CSR-6 | Bdis_CSR-6 | D | Italy: Sicily, Cesaro | 37.868 | 14.677 |
| *B. distachyon* | CSR-7 | Bdis_CSR-7 | D | Italy: Sicily, Cesaro | 37.868 | 14.677 |
| *B. distachyon* | CU1.6-1 | Bdis_CU1.6-1 | D | Spain: Cuenca, near Gabaldón | 39.967 | -2.383 |
| *B. distachyon* | CU5.5 | Bdis_CU5.5 | D | Spain: Cuenca: Serranía de Cuenca | 40.219 | -1.776 |
| *B. distachyon* | Foz1 | Bdis_Foz1 | D | Spain: Navarra, Foz de Lumbier | 42.566 | -1.302 |
| *B. distachyon* | G30i2 | Bdis_G30i2 | D | Georgia | 42.315 | 43.357 |
| *B. distachyon* | G31i4 | Bdis_G31i4 | D | Georgia | 42.315 | 43.357 |
| *B. distachyon* | G32i2 | Bdis_G32i2 | D | Georgia | 42.315 | 43.357 |
| *B. distachyon* | G33i4 | Bdis_G33i4 | D | Georgia | 42.315 | 43.357 |
| *B. distachyon* | G33i6 | Bdis_G33i6 | D | Georgia | 42.315 | 43.357 |
| *B. distachyon* | G34i2 | Bdis_G34i2 | D | Georgia | 42.315 | 43.357 |
| *B. distachyon* | G34i6 | Bdis_G34i6 | D | Georgia | 42.315 | 43.357 |
| *B. distachyon* | Gaz-1 | Bdis_Gaz-1 | D | Turkey: Gaziantep | 37.066 | 37.383 |
| *B. distachyon* | Gaz-2 | Bdis_Gaz-2 | D | Turkey: Gaziantep | 37.066 | 37.383 |
| *B. distachyon* | Gaz-7 | Bdis_Gaz-7 | D | Turkey: Gaziantep | 37.066 | 37.383 |
| *B. distachyon* | Gaz-8 | Bdis_Gaz-8 | D | Turkey: Gaziantep | 37.066 | 37.383 |
| *B. distachyon* | GR6.4 | Bdis_GR6.4 | D | Spain: Granada, P. N. Sierra de Baza | 37.513 | -2.983 |
| ***B. distachyon*** | **Graz** | **Bdis_Graz** | **D** | **Spain: Cádiz, Grazalema** | **36.762** | **-5.396** |
| *B. distachyon* | GU5-1 | Bdis_GU5-1 | D | Spain: Guadalajara, Valsalobre | 40.901 | -1.950 |
| *B. distachyon* | HU3.4 | Bdis_HU3.4 | D | Spain: Huesca, El Grado | 42.117 | 0.367 |
| *B. distachyon* | J4.3.1 | Bdis_J4.3.1 | D | Spain: jaen: Puerto de Tiscar | 37.886 | -3.169 |
| *B. distachyon* | J6.2 | Bdis_J6.2 | D | Spain: Jaen, P. N. Sierra de Segura | 38.209 | -2.743 |
| *B. distachyon* | Jer1 | Bdis_Jer1 | D | Spain: Huesca, Peraltila, Ermita de San Jerónimo | 41.878 | 0.108 |
| *B. distachyon* | Kah6 | Bdis_Kah1 | D | Turkey: Kahta | 37.770 | 38.626 |
| *B. distachyon* | Kah1 | Bdis_Kah6 | D | Turkey: Kozluk | 38.190 | 41.486 |
| *B. distachyon* | Koz-1 | Bdis_Koz-1 | D | Turkey: Kozluk | 38.190 | 41.486 |
| *B. distachyon* | Koz-2 | Bdis_Koz-2 | D | Turkey: Kozluk | 38.190 | 41.486 |
| *B. distachyon* | Koz-5 | Bdis_Koz-5 | D | Turkey: Kozluk | 38.190 | 41.486 |
| *B. distachyon* | Koz3 | Bdis_Koz3 | D | Turkey: Kozluk | 38.190 | 41.486 |
| *B. distachyon* | LPA3.2 | Bdis_LPA3.2 | D | Italy: Sicily, Lago di Piana degli Albanesi | 37.985 | 13.287 |
| *B. distachyon* | Luc1 | Bdis_Luc1 | D | Spain: Huesca, Berdún, Ermita de Santa Lucía | 42.663 | -0.933 |
| *B. distachyon* | Mig3 | Bdis_Mig3 | D | Spain: Huesca, Ibieca, San Miguel de Foces | 42.165 | 0.512 |
| *B. distachyon* | Mon3 | Bdis_Mon3 | D | Spain: Huesca, Castejón de Monegros, Pto. Pallaruelo | 41.943 | 0.126 |
| *B. distachyon* | Mur1 | Bdis_Mur1 | D | Spain: Lleida, Castillo de Mur | 42.095 | 0.685 |
| *B. distachyon* | Nvk1 | Bdis_Nvk1 | D | Ukraine: Krym | 45.251 | 34.050 |
| ***B. distachyon*** | **PC10_17** | **Bdis_PC10_17** | **D** | **Morocco: Talassemtane National Park** | **35.127** | **-5.220** |
| ***B. distachyon*** | **PC95_17** | **Bdis_PC95_17** | **D** | **Morocco: Alcazarseguir** | **35.171** | **-5.191** |
| *B. distachyon* | Per1 | Bdis_Per1 | D | Spain: Navarra, Puerto del Perdón | 42.679 | -1.647 |
| *B. distachyon* | S11A.3 | Bdis_S11A.3 | D | Spain: Aragón | 41.649 | -0.889 |
| *B. distachyon* | S11B-2 | Bdis_S11B-2 | D | Spain: Aragón | 41.649 | -0.889 |
| *B. distachyon* | S11D-2 | Bdis_S11D-2 | D | Spain: Aragón | 41.649 | -0.889 |
| *B. distachyon* | S12B-4 | Bdis_S12B-4 | D | Spain: Aragón | 41.649 | -0.889 |
| *B. distachyon* | S16D | Bdis_S16D | D | Spain: Aragón | 41.649 | -0.889 |
| *B. distachyon* | S17D-1 | Bdis_S17D-1 | D | Spain: Aragón | 41.649 | -0.889 |
| *B. distachyon* | S19A | Bdis_S19A | D | Spain: Aragón | 41.649 | -0.889 |
| *B. distachyon* | S22B-1 | Bdis_S22B-1 | D | Spain: Aragón | 41.649 | -0.889 |
| *B. distachyon* | S22C.1 | Bdis_S22C.1 | D | Spain: Aragón | 41.649 | -0.889 |
| *B. distachyon* | S6B-3 | Bdis_S6B-3 | D | Spain: Aragón | 41.649 | -0.889 |
| *B. distachyon* | S6D-5 | Bdis_S6D-5 | D | Spain: Aragón | 41.649 | -0.889 |
| *B. distachyon* | S6E | Bdis_S6E | D | Spain: Aragón | 41.649 | -0.889 |
| *B. distachyon* | S8iiC | Bdis_S8iiC | D | Spain: Aragón | 41.874 | 0.184 |
| *B. distachyon* | Sg2.1.1 | Bdis_Sg2.1.1 | D | Spain: Segovia, Carabias | 41.068 | -3.426 |
| *B. distachyon* | SLZ2.1 | Bdis_SLZ2.1 | D | Italy: Sicily, Schilizzi | 38.185 | 14.494 |
| *B. distachyon* | Sou-AB1.4-2 | Bdis_Sou-AB1.4-2 | D | Spain: Albacete, near Riopar | 38.367 | -2.417 |
| *B. distachyon* | Sou-AB3.3 | Bdis_Sou-AB3.3 | D | Spain: Albacete, between Gineta and Tarazona | 39.117 | -1.783 |
| *B. distachyon* | Tek-2 | Bdis_Tek-2 | D | Turkey: Tekirdag | 40.983 | 27.517 |
| *B. distachyon* | Tek-9 | Bdis_Tek-9 | D | Turkey: Tekirdag | 40.983 | 27.517 |
| *B. distachyon* | Uni2 | Bdis_Uni2 | D | Spain: Huesca, Escuela Politécnica Superior | 42.133 | -0.417 |
| *B. distachyon* | Z3.6 | Bdis_Z3.6 | D | Spain: Zaragoza: Torralba de los Frailes | 41.150 | -0.383 |
| ***B. stacei*** | **11_19_1_T** | **Bsta_11_19_1_T** | **S** | **Turkey** | **39.534** | **2.858** |
| *B. stacei* | ABR114 | Bsta_ABR114_reference | S | Spain: Majorca | 39.534 | 2.858 |
| ***B. stacei*** | **Alsur1** | **Bsta_Alsur1** | **S** | **Spain: Almeria, Sorbas** | **37.1** | **-2.131** |
| *B. stacei* | AS-18 | Bsta_AS-18 | S | Israel: Evolution Canyon I | 32.71 | 34.97 |
| *B. stacei* | AS-19 | Bsta_AS-19 | S | Israel: Evolution Canyon I | 32.71 | 34.97 |
| *B. stacei* | AS-20 | Bsta_AS-20 | S | Israel: Evolution Canyon I | 32.71 | 34.97 |
| *B. stacei* | AS-21 | Bsta_AS-21 | S | Israel: Evolution Canyon I | 32.71 | 34.97 |
| *B. stacei* | AS-22 | Bsta_AS-22 | S | Israel: Evolution Canyon I | 32.71 | 34.97 |
| *B. stacei* | AS-23 | Bsta_AS-23 | S | Israel: Evolution Canyon I | 32.71 | 34.97 |
| *B. stacei* | AS-24 | Bsta_AS-24 | S | Israel: Evolution Canyon I | 32.71 | 34.97 |
| *B. stacei* | AS-25 | Bsta_AS-25 | S | Israel: Evolution Canyon I | 32.71 | 34.97 |
| *B. stacei* | AS-26 | Bsta_AS-26 | S | Israel: Evolution Canyon I | 32.71 | 34.97 |
| *B. stacei* | AS-27 | Bsta_AS-27 | S | Israel: Evolution Canyon I | 32.71 | 34.97 |
| *B. stacei* | AS-28 | Bsta_AS-28 | S | Israel: Evolution Canyon I | 32.71 | 34.97 |
| *B. stacei* | AS-29 | Bsta_AS-29 | S | Israel: Evolution Canyon I | 32.71 | 34.97 |
| *B. stacei* | AS-30 | Bsta_AS-30 | S | Israel: Evolution Canyon I | 32.71 | 34.97 |
| *B. stacei* | AS-31 | Bsta_AS-31 | S | Israel: Evolution Canyon I | 32.71 | 34.97 |
| *B. stacei* | AS-6 | Bsta_AS-6 | S | Israel: Evolution Canyon I | 32.71 | 34.97 |
| *B. stacei* | **BG15** | **Bsta_BG15** | **S** | **Greece: Tyros** | **37.24** | **22.862** |
| *B. stacei* | **BG19** | **Bsta_BG19** | **S** | **Greece: Thera** | **36.393** | **25.462** |
| *B. stacei* | **BG26** | **Bsta_BG26** | **S** | **Greece: Falassarna** | **35.467** | **23.592** |
| *B. stacei* | **BG31** | **Bsta_BG31** | **S** | **Greece: Preveli** | **35.162** | **24.438** |
| *B. stacei* | **BG7** | **Bsta_BG7** | **S** | **Greece: Trizina** | **37.602** | **23.452** |
| *B. stacei* | **BJAE2** | **Bsta_BJAE2** | **S** | **Spain: Jaen, Quesada, Tiscar** | **37.841** | **-3.13** |
| *B. stacei* | **BONA** | **Bsta_BONA** | **S** | **Spain: Majorca, Petra, Bonany** | **39.616** | **3.057** |
| *B. stacei* | Br-135 | Bsta_Br-135 | S | Israel: Lakhish | 31.55 | 34.85 |
| *B. stacei* | **Bra223** | **Bsta_Bra223** | **S** | **Israel: Modi'in** | **31.901** | **35.011** |
| *B. stacei* | **Bra271** | **Bsta_Bra271** | **S** | **Israel: Gilon-Zurit** | **33.031** | **35.371** |
| *B. stacei* | **Bra89** | **Bsta_Bra89** | **S** | **Israel: Ziff** | **32.998** | **35.229** |
| *B. stacei* | Bsta_ECI | Bsta_ECI_reference | S | Israel: Evolution Canyon I | 32.71 | 34.97 |
| *B. stacei* | Bsta-1 | Bsta-1 | S | Israel: Hazazon | 31.55 | 35.4 |
| *B. stacei* | **Bsta2** | **Bsta2** | **S** | **Spain: Jaen, Baeza** | **37.995** | **-3.471** |
| *B. stacei* | **Bsta4** | **Bsta4** | **S** | **Spain: Granada, Moclín** | **37.315** | **-4.003** |
| *B. stacei* | **Bsta5** | **Bsta5** | **S** | **Spain: Alicante, Cabo la Nao** | **38.742** | **0.18** |
| *B. stacei* | **CAMPA** | **Bsta_CAMPA** | **S** | **Spain: Majorca, Campanet, Coves** | **39.806** | **3.033** |
| *B. stacei* | Cef2 | Bsta_Cef2 | S | Italy: Sicily, Chiesa di Cefalu | 14.02 | 38.03 |
| *B. stacei* | **DRAGO891** | **Bsta_DRAGO891** | **S** | **Spain: Majorca, Sa Dragonera, Gambes** | **39.596** | **2.313** |
| *B. stacei* | ES-12 | Bsta_ES-12 | S | Israel: Evolution Canyon I | 32.71 | 34.97 |
| *B. stacei* | ES-13 | Bsta_ES-13 | S | Israel: Evolution Canyon I | 32.71 | 34.97 |
| *B. stacei* | ES-14 | Bsta_ES-14 | S | Israel: Evolution Canyon I | 32.71 | 34.97 |
| *B. stacei* | ES-15 | Bsta_ES-15 | S | Israel: Evolution Canyon I | 32.71 | 34.97 |
| *B. stacei* | ES-16 | Bsta_ES-16 | S | Israel: Evolution Canyon I | 32.71 | 34.97 |
| *B. stacei* | ES-17 | Bsta_ES-17 | S | Israel: Evolution Canyon I | 32.71 | 34.97 |
| *B. stacei* | ES-18 | Bsta_ES-18 | S | Israel: Evolution Canyon I | 32.71 | 34.97 |
| *B. stacei* | ES-19 | Bsta_ES-19 | S | Israel: Evolution Canyon I | 32.71 | 34.97 |
| *B. stacei* | ES-2 | Bsta_ES-2 | S | Israel: Evolution Canyon I | 32.71 | 34.97 |
| *B. stacei* | ES-20 | Bsta_ES-20 | S | Israel: Evolution Canyon I | 32.71 | 34.97 |
| *B. stacei* | ES-21 | Bsta_ES-21 | S | Israel: Evolution Canyon I | 32.71 | 34.97 |
| *B. stacei* | ES-22 | Bsta_ES-22 | S | Israel: Evolution Canyon I | 32.71 | 34.97 |
| *B. stacei* | ES-23 | Bsta_ES-23 | S | Israel: Evolution Canyon I | 32.71 | 34.97 |
| *B. stacei* | ES-24 | Bsta_ES-24 | S | Israel: Evolution Canyon I | 32.71 | 34.97 |
| *B. stacei* | ES-25 | Bsta_ES-25 | S | Israel: Evolution Canyon I | 32.71 | 34.97 |
| *B. stacei* | **FELEN** | **Bsta_FELEN** | **S** | **Spain: Majorca, Felenitx, San Salvador** | **39.43** | **3.194** |
| *B. stacei* | **Hada** | **Bsta_Hada** | **S** | **Tunisia: Hadada** | **36.775** | **10.227** |
| *B. stacei* | **IBD_10** | **Bsta_IBD_10** | **S** | **Israel** | **31.046** | **34.852** |
| *B. stacei* | **Larva3** | **Bsta_Larva3** | **S** | **Spain: Jaen, Larva** | **37.76** | **-3.207** |
| *B. stacei* | **LP6_1** | **Bsta_LP6_1** | **S** | **Spain: Canary Islands, Lanzarote, Teguisse** | **27.938** | **-15.685** |
| *B. stacei* | **MONSUL** | **Bsta_MONSUL** | **S** | **Spain: Almeria, Playa de Monsul** | **36.733** | **-2.145** |
| *B. stacei* | **Ourich** | **Bsta_Ourich** | **S** | **Morocco: Ourich** | **32.237** | **-6.324** |
| *B. stacei* | **PC23_17** | **Bsta_PC23_17** | **S** | **Morocco: Xaouen** | **35.172** | **-5.269** |
| *B. stacei* | **PC47_17** | **Bsta_PC47_17** | **S** | **Morocco: Beni Ferten** | **32.399** | **-6.367** |
| *B. stacei* | **PC49_17** | **Bsta_PC49_17** | **S** | **Morocco: Targha** | **32.28** | **-6.334** |
| *B. stacei* | **QVL1** | **Bsta_QVL1** | **S** | **Italy: Sicily, Quattro Vanelle. SH QVL1** | **37.846** | **13.13** |
| *B. stacei* | TE4.3 | Bsta_TE4.3 | S | Spain: Canary Islands, Gomera, Agulo | 27.938 | -15.685 |
| *B. stacei* | **VMO1** | **Bsta_VMO1** | **S** | **Italy: Sicily, Valdesi/Mondello. SH VMO1** | **38.202** | **13.332** |
| *B. hybridum* | **10_11_2_I** | **Bhyb_10_11_2_I** | **S** | **Israel: Carmel-Rakafot** | **32.734** | **35.032** |
| *B. hybridum* | **10_19_5_I** | **Bhyb_10_19_5_I** | **S** | **Israel: Carmel-Rakafot** | **32.734** | **35.032** |
| *B. hybridum* | **11_8_T** | **Bhyb_11_8_T** | **S** | **Turkey: Ölüdeniz** | **36.55** | **29.12** |
| *B. hybridum* | **12_23_2_I** | **Bhyb_12_23_2_I** | **S** | **Israel: Carmel-Oren** | **32.734** | **35.032** |
| *B. hybridum* | **12_5_3_I** | **Bhyb_12_5_3_I** | **S** | **Israel: Carmel-Oren** | **32.734** | **35.032** |
| *B. hybridum* | 13_19_2_I | Bhyb_13_19_2_I | S | Israel: Modi'in | 31.917 | 34.957 |
| *B. hybridum* | 13_3_5_I | Bhyb_13_3_5_I | S | Israel: Modi'in | 31.917 | 34.957 |
| *B. hybridum* | 14_2_2_I | Bhyb_14_2_2_I | S | Israel: Kiryat Anavim | 31.812 | 35.123 |
| *B. hybridum* | 16_2_5_I | Bhyb_16_2_5_I | S | Israel: Bet-Shemesh | 31.729 | 34.974 |
| *B. hybridum* | 18_19_T | Bhyb_18_19_T | S | Turkey: Alanya | 36.54 | 31.99 |
| *B. hybridum* | 18_6_1_I | Bhyb_18_6_1_I | S | Israel: Amatzya | 31.532 | 34.912 |
| *B. hybridum* | 19_16_5_I | Bhyb_19_16_5_I | S | Israel: Lahav | 31.376 | 34.87 |
| *B. hybridum* | 19_6_2_T | Bhyb_19_6_2_T | S | Turkey: Anamur | 36.08 | 32.821 |
| *B. hybridum* | 2_2_2_I | Bhyb_2_2_2_I | S | Israel: Hermon, Majdal | 33.255 | 35.746 |
| *B. hybridum* | 2_6_1_I | Bhyb_2_6_1_I | S | Israel: Hermon, Majdal | 33.255 | 35.746 |
| *B. hybridum* | 20_15_T | Bhyb_20_15_T | S | Turkey: Mersin | 36.783 | 34.536 |
| *B. hybridum* | 24_1_I | Bhyb_24_1_I | S | Israel: Sde Boker | 30.875 | 34.79 |
| *B. hybridum* | 28_28_I | Bhyb_28_28_I | S | Israel: Ha'Negev | 31.369 | 34.716 |
| *B. hybridum* | 3_4_2_I | Bhyb_3_4_2_I | S | Israel: Hermon | 33.255 | 35.746 |
| *B. hybridum* | 3_4_2_T | Bhyb_3_4_2_T | S | Turkey: Bahçeköy | 41.171 | 28.993 |
| *B. hybridum* | 3_7_2_T | Bhyb_3_7_2_T | S | Turkey: Bahçeköy | 41.171 | 28.993 |
| *B. hybridum* | 7_19_2_I | Bhyb_7_19_2_I | S | Israel: Shtula | 33.087 | 35.312 |
| *B. hybridum* | 7_27_1_I | Bhyb_7_27_1_I | S | Israel: Shtula | 33.087 | 35.312 |
| *B. hybridum* | 9_1_3_I | Bhyb_9_1_3_I | S | Israel: Karmiel | 32.917 | 35.288 |
| *B. hybridum* | 9_7_6_I | Bhyb_9_7_6_I | S | Israel: Karmiel | 32.917 | 35.288 |
| *B. hybridum* | ABR100 | Bhyb_ABR100 | S | Iran: Kalafabad | 30.508 | 49.841 |
| *B. hybridum* | ABR101 | Bhyb_ABR101 | S | South Africa: Darling | NA | NA |
| *B. hybridum* | ABR103 | Bhyb_ABR103 | S | Iran: Shustar | 32.055 | 48.849 |
| *B. hybridum* | ABR105 | Bhyb_ABR105 | S | Morocco: Nr Ongda | 31.792 | -7.093 |
| *B. hybridum* | ABR107 | Bhyb_ABR107 | S | Greece | 39.074 | 21.824 |
| *B. hybridum* | ABR112 | Bhyb_ABR112 | S | Iraq | 33.223 | 43.679 |
| *B. hybridum* | ABR113 | Bhyb_ABR113_reference | S | Portugal: Lisbon | 38.733 | -9.18 |
| *B. hybridum* | ABR115 | Bhyb_ABR115 | S | South Africa | NA | NA |
| *B. hybridum* | ABR116 | Bhyb_ABR116 | S | Afghanistan | 33.939 | 67.71 |
| *B. hybridum* | ABR117 | Bhyb_ABR117 | S | Afghanistan | 33.939 | 67.71 |
| *B. hybridum* | ABR118 | Bhyb_ABR118 | S | Afghanistan | 33.939 | 67.71 |
| *B. hybridum* | ABR127 | Bhyb_ABR127 | S | Iran | 32.428 | 53.688 |
| *B. hybridum* | ABR132 | Bhyb_ABR132 | S | Spain | NA | NA |
| *B. hybridum* | ABR134 | Bhyb_ABR134 | S | Israel | 31.046 | 34.852 |
| *B. hybridum* | ABR135 | Bhyb_ABR135 | S | Uruguay | NA | NA |
| *B. hybridum* | ABR137 | Bhyb_ABR137 | S | Australia | NA | NA |
| *B. hybridum* | ABR138 | Bhyb_ABR138 | S | Portugal: Quinta da Pacheca | 41.155 | -7.799 |
| *B. hybridum* | Adi_P1 | Bhyb_Adi_P1 | S | Turkey | 38.964 | 35.243 |
| *B. hybridum* | ALCU | Bhyb_ALCU | S | Spain: Majorca, Alcudia, Punta Negra | 39.923 | 3.067 |
| *B. hybridum* | ALFR | Bhyb_ALFR | S | Spain: Zaragoza, Alfranca | 41.608 | -0.752 |
| *B. hybridum* | ALGE | Bhyb_ALGE | S | Spain: Cádiz, Algeciras | 36.124 | -5.475 |
| *B. hybridum* | ARTA | Bhyb_ARTA | S | Spain: Mallorca, Artá | 39.736 | 3.335 |
| *B. hybridum* | Bal_P1 | Bhyb_Bal_P1 | S | Turkey: Ballı | 41.005 | 42.184 |
| *B. hybridum* | BANYA | Bhyb_BANYA | S | Spain: Majorca, Banyalbufar, Ses Animes | 39.682 | 2.51 |
| *B. hybridum* | Bd12_2 | Bhyb_Bd12_2 | S | Turkey | 38.964 | 35.243 |
| *B. hybridum* | Bd13_1 | Bhyb_Bd13_1 | S | Turkey | 38.964 | 35.243 |
| *B. hybridum* | Bd14_1 | Bhyb_Bd14_1 | S | Turkey | 38.964 | 35.243 |
| *B. hybridum* | Bd17_2 | Bhyb_Bd17_2 | S | Turkey | 38.964 | 35.243 |
| *B. hybridum* | Bd23_1 | Bhyb_Bd23_1 | S | Turkey | 38.964 | 35.243 |
| *B. hybridum* | Bd25_1 | Bhyb_Bd25_1 | S | Turkey | 38.964 | 35.243 |
| *B. hybridum* | Bd26_1 | Bhyb_Bd26_1 | S | Turkey | 38.964 | 35.243 |
| *B. hybridum* | Bd27_1 | Bhyb_Bd27_1 | S | Turkey | 38.964 | 35.243 |
| *B. hybridum* | Bd4_2 | Bhyb_Bd4_2 | S | Turkey | 38.964 | 35.243 |
| *B. hybridum* | BdTR4F | Bhyb_BdTR4F | S | Turkey | 38.964 | 35.243 |
| *B. hybridum* | BdTR6g | Bhyb_BdTR6g | S | Turkey | 38.964 | 35.243 |
| *B. hybridum* | BELC | Bhyb_BELC | S | Spain: Zaragoza, Belchite | 41.305 | -0.752 |
| *B. hybridum* | BEMP | Bhyb_BEMP | D | Portugal: Trás-os-Montes, Bemposta | 41.306 | -6.495 |
| *B. hybridum* | BG1 | Bhyb_BG1 | S | Greece: Epidauros | 37.621 | 23.129 |
| *B. hybridum* | BG11 | Bhyb_BG11 | S | Greece: Arkadiko | 37.596 | 22.956 |
| *B. hybridum* | BG16 | Bhyb_BG16 | S | Greece: Poulithra | 37.111 | 22.886 |
| *B. hybridum* | BG18 | Bhyb_BG18 | S | Greece: Kamari | 36.38 | 25.481 |
| *B. hybridum* | BG21 | Bhyb_BG21 | S | Greece: Therasia | 36.433 | 25.338 |
| *B. hybridum* | BG23 | Bhyb_BG23 | S | Greece: Chania | 35.507 | 24.02 |
| *B. hybridum* | BG3 | Bhyb_BG3 | S | Greece: Kosta | 37.298 | 23.168 |
| *B. hybridum* | BG30 | Bhyb_BG30 | S | Greece: Platias | 35.188 | 24.404 |
| *B. hybridum* | Bhyb123-1 | Bhyb_Bhyb123-1 | S | Spain: Jaen: Srta Segura: Cortijos Nuevos | 38.248 | -2.722 |
| *B. hybridum* | Bhyb_127 | Bhyb_Bhyb_127 | S | Spain: Jaen: Hinojares | 37.715 | -2.999 |
| *B. hybridum* | Bhyb24 | Bhyb_Bhyb24 | S | Israel: Sde Boker | 30.875 | 34.79 |
| *B. hybridum* | Bhyb_30 | Bhyb_Bhyb_30 | S | Spain: Huelva, Lepe | 37.307 | -7.222 |
| *B. hybridum* | Bhyb30-1 | Bhyb_Bhyb30-1 | S | Spain: Huelva, Lepe | 37.307 | -7.222 |
| *B. hybridum* | Bhyb118-5 | Bhyb_Bhyb118-5 | D | Spain: Almeria: Cabo de Gata: Playa Genoveses | 36.746 | -2.122 |
| *B. hybridum* | Bhyb118-8 | Bhyb_Bhyb118-8 | S | Spain: Almeria: Cabo de Gata: Playa Genoveses | 36.746 | -2.122 |
| *B. hybridum* | Bhyb17 | Bhyb_Bhyb17 | S | Iran: Kohkiluyeh, Yasooj. MRR:B98-H137 | 30.654 | 51.589 |
| *B. hybridum* | Bhyb26 | Bhyb_Bhyb26_reference | D | Spain: Jaen,La Cimbarra | 38.401 | -3.39 |
| *B. hybridum* | Bhyb26_2 | Bhyb_Bhyb26_2 | S | Spain: Jaen,La Cimbarra | 38.401 | -3.39 |
| *B. hybridum* | Bhyb50 | Bhyb_Bhyb50 | S | Israel: Petah Tikwa. AB&SE BRA143 | 32.109 | 34.885 |
| *B. hybridum* | Bhyb51 | Bhyb_Bhyb51 | S | Israel: Dorot. AB&SE BRA144 | 31.51 | 34.642 |
| *B. hybridum* | BhybCA_UCM | Bhyb_BhybCA_UCM | D | Spain: Madrid, El Molar | 40.734 | -3.58 |
| *B. hybridum* | BhybEM_UCM | Bhyb_BhybEM_UCM | D | Spain: Badajoz, Castuera | 38.726 | -5.541 |
| *B. hybridum* | Bra117 | Bhyb_Bra117 | S | Israel: Ba'al Hazor. AB&SE BRA117 | 33.018 | 35.567 |
| *B. hybridum* | Bra142 | Bhyb_Bra142 | S | Israel: Nahal Karkom. AB&SE BRA142 | 30.321 | 34.716 |
| *B. hybridum* | Bra143 | Bhyb_Bra143 | S | Israel: Petah Tikwa. AB&SE BRA143 | 32.109 | 34.885 |
| *B. hybridum* | Bra151 | Bhyb_Bra151 | S | Israel: Keshet Cave. AB&SE BRA151 | 33.077 | 35.195 |
| *B. hybridum* | Bra293 | Bhyb_Bra293 | S | Israel: Sion. AB&SE BRA293 | 31.771 | 35.228 |
| *B. hybridum* | Bra32 | Bhyb_Bra32 | S | Israel: Carmel. AB&SE BRA32 | 32.917 | 35.288 |
| *B. hybridum* | Bra9 | Bhyb_Bra9 | S | Israel: Ein Fesh'ha. AB&SE BRA9 | 31.725 | 35.457 |
| *B. hybridum* | Bhyb6 | Bhyb_Bhyb6 | S | Iran: Fars, Kazeroon, Davan. MRR B100-H141 | 29.703 | 51.676 |
| *B. hybridum* | Bhyb7 | Bhyb_Bhyb7 | S | Iran: 25 Km to Ramhorm from Ahwaz. MRR BdisIran2 | 31.283 | 49.574 |
| *B. hybridum* | Bhyb8 | Bhyb_Bhyb8 | S | Iran: Asalooeyeh to Booshehr MRR BdisIran1 | 28.164 | 51.719 |
| *B. hybridum* | CIW1_1 | Bhyb_CIW1_1 | S | Turkey | 38.964 | 35.243 |
| *B. hybridum* | DRAGO881 | Bhyb_DRAGO881 | S | Spain: Mallorca, Sa Dragonera, Cova des Moro | 39.926 | 3.912 |
| *B. hybridum* | ESPU | Bhyb_ESPU | S | Spain: Murcia, Sierra Espuña | 37.936 | -1.542 |
| *B. hybridum* | FARO | Bhyb_FARO | S | Portugal: Algarve, Faro | 37.019 | -7.93 |
| *B. hybridum* | GROSE | Bhyb_GROSE | S | Spain: Girona, Roses, Castell de la Trinitat | 42.258 | 3.161 |
| *B. hybridum* | IBD_107 | Bhyb_IBD_107 | S | Israel | 31.046 | 34.852 |
| *B. hybridum* | IBD_189 | Bhyb_IBD_189 | S | Israel | 31.046 | 34.852 |
| *B. hybridum* | Isk_P1 | Bhyb_Isk_P1 | S | Turkey | 38.964 | 35.243 |
| *B. hybridum* | LLIG | Bhyb_LLIG | S | Spain: Girona, Cadaqués, Port Lligat | 42.291 | 3.277 |
| *B. hybridum* | MT110_4 | Bhyb_MT110_4 | S | Turkey | 38.964 | 35.243 |
| *B. hybridum* | MT118_1 | Bhyb_MT118_1 | S | Turkey | 38.964 | 35.243 |
| *B. hybridum* | PC1_17 | Bhyb_PC1_17 | S | Morocco: Tetuan | 35.57 | -5.368 |
| *B. hybridum* | PC31_17 | Bhyb_PC31_17 | S | Morocco: Djbel Tazzaot | 34.217 | -4.529 |
| *B. hybridum* | PC63_17 | Bhyb_PC63_17 | S | Morocco: Xauen | 35.172 | -5.269 |
| *B. hybridum* | PC64_17 | Bhyb_PC64_17 | S | Morocco: Beni Oulla | 34.657 | -3.498 |
| *B. hybridum* | PC66_17 | Bhyb_PC66_17 | S | Morocco: Tazza-Bou Idir | 34.233 | -4.135 |
| *B. hybridum* | PC76_17 | Bhyb_PC76_17 | S | Morocco: Ifrane National Park, Azrou | 33.443 | -5.165 |
| *B. hybridum* | PC85_17 | Bhyb_PC85_17 | S | Morocco: Mulay-Idriss | 34.064 | -5.337 |
| *B. hybridum* | PC89_17 | Bhyb_PC89_17 | S | Morocco: Larache | 35.178 | -6.155 |
| *B. hybridum* | PC90_17 | Bhyb_PC90_17 | S | Morocco: Tanger | 35.76 | -5.833 |
| *B. hybridum* | PC92_17 | Bhyb_PC92_17 | S | Morocco: Hassana | 34.19 | -6.344 |
| *B. hybridum* | POBL | Bhyb_POBL | S | Spain: Tarragona, Poble Nou del Delta | 40.735 | 0.805 |
| *B. hybridum* | PRAT | Bhyb_PRAT | S | Spain: Barcelona, El Prat Del Llobregat | 41.297 | 2.074 |
| *B. hybridum* | SORB | Bhyb_SORB | S | Spain: Almería, Sorbas | 37.048 | -2.161 |
| *B. hybridum* | TISC | Bhyb_TISC | S | Spain: Jaén, Quesada, Sierra Cazorla | 37.85 | -3.002 |
| *B. hybridum* | TORO | Bhyb_TORO | S | Spain: Mallorca, El Toro | 39.523 | 2.491 |
| *B. hybridum* | Bhyb_ECI | Bhyb_ECI_reference | S | Israel: Evolution Canyon I | 32.71 | 34.97 |
| *B. hybridum* | Bhyb-AS-1 | Bhyb-AS-1 | S | Israel: Evolution Canyon I | 32.71 | 34.97 |
| *B. hybridum* | Bhyb-AS-2 | Bhyb-AS-2 | S | Israel: Evolution Canyon I | 32.71 | 34.97 |
| *B. hybridum* | Bhyb_AS-5 | Bhyb_AS-5 | S | Israel: Evolution Canyon I | 32.71 | 34.97 |
| *B. hybridum* | Bhyb_AS-6 | Bhyb_AS-6 | S | Israel: Evolution Canyon I | 32.71 | 34.97 |
| *B. hybridum* | Bhyb-AS-10 | Bhyb-AS-10 | S | Israel: Evolution Canyon I | 32.71 | 34.97 |
| *B. hybridum* | Bhyb-AS-11 | Bhyb-AS-11 | S | Israel: Evolution Canyon I | 32.71 | 34.97 |
| *B. hybridum* | Bhyb-AS-12 | Bhyb-AS-12 | S | Israel: Evolution Canyon I | 32.71 | 34.97 |
| *B. hybridum* | Bhyb-7 | Bhyb-7 | S | Israel: Lehavim | 31.369 | 34.802 |

**Table S2.** Summary statistics based on adapter-free and contamination-filtered resequencing data of the studied *Brachypodium* accessions and their SNP mapped against the concatenated reference genomes of the diploid progenitor species *B. distachyon* (Bd21 v3_2; https://phytozome-next.jgi.doe.gov/info/Bdistachyon_v3_2) and *B. stacei* (ABR114 v2_1; https://phytozome-next.jgi.doe.gov/info/Bstacei_v2_1). The table indicates the species, accession code, total number of mapped reads (ReadsMapped), percentage of mapped reads to *B. distachyon* (%MappedBdis) and *B. stacei* (%MappedBsta), genome coverage for both species (BdisCoverage and BstaCoverage), total number of positions in the raw alignment (Positions), and the number of identified syntenic SNPs (identified unique homeologous polymorphisms (SHP, equivalent to syntenic SNPs) using the AlloSHP script by Sancho et al. (2025) per accession.

| Species | Accession | ReadsMapped | Depth | %<br>MappedBdis | % MappedBsta | BdisCoverage | BstaCoverage | Positions | SyntenicSNPs |
| --- | --- | --- | --- | --- | --- | --- | --- | --- | --- |
| *Oryza sativa* | Oryza_sativa | 8127292 |  | 56.07% | 43.93% | 3.78% | 3.80% | 12331508 | 367184 |
| *B. distachyon* | Bdis_ABR2 | 349998595 | 113.81 | 97.89% | 2.11% | 98.18% | 1.02% | 267047387 | 7873347 |
| *B. distachyon* | Bdis_ABR3 | 332755977 | 78.96 | 97.82% | 2.18% | 98.17% | 0.95% | 266883731 | 7870958 |
| *B. distachyon* | Bdis_ABR4 | 349998595 | 113.81 | 97.89% | 2.11% | 98.18% | 1.02% | 267047387 | 7873347 |
| *B. distachyon* | Bdis_ABR5 | 313689354 | 103.16 | 97.62% | 2.38% | 98.17% | 0.94% | 265026391 | 7872551 |
| *B. distachyon* | Bdis_ABR6 | 324325317 | 110.27 | 98.01% | 1.99% | 97.95% | 1.20% | 264130428 | 7848656 |
| *B. distachyon* | Bdis_ABR7 | 379440897 | 120.48 | 97.00% | 3.00% | 98.23% | 1.04% | 266973388 | 7840691 |
| *B. distachyon* | Bdis_ABR9 | 239194070 | 6.11 | 96.21% | 3.79% | 30.43% | 0.23% | 226811384 | 2310630 |
| *B. distachyon* | Bdis_Adi_10 | 178275426 | 57.68 | 96.87% | 3.13% | 98.21% | 0.78% | 265894813 | 7878126 |
| *B. distachyon* | Bdis_Adi_12 | 89898077 | 62.77 | 97.14% | 2.86% | 98.28% | 0.76% | 262978817 | 7889277 |
| *B. distachyon* | Bdis_Adi_15 | 194248450 | 50.89 | 97.46% | 2.54% | 98.31% | 0.21% | 261369152 | 7878126 |
| *B. distachyon* | Bdis_Adi_4 | 109893953 | 22.14 | 97.96% | 2.04% | 98.23% | 0.13% | 258706426 | 7890870 |
| *B. distachyon* | Bdis_Adi_9 | 46247439 | 42.76 | 97.84% | 2.16% | 98.33% | 0.18% | 260647370 | 7884498 |
| *B. distachyon* | Bdis_AL1A.1 | 61275878 | 27.06 | 96.70% | 3.30% | 97.88% | 0.23% | 258277889 | 7838301 |
| *B. distachyon* | Bdis_AL2D | 95421145 | 44.37 | 97.48% | 2.52% | 97.95% | 0.33% | 258168533 | 7847859 |
| *B. distachyon* | Bdis_AL2E | 91817238 | 41.94 | 97.23% | 2.77% | 97.95% | 0.31% | 258212163 | 7850249 |
| *B. distachyon* | Bdis_AL2F | 98281567 | 46.01 | 97.42% | 2.58% | 97.96% | 0.32% | 258051703 | 7851045 |
| *B. distachyon* | Bdis_Arm2B | 106024929 | 39.18 | 97.21% | 2.79% | 98.22% | 0.24% | 260784081 | 7878126 |
| *B. distachyon* | Bdis_Arm3A | 103212798 | 50.42 | 98.15% | 1.85% | 98.25% | 0.25% | 260476924 | 7881312 |
| *B. distachyon* | Bdis_Arm3G | 99778948 | 48.39 | 97.75% | 2.25% | 98.25% | 0.24% | 260454528 | 7881312 |
| *B. distachyon* | Bdis_Arm3i | 82018168 | 36.91 | 97.05% | 2.95% | 98.25% | 0.19% | 259826479 | 7881312 |
| *B. distachyon* | Bdis_Arn1 | 386787171 | 120.62 | 97.45% | 2.55% | 98.02% | 1.31% | 264492620 | 7855824 |
| *B. distachyon* | Bdis_Bd1-1 | 162258927 | 121 | 96.82% | 3.18% | 98.01% | 0.46% | 256829388 | 7856621 |
| *B. distachyon* | Bdis_Bd130-1 | 299340836 | 216.16 | 97.76% | 2.24% | 98.22% | 0.41% | 270144186 | 7878923 |
| *B. distachyon* | Bdis_Bd162 | 218023967 | 174.99 | 97.74% | 2.26% | 98.17% | 0.39% | 259143069 | 7873347 |
| *B. distachyon* | Bdis_Bd18-1 | 231124230 | 72.6 | 96.84% | 3.16% | 98.28% | 1.11% | 264229468 | 7883702 |
| *B. distachyon* | Bdis_Bd21_reference | 174129585 | 52.07 | 97.81% | 2.19% | 92.71% | 0.57% | 266651322 | 7463153 |
| *B. distachyon* | Bdis_Bd2-3 | 146938609 | 47.33 | 97.27% | 2.73% | 98.27% | 0.65% | 263496796 | 7884498 |
| *B. distachyon* | Bdis_Bd30-1 | 204252096 | 152.47 | 96.74% | 3.26% | 98.21% | 0.41% | 257620010 | 7878126 |
| *B. distachyon* | Bdis_Bd3-1 | 312552011 | 78.88 | 97.57% | 2.43% | 98.45% | 0.21% | 270727163 | 7860603 |
| *B. distachyon* | Bdis_BdTR10C | 177020621 | 58.05 | 97.06% | 2.94% | 98.21% | 0.69% | 263195924 | 7878923 |
| *B. distachyon* | Bdis_BdTR10D | 103424641 | 50.05 | 97.82% | 2.18% | 98.31% | 0.24% | 261436080 | 7886888 |
| *B. distachyon* | Bdis_BdTR11A | 124070385 | 31.88 | 96.94% | 3.06% | 98.27% | 0.54% | 262459286 | 7883702 |
| *B. distachyon* | Bdis_BdTR11E | 87466671 | 41.19 | 97.67% | 2.33% | 98.33% | 0.17% | 259955484 | 7890870 |
| *B. distachyon* | Bdis_BdTR11G | 158474925 | 50.47 | 97.21% | 2.79% | 98.31% | 0.46% | 262194746 | 7888481 |
| *B. distachyon* | Bdis_BdTR11I | 237872631 | 76.87 | 97.32% | 2.68% | 98.32% | 1.06% | 264398411 | 7890074 |
| *B. distachyon* | Bdis_BdTR13B | 116559146 | 55.73 | 97.62% | 2.38% | 98.34% | 0.22% | 261191223 | 7891667 |
| *B. distachyon* | Bdis_BdTR13C | 116385790 | 38.16 | 97.56% | 2.44% | 98.25% | 0.53% | 262245981 | 7883702 |
| *B. distachyon* | Bdis_BdTR13N | 97844215 | 44.69 | 97.23% | 2.77% | 98.33% | 0.19% | 260415952 | 7890870 |
| *B. distachyon* | Bdis_BdTR1i | 335625851 | 105.32 | 96.95% | 3.05% | 98.28% | 0.75% | 264484067 | 7886091 |
| *B. distachyon* | Bdis_BdTR2B | 279622662 | 81.76 | 96.52% | 3.48% | 98.23% | 0.98% | 265950914 | 7882109 |
| *B. distachyon* | Bdis_BdTR2G | 148304581 | 43.98 | 96.10% | 3.90% | 98.20% | 0.63% | 262243051 | 7877330 |
| *B. distachyon* | Bdis_BdTR3C | 142525659 | 45.02 | 96.68% | 3.32% | 98.12% | 0.69% | 263421507 | 7870958 |
| *B. distachyon* | Bdis_BdTR3M | 114732129 | 52.6 | 97.44% | 2.56% | 98.32% | 0.22% | 260724838 | 7888481 |
| *B. distachyon* | Bdis_BdTR5A | 102960002 | 47.19 | 97.46% | 2.54% | 98.33% | 0.19% | 260689430 | 7891667 |
| *B. distachyon* | Bdis_BdTR5I | 117389793 | 35.02 | 96.39% | 3.61% | 98.24% | 0.51% | 261905955 | 7882905 |
| *B. distachyon* | Bdis_BdTR7a | 246880868 | 78.55 | 97.47% | 2.53% | 97.94% | 0.89% | 263804489 | 7847859 |
| *B. distachyon* | Bdis_BdTR8i | 320711251 | 100.24 | 95.66% | 4.34% | 97.95% | 6.16% | 275331792 | 7846266 |
| *B. distachyon* | Bdis_BdTR9K | 93173084 | 28.16 | 96.93% | 3.07% | 98.22% | 0.40% | 261301441 | 7878923 |
| *B. distachyon* | Bdis_BdTR9M | 85237028 | 40.82 | 98.07% | 1.93% | 98.33% | 0.19% | 260250905 | 7889277 |
| *B. distachyon* | Bdis_Bis1 | 44596332 | 14.19 | 97.26% | 2.74% | 97.54% | 0.12% | 259841628 | 7823964 |
| *B. distachyon* | Bdis_BNT3 | 85022874 | 47.37 | 98.30% | 1.70% | 97.97% | 1.60% | 262240434 | 7850249 |
| *B. distachyon* | Bdis_BNT4.1 | 82714964 | 38.83 | 97.37% | 2.63% | 97.95% | 0.32% | 258012969 | 7847063 |
| *B. distachyon* | Bdis_BNT8 | 119877810 | 61.23 | 97.37% | 2.63% | 98.00% | 0.41% | 259732565 | 7854231 |
| *B. distachyon* | Bdis_CREAF2 | 415817064 | 184.02 | 97.48% | 2.52% | 98.21% | 0.85% | 356761909 | 7877330 |
| *B. distachyon* | Bdis_CSR-6 | 115116867 | 52.17 | 97.11% | 2.89% | 97.82% | 0.46% | 278743908 | 7833522 |
| *B. distachyon* | Bdis_CSR-7 | 80104784 | 35.49 | 95.97% | 4.03% | 97.76% | 0.43% | 350942429 | 7825557 |
| *B. distachyon* | Bdis_CU1.6-1 | 92210867 | 43.92 | 97.90% | 2.10% | 98.18% | 0.28% | 259105975 | 7874144 |
| *B. distachyon* | Bdis_CU5.5 | 79756022 | 40.03 | 98.06% | 1.94% | 98.18% | 0.24% | 258949305 | 7874144 |
| *B. distachyon* | Bdis_Foz1 | 304381599 | 94.71 | 97.17% | 2.83% | 98.18% | 0.96% | 318132801 | 7870958 |
| *B. distachyon* | Bdis_G30i2 | 85343853 | 38.06 | 96.99% | 3.01% | 98.01% | 0.27% | 258174522 | 7855824 |
| *B. distachyon* | Bdis_G31i4 | 88118036 | 42.79 | 98.02% | 1.98% | 98.26% | 0.19% | 260053388 | 7882905 |
| *B. distachyon* | Bdis_G32i2 | 80935093 | 37.89 | 97.42% | 2.58% | 98.26% | 0.18% | 259621739 | 7882109 |
| *B. distachyon* | Bdis_G33i4 | 85116419 | 41.29 | 96.85% | 3.15% | 97.98% | 0.28% | 258992000 | 7851842 |
| *B. distachyon* | Bdis_G33i6 | 90188914 | 41.95 | 97.44% | 2.56% | 97.99% | 0.29% | 258868911 | 7853435 |
| *B. distachyon* | Bdis_G34i2 | 106565166 | 48.11 | 97.31% | 2.69% | 98.28% | 0.23% | 260547533 | 7883702 |
| *B. distachyon* | Bdis_G34i6 | 85297837 | 40.12 | 97.63% | 2.37% | 98.27% | 0.20% | 259867872 | 7882905 |
| *B. distachyon* | Bdis_Gaz-1 | 93613510 | 44.97 | 97.66% | 2.34% | 98.32% | 0.24% | 260026784 | 7889277 |
| *B. distachyon* | Bdis_Gaz-2 | 135389140 | 65.6 | 97.92% | 2.08% | 98.33% | 0.27% | 261451502 | 7890870 |
| *B. distachyon* | Bdis_Gaz-7 | 89235664 | 32.42 | 97.27% | 2.73% | 98.30% | 0.19% | 259907118 | 7887684 |
| *B. distachyon* | Bdis_Gaz-8 | 172867611 | 54.8 | 97.04% | 2.96% | 98.01% | 0.80% | 263983336 | 7864586 |
| *B. distachyon* | Bdis_GR6.4 | 67822394 | 32.72 | 97.70% | 2.30% | 97.94% | 0.28% | 257742287 | 7848656 |
| *B. distachyon* | Bdis_Graz | 143871869 | 69.63 | 97.55% | 2.45% | 98.14% | 0.40% | 283537482 | 7869365 |
| *B. distachyon* | Bdis_GU5-1 | 107056943 | 57.4 | 97.91% | 2.09% | 98.18% | 0.30% | 259960331 | 7873347 |
| *B. distachyon* | Bdis_HU3.4 | 100060343 | 46.75 | 97.50% | 2.50% | 98.15% | 0.31% | 259044708 | 7869365 |
| *B. distachyon* | Bdis_J4.3.1 | 78911736 | 36.37 | 97.27% | 2.73% | 98.14% | 0.23% | 258203459 | 7868568 |
| *B. distachyon* | Bdis_J6.2 | 84575735 | 40.86 | 97.87% | 2.13% | 98.15% | 0.26% | 258622592 | 7870161 |
| *B. distachyon* | Bdis_Jer1 | 340179489 | 104.9 | 97.58% | 2.42% | 98.17% | 1.11% | 266001087 | 7872551 |
| *B. distachyon* | Bdis_Kah1 | 62223864 | 19.28 | 97.21% | 2.79% | 97.89% | 0.22% | 261336294 | 7849452 |
| *B. distachyon* | Bdis_Kah6 | 92743275 | 44.19 | 97.64% | 2.36% | 98.30% | 0.21% | 260139434 | 7886888 |
| *B. distachyon* | Bdis_Koz-1 | 269127474 | 84.39 | 96.85% | 3.15% | 98.31% | 0.72% | 263566572 | 7887684 |
| *B. distachyon* | Bdis_Koz-2 | 92959395 | 42.91 | 96.98% | 3.02% | 98.30% | 0.19% | 260112673 | 7886091 |
| *B. distachyon* | Bdis_Koz3 | 326186044 | 18.31 | 97.72% | 2.28% | 90.89% | 0.34% | 261479623 | 7291110 |
| *B. distachyon* | Bdis_Koz-5 | 104201845 | 47.68 | 97.23% | 2.77% | 98.33% | 0.20% | 260935100 | 7891667 |
| *B. distachyon* | Bdis_LPA3.2 | 108353788 | 44.92 | 96.84% | 3.16% | 97.81% | 0.40% | 257515011 | 7832726 |
| *B. distachyon* | Bdis_Luc1 | 291934369 | 92.71 | 97.40% | 2.60% | 98.20% | 0.84% | 263778255 | 7875737 |
| *B. distachyon* | Bdis_Mig3 | 385504049 | 114.49 | 97.13% | 2.87% | 98.23% | 1.08% | 268197080 | 7816796 |
| *B. distachyon* | Bdis_Mon3 | 286658828 | 90.3 | 97.43% | 2.57% | 97.97% | 1.09% | 289263229 | 7851045 |
| *B. distachyon* | Bdis_Mur1 | 342284154 | 103.5 | 97.48% | 2.52% | 98.18% | 1.05% | 266968943 | 7872551 |
| *B. distachyon* | Bdis_Nvk1 | 118549735 | 61.61 | 97.65% | 2.35% | 98.04% | 0.38% | 258567281 | 7859010 |
| *B. distachyon* | Bdis_PC10_17 | 103496710 | 51 | 97.48% | 2.52% | 98.05% | 0.38% | 265893727 | 7859010 |
| *B. distachyon* | Bdis_PC95_17 | 269148585 | 122.24 | 97.61% | 2.39% | 98.15% | 0.60% | 321571052 | 7870958 |
| *B. distachyon* | Bdis_Per1 | 241478008 | 76.39 | 97.26% | 2.74% | 98.24% | 0.71% | 265305400 | 7875737 |
| *B. distachyon* | Bdis_S11A.3 | 72235175 | 33.82 | 97.06% | 2.94% | 98.18% | 0.23% | 258587902 | 7873347 |
| *B. distachyon* | Bdis_S11B-2 | 86941344 | 39.97 | 97.04% | 2.96% | 98.20% | 0.28% | 283116059 | 7874940 |
| *B. distachyon* | Bdis_S11D-2 | 70861270 | 35.34 | 97.99% | 2.01% | 98.19% | 0.25% | 258619828 | 7873347 |
| *B. distachyon* | Bdis_S12B-4 | 75757868 | 36.24 | 97.97% | 2.03% | 98.16% | 0.25% | 258534888 | 7870161 |
| *B. distachyon* | Bdis_S16D | 80726296 | 29.83 | 97.16% | 2.84% | 98.17% | 0.25% | 258437817 | 7871754 |
| *B. distachyon* | Bdis_S17D-1 | 87405060 | 41.37 | 97.73% | 2.27% | 98.20% | 0.28% | 258812554 | 7874940 |
| *B. distachyon* | Bdis_S19A | 85398589 | 37.4 | 96.66% | 3.34% | 98.13% | 0.26% | 258480895 | 7866975 |
| *B. distachyon* | Bdis_S22B-1 | 55096887 | 25.94 | 97.17% | 2.83% | 98.13% | 0.18% | 257693209 | 7869365 |
| *B. distachyon* | Bdis_S22C.1 | 134463413 | 64.23 | 97.58% | 2.42% | 98.22% | 0.37% | 260367002 | 7876533 |
| *B. distachyon* | Bdis_S6B-3 | 98276003 | 45.63 | 97.27% | 2.73% | 98.19% | 0.29% | 280126665 | 7874144 |
| *B. distachyon* | Bdis_S6D-5 | 79462072 | 38.15 | 97.73% | 2.27% | 97.94% | 0.31% | 258734843 | 7847063 |
| *B. distachyon* | Bdis_S6E | 104126469 | 48.27 | 96.75% | 3.25% | 98.18% | 0.29% | 258834759 | 7874144 |
| *B. distachyon* | Bdis_S8iiC | 392878016 | 116.87 | 97.07% | 2.93% | 98.22% | 1.08% | 265430933 | 7877330 |
| *B. distachyon* | Bdis_Sg2.1.1 | 91419587 | 42.24 | 97.37% | 2.63% | 98.18% | 0.30% | 259309736 | 7872551 |
| *B. distachyon* | Bdis_SLZ2.1 | 100034584 | 49.45 | 98.08% | 1.92% | 97.99% | 0.36% | 258319791 | 7853435 |
| *B. distachyon* | Bdis_Sou-AB1.4-2 | 89694631 | 40.7 | 97.37% | 2.63% | 98.16% | 0.23% | 258749535 | 7870958 |
| *B. distachyon* | Bdis_Sou-AB3.3 | 76840239 | 42.03 | 98.24% | 1.76% | 98.09% | 0.27% | 258078127 | 7862993 |
| *B. distachyon* | Bdis_Tek-2 | 86500191 | 27.35 | 96.95% | 3.05% | 97.83% | 0.45% | 260881983 | 7827150 |
| *B. distachyon* | Bdis_Tek-9 | 84220221 | 37.99 | 96.85% | 3.15% | 97.94% | 0.31% | 258084382 | 7847063 |
| *B. distachyon* | Bdis_Uni2 | 389650989 | 125.03 | 97.12% | 2.88% | 98.24% | 1.25% | 267474972 | 7812017 |
| *B. distachyon* | Bdis_Z3.6 | 83010595 | 42.67 | 97.95% | 2.05% | 98.16% | 0.28% | 258343110 | 7870161 |
| *B. stacei* | Bsta_11_19_1_T | 143547328 | 69.22 | 6.51% | 93.49% | 0.34% | 99.69% | 251509389 | 7927509 |
| *B. stacei* | Bsta_ABR114_reference | 291560374 | 52.21 | 3.66% | 96.34% | 0.68% | 91.70% | 252977301 | 7315005 |
| *B. stacei* | Bsta_Alsur1 | 115349849 | 54.39 | 4.43% | 95.57% | 0.64% | 99.58% | 251019230 | 7916358 |
| *B. stacei* | Bsta_AS-18 | 63024975 | 32.75 | 4.14% | 95.86% | 0.23% | 99.62% | 247524352 | 7918747 |
| *B. stacei* | Bsta_AS-19 | 57129499 | 30.71 | 3.05% | 96.95% | 0.20% | 99.61% | 246035644 | 7917154 |
| *B. stacei* | Bsta_AS-20 | 56489572 | 28.91 | 3.68% | 96.32% | 0.21% | 99.62% | 245528098 | 7917951 |
| *B. stacei* | Bsta_AS-21 | 51257017 | 31.96 | 2.65% | 97.35% | 0.19% | 99.56% | 246016825 | 7912375 |
| *B. stacei* | Bsta_AS-22 | 61109696 | 31.05 | 3.31% | 96.69% | 0.24% | 99.63% | 245535266 | 7918747 |
| *B. stacei* | Bsta_AS-23 | 59417704 | 32.24 | 3.69% | 96.31% | 0.22% | 99.61% | 246310750 | 7917154 |
| *B. stacei* | Bsta_AS-24 | 61279148 | 27.85 | 3.38% | 96.62% | 0.20% | 99.61% | 245670518 | 7917154 |
| *B. stacei* | Bsta_AS-25 | 52055234 | 66.03 | 3.06% | 96.94% | 0.20% | 99.60% | 245884765 | 7916358 |
| *B. stacei* | Bsta_AS-26 | 53585848 | 27.67 | 3.52% | 96.48% | 0.22% | 99.60% | 245326483 | 7917154 |
| *B. stacei* | Bsta_AS-27 | 48928780 | 68.25 | 3.17% | 96.83% | 0.20% | 99.58% | 245959643 | 7913968 |
| *B. stacei* | Bsta_AS-28 | 54315589 | 44.36 | 6.19% | 93.81% | 0.23% | 99.61% | 245705602 | 7916358 |
| *B. stacei* | Bsta_AS-29 | 48449957 | 31.01 | 4.06% | 95.94% | 0.21% | 99.59% | 249865109 | 7915561 |
| *B. stacei* | Bsta_AS-30 | 51231451 | 28.76 | 3.88% | 96.12% | 0.19% | 99.58% | 250968253 | 7914765 |
| *B. stacei* | Bsta_AS-31 | 47216284 | 29.31 | 3.91% | 96.09% | 0.18% | 99.56% | 250181280 | 7912375 |
| *B. stacei* | Bsta_AS-6 | 134714156 | 32.37 | 3.99% | 96.01% | 0.30% | 99.66% | 248937431 | 7922730 |
| *B. stacei* | Bsta_BG15 | 439706105 | 189.07 | 8.79% | 91.21% | 0.61% | 99.72% | 307736045 | 7931491 |
| *B. stacei* | Bsta_BG19 | 154684415 | 73.6 | 5.28% | 94.72% | 0.26% | 99.68% | 249365858 | 7928305 |
| *B. stacei* | Bsta_BG26 | 162571504 | 73.27 | 6.59% | 93.41% | 0.31% | 99.66% | 273727173 | 7925119 |
| *B. stacei* | Bsta_BG31 | 159607852 | 78.11 | 5.42% | 94.58% | 0.32% | 99.68% | 247300289 | 7925916 |
| *B. stacei* | Bsta_BG7 | 158943207 | 77.34 | 6.66% | 93.34% | 0.40% | 99.66% | 248957039 | 7924323 |
| *B. stacei* | Bsta_BJAE2 | 162096613 | 85.91 | 3.70% | 96.30% | 0.32% | 99.73% | 250328240 | 7932288 |
| *B. stacei* | Bsta_BONA | 157163161 | 82.1 | 3.91% | 96.09% | 0.34% | 99.67% | 250077416 | 7924323 |
| *B. stacei* | Bsta_Br-135 | 129984888 | 64.06 | 5.56% | 94.44% | 0.43% | 99.63% | 248909639 | 7921137 |
| *B. stacei* | Bsta_Bra223 | 111442774 | 57.45 | 3.08% | 96.92% | 0.36% | 99.65% | 251436361 | 7923526 |
| *B. stacei* | Bsta_Bra271 | 134687786 | 67.37 | 4.83% | 95.17% | 0.32% | 99.67% | 247298980 | 7924323 |
| *B. stacei* | Bsta_Bra89 | 143854119 | 72.13 | 4.20% | 95.80% | 0.33% | 99.67% | 247893142 | 7925119 |
| *B. stacei* | Bsta_CAMPA | 169664437 | 86.55 | 3.60% | 96.40% | 0.35% | 99.69% | 252146598 | 7927509 |
| *B. stacei* | Bsta_Cef2 | 58428819 | 18.32 | 7.38% | 92.62% | 0.16% | 99.13% | 257514210 | 7876533 |
| *B. stacei* | Bsta_DRAGO891 | 133447807 | 69.21 | 4.38% | 95.62% | 0.33% | 99.67% | 249483486 | 7925916 |
| *B. stacei* | Bsta_ECI_reference | 77261395 | 72.04 | 5.60% | 94.40% | 0.55% | 99.69% | 253723955 | 7927509 |
| *B. stacei* | Bsta_ES-12 | 29271120 | 61.89 | 2.65% | 97.35% | 0.14% | 98.93% | 244739875 | 7857417 |
| *B. stacei* | Bsta_ES-13 | 49361407 | 38.92 | 4.31% | 95.69% | 0.22% | 99.53% | 247211708 | 7909189 |
| *B. stacei* | Bsta_ES-14 | 53484014 | 24 | 4.30% | 95.70% | 0.22% | 99.55% | 250386321 | 7911579 |
| *B. stacei* | Bsta_ES-15 | 57914334 | 22.1 | 4.30% | 95.70% | 0.23% | 99.57% | 250423976 | 7913172 |
| *B. stacei* | Bsta_ES-16 | 56078269 | 22.1 | 3.55% | 96.45% | 0.21% | 99.58% | 246005485 | 7914765 |
| *B. stacei* | Bsta_ES-17 | 48715507 | 19.67 | 4.55% | 95.45% | 0.19% | 99.45% | 243578799 | 7902021 |
| *B. stacei* | Bsta_ES-18 | 38920949 | 12.84 | 3.96% | 96.04% | 0.17% | 99.42% | 244491569 | 7900428 |
| *B. stacei* | Bsta_ES-19 | 24834464 | 20.21 | 4.41% | 95.59% | 0.12% | 96.63% | 245752299 | 7663869 |
| *B. stacei* | Bsta_ES-2 | 42156909 | 66.37 | 4.35% | 95.65% | 0.70% | 99.54% | 246069732 | 7882905 |
| *B. stacei* | Bsta_ES-20 | 38394023 | 25.12 | 4.32% | 95.68% | 0.16% | 99.32% | 246176375 | 7889277 |
| *B. stacei* | Bsta_ES-21 | 45665772 | 26.76 | 4.31% | 95.69% | 0.20% | 99.47% | 247567693 | 7865382 |
| *B. stacei* | Bsta_ES-22 | 42163542 | 29.11 | 2.98% | 97.02% | 0.72% | 99.54% | 245859147 | 7882905 |
| *B. stacei* | Bsta_ES-23 | 46930941 | 29.04 | 3.60% | 96.40% | 0.19% | 99.52% | 245308649 | 7909189 |
| *B. stacei* | Bsta_ES-24 | 23123023 | 24.93 | 2.77% | 97.23% | 0.11% | 95.26% | 243438738 | 7549970 |
| *B. stacei* | Bsta_ES-25 | 34169552 | 23.31 | 3.35% | 96.65% | 0.17% | 99.26% | 245276582 | 7884498 |
| *B. stacei* | Bsta_FELEN | 183430246 | 90.85 | 5.25% | 94.75% | 0.40% | 99.69% | 257508986 | 7928305 |
| *B. stacei* | Bsta_Hada | 148029780 | 75.79 | 4.70% | 95.30% | 0.35% | 99.67% | 250587624 | 7924323 |
| *B. stacei* | Bsta_IBD_10 | 278543912 | 124 | 7.04% | 92.96% | 0.45% | 99.69% | 249058905 | 7927509 |
| *B. stacei* | Bsta_Larva3 | 273966594 | 134.68 | 4.40% | 95.60% | 0.54% | 99.66% | 252135007 | 7923526 |
| *B. stacei* | Bsta_LP6_1 | 365229078 | 167.53 | 6.69% | 93.31% | 0.55% | 99.63% | 250047495 | 7921137 |
| *B. stacei* | Bsta_MONSUL | 146151789 | 73.06 | 4.60% | 95.40% | 0.28% | 99.70% | 257018419 | 7929898 |
| *B. stacei* | Bsta_Ourich | 176617482 | 94.08 | 3.41% | 96.59% | 0.38% | 99.64% | 250130281 | 7921933 |
| *B. stacei* | Bsta_PC23_17 | 95710082 | 45.89 | 6.35% | 93.65% | 0.32% | 99.60% | 247862798 | 7916358 |
| *B. stacei* | Bsta_PC47_17 | 377321874 | 173.66 | 7.11% | 92.89% | 1.35% | 99.70% | 396700319 | 7929898 |
| *B. stacei* | Bsta_PC49_17 | 273099366 | 134.25 | 4.51% | 95.49% | 0.45% | 99.69% | 258015748 | 7928305 |
| *B. stacei* | Bsta_QVL1 | 115626766 | 58.26 | 5.08% | 94.92% | 0.32% | 99.68% | 251839986 | 7926712 |
| *B. stacei* | Bsta_TE4.3 | 65467241 | 98.76 | 7.35% | 92.65% | 0.29% | 99.50% | 243661748 | 7907596 |
| *B. stacei* | Bsta_VMO1 | 124439040 | 65.01 | 3.40% | 96.60% | 0.35% | 99.72% | 248835218 | 7932288 |
| *B. stacei* | Bsta-1 | 103380585 | 45.28 | 7.84% | 92.16% | 0.46% | 99.58% | 251659949 | 7915561 |
| *B. stacei* | Bsta2 | 110374823 | 55.49 | 5.24% | 94.76% | 0.39% | 99.62% | 254466658 | 7919544 |
| *B. stacei* | Bsta4 | 150560735 | 75.82 | 4.76% | 95.24% | 0.36% | 99.64% | 247847586 | 7921933 |
| *B. stacei* | Bsta5 | 435275946 | 197.17 | 6.37% | 93.63% | 0.59% | 99.70% | 251248612 | 7929898 |
| *B. hybridum* | Bhyb_10_11_2_I | 500154405 | 133.86 | 51.68% | 48.32% | 97.95% | 99.65% | 501221804 | 15770589 |
| *B. hybridum* | Bhyb_10_19_5_I | 263320207 | 67.19 | 51.61% | 48.39% | 97.86% | 99.62% | 498587722 | 15758642 |
| *B. hybridum* | Bhyb_11_8_T | 148128379 | 38.53 | 51.64% | 48.36% | 97.81% | 99.54% | 496359631 | 15746694 |
| *B. hybridum* | Bhyb_12_23_2_I | 196184289 | 47.77 | 52.43% | 47.57% | 97.90% | 99.62% | 497235979 | 15757049 |
| *B. hybridum* | Bhyb_12_5_3_I | 260100315 | 65.95 | 51.59% | 48.41% | 97.90% | 99.62% | 498723572 | 15758642 |
| *B. hybridum* | Bhyb_13_19_2_I | 306192431 | 77.31 | 51.39% | 48.61% | 97.98% | 99.70% | 499928882 | 15777758 |
| *B. hybridum* | Bhyb_13_3_5_I | 214442601 | 55.79 | 51.81% | 48.19% | 97.89% | 99.59% | 498237656 | 15755456 |
| *B. hybridum* | Bhyb_14_2_2_I | 333472820 | 86.49 | 51.94% | 48.06% | 97.89% | 99.60% | 498714344 | 15754659 |
| *B. hybridum* | Bhyb_16_2_5_I | 306202207 | 79.06 | 51.78% | 48.22% | 97.90% | 99.62% | 499006474 | 15761828 |
| *B. hybridum* | Bhyb_18_19_T | 180060719 | 67.6 | 51.77% | 48.23% | 97.91% | 99.57% | 498842852 | 15761031 |
| *B. hybridum* | Bhyb_18_6_1_I | 265540946 | 45.79 | 51.84% | 48.16% | 97.85% | 99.61% | 499405781 | 15753863 |
| *B. hybridum* | Bhyb_19_16_5_I | 281975182 | 54.48 | 51.80% | 48.20% | 97.90% | 99.63% | 498631196 | 15763421 |
| *B. hybridum* | Bhyb_19_6_2_T | 215351178 | 71.7 | 51.67% | 48.33% | 97.89% | 99.56% | 497503015 | 15755456 |
| *B. hybridum* | Bhyb_2_2_2_I | 251844880 | 51.26 | 52.06% | 47.94% | 97.92% | 99.67% | 499581562 | 15768996 |
| *B. hybridum* | Bhyb_2_6_1_I | 215843154 | 65.92 | 51.66% | 48.34% | 97.92% | 99.67% | 500958930 | 15764217 |
| *B. hybridum* | Bhyb_20_15_T | 195819889 | 60.78 | 51.51% | 48.49% | 97.87% | 99.57% | 497314497 | 15752270 |
| *B. hybridum* | Bhyb_24_1_I | 243540751 | 54.55 | 51.60% | 48.40% | 98.03% | 99.68% | 502451471 | 15664655 |
| *B. hybridum* | Bhyb_28_28_I | 266570094 | 68.39 | 51.69% | 48.31% | 97.93% | 99.59% | 498981157 | 15761828 |
| *B. hybridum* | Bhyb_3_4_2_I | 384471571 | 99.8 | 50.98% | 49.02% | 97.98% | 99.62% | 500394378 | 15772979 |
| *B. hybridum* | Bhyb_3_4_2_T | 595466639 | 153.78 | 51.35% | 48.65% | 97.93% | 99.64% | 500072785 | 15768996 |
| *B. hybridum* | Bhyb_3_7_2_T | 202439398 | 51.04 | 51.00% | 49.00% | 97.85% | 99.60% | 497002408 | 15753066 |
| *B. hybridum* | Bhyb_7_19_2_I | 143278780 | 36.7 | 52.14% | 47.86% | 97.78% | 99.43% | 495747453 | 15725189 |
| *B. hybridum* | Bhyb_7_27_1_I | 263902300 | 68.88 | 52.42% | 47.58% | 97.94% | 99.61% | 498701806 | 15761031 |
| *B. hybridum* | Bhyb_9_1_3_I | 210562303 | 53.54 | 51.44% | 48.56% | 97.92% | 99.61% | 498708906 | 15761828 |
| *B. hybridum* | Bhyb_9_7_6_I | 294936645 | 74.57 | 51.90% | 48.10% | 97.85% | 99.60% | 499059078 | 15753863 |
| *B. hybridum* | Bhyb_ABR100 | 262389983 | 56.2 | 51.84% | 48.16% | 97.81% | 99.50% | 497055550 | 15740322 |
| *B. hybridum* | Bhyb_ABR101 | 271444815 | 69.35 | 51.53% | 48.47% | 97.93% | 99.58% | 498934023 | 15757049 |
| *B. hybridum* | Bhyb_ABR103 | 246855817 | 62.34 | 51.54% | 48.46% | 97.80% | 99.33% | 495240478 | 15718817 |
| *B. hybridum* | Bhyb_ABR105 | 401319488 | 94.84 | 51.19% | 48.81% | 98.00% | 99.54% | 497016041 | 15760235 |
| *B. hybridum* | Bhyb_ABR107 | 211826623 | 54.74 | 51.85% | 48.15% | 97.86% | 99.39% | 496297883 | 15731561 |
| *B. hybridum* | Bhyb_ABR112 | 196089207 | 43.66 | 51.71% | 48.29% | 97.83% | 99.53% | 497634766 | 15741915 |
| *B. hybridum* | Bhyb_ABR113_reference | 100954311 | 75.23 | 50.51% | 49.49% | 76.42% | 77.80% | 457070920 | 12145743 |
| *B. hybridum* | Bhyb_ABR115 | 196689399 | 51.33 | 51.50% | 48.50% | 97.75% | 99.55% | 497707739 | 15736340 |
| *B. hybridum* | Bhyb_ABR116 | 329063744 | 83.8 | 51.20% | 48.80% | 97.84% | 99.36% | 497986081 | 15728375 |
| *B. hybridum* | Bhyb_ABR117 | 481874823 | 111.18 | 52.33% | 47.67% | 97.93% | 99.44% | 497410825 | 15746694 |
| *B. hybridum* | Bhyb_ABR118 | 246656632 | 62.89 | 51.62% | 48.38% | 97.91% | 99.58% | 500591498 | 15753066 |
| *B. hybridum* | Bhyb_ABR127 | 249719724 | 63.85 | 51.81% | 48.19% | 97.90% | 99.44% | 495075449 | 15740322 |
| *B. hybridum* | Bhyb_ABR132 | 243776495 | 64.06 | 51.26% | 48.74% | 97.85% | 99.54% | 497570790 | 15746694 |
| *B. hybridum* | Bhyb_ABR134 | 320961775 | 82.56 | 51.41% | 48.59% | 97.84% | 99.67% | 500012976 | 15763421 |
| *B. hybridum* | Bhyb_ABR135 | 172696625 | 45.47 | 51.59% | 48.41% | 97.93% | 99.58% | 499875131 | 15756252 |
| *B. hybridum* | Bhyb_ABR137 | 246059172 | 64.44 | 51.29% | 48.71% | 97.93% | 99.58% | 504849832 | 15756252 |
| *B. hybridum* | Bhyb_ABR138 | 212438881 | 54.9 | 51.77% | 48.23% | 97.89% | 99.56% | 496435662 | 15750677 |
| *B. hybridum* | Bhyb_Adi_P1 | 306789498 | 66.06 | 51.79% | 48.21% | 97.83% | 99.61% | 499078739 | 15757845 |
| *B. hybridum* | Bhyb_ALCU | 178479656 | 46.66 | 51.76% | 48.24% | 97.86% | 99.53% | 497288843 | 15745898 |
| *B. hybridum* | Bhyb_ALFR | 242027116 | 64.3 | 51.68% | 48.32% | 97.96% | 99.60% | 498783822 | 15760235 |
| *B. hybridum* | Bhyb_ALGE | 293737147 | 76.08 | 51.58% | 48.42% | 97.86% | 99.53% | 497772560 | 15746694 |
| *B. hybridum* | Bhyb_ARTA | 123535520 | 28.45 | 51.66% | 48.34% | 97.71% | 99.37% | 494252040 | 15710056 |
| *B. hybridum* | Bhyb_AS-5 | 51256954 | 3.87 | 51.80% | 48.20% | 41.80% | 42.49% | 470813122 | 6565503 |
| *B. hybridum* | Bhyb_AS-6 | 47242739 | 3.85 | 51.72% | 48.28% | 40.64% | 41.55% | 470383480 | 6387885 |
| *B. hybridum* | Bhyb_Bal_P1 | 313931388 | 66.57 | 51.28% | 48.72% | 97.96% | 99.49% | 498823351 | 15753863 |
| *B. hybridum* | Bhyb_BANYA | 123101421 | 32.23 | 51.89% | 48.11% | 97.82% | 99.50% | 495722892 | 15736340 |
| *B. hybridum* | Bhyb_Bd12_2 | 250518202 | 51.45 | 51.94% | 48.06% | 97.88% | 99.40% | 495921811 | 15733154 |
| *B. hybridum* | Bhyb_Bd13_1 | 282418278 | 65.35 | 51.62% | 48.38% | 98.02% | 99.66% | 499630037 | 15778554 |
| *B. hybridum* | Bhyb_Bd14_1 | 300631322 | 66.31 | 51.93% | 48.07% | 97.82% | 99.50% | 497475012 | 15742712 |
| *B. hybridum* | Bhyb_Bd17_2 | 360398638 | 81.3 | 51.61% | 48.39% | 97.92% | 99.46% | 497482766 | 15749084 |
| *B. hybridum* | Bhyb_Bd23_1 | 225422710 | 47.79 | 51.25% | 48.75% | 97.84% | 99.53% | 497566065 | 15745101 |
| *B. hybridum* | Bhyb_Bd25_1 | 311448748 | 66.67 | 51.79% | 48.21% | 97.82% | 99.65% | 498983807 | 15757845 |
| *B. hybridum* | Bhyb_Bd26_1 | 220829686 | 47.38 | 51.53% | 48.47% | 97.91% | 99.56% | 498160906 | 15751473 |
| *B. hybridum* | Bhyb_Bd27_1 | 249355118 | 62.95 | 52.47% | 47.53% | 97.85% | 99.62% | 498743476 | 15754659 |
| *B. hybridum* | Bhyb_Bd4_2 | 245690564 | 51.04 | 51.52% | 48.48% | 97.78% | 99.57% | 498417130 | 15741915 |
| *B. hybridum* | Bhyb_BdTR4F | 359555735 | 79.13 | 51.33% | 48.67% | 97.87% | 99.55% | 498873813 | 15753066 |
| *B. hybridum* | Bhyb_BdTR6g | 329889543 | 83.45 | 50.23% | 49.77% | 97.98% | 99.50% | 499622006 | 15757845 |
| *B. hybridum* | Bhyb_BELC | 239874911 | 64.38 | 51.53% | 48.47% | 97.92% | 99.55% | 497533969 | 15752270 |
| *B. hybridum* | Bhyb_BEMP | 308088499 | 76.8 | 54.89% | 45.11% | 96.19% | 97.46% | 469653348 | 15355616 |
| *B. hybridum* | Bhyb_BG1 | 150686402 | 75.92 | 51.65% | 48.35% | 97.72% | 99.53% | 495930794 | 15731561 |
| *B. hybridum* | Bhyb_BG11 | 300506941 | 69.94 | 52.00% | 48.00% | 97.91% | 99.60% | 499827698 | 15762624 |
| *B. hybridum* | Bhyb_BG16 | 269939246 | 81.27 | 51.70% | 48.30% | 97.95% | 99.62% | 499624012 | 15761828 |
| *B. hybridum* | Bhyb_BG18 | 316877694 | 42.19 | 51.18% | 48.82% | 97.96% | 99.65% | 499579736 | 15772979 |
| *B. hybridum* | Bhyb_BG21 | 334843936 | 14.84 | 51.23% | 48.77% | 90.33% | 92.22% | 498818853 | 14786122 |
| *B. hybridum* | Bhyb_BG23 | 304844691 | 76.19 | 51.87% | 48.13% | 97.86% | 99.60% | 498716935 | 15756252 |
| *B. hybridum* | Bhyb_BG3 | 283204498 | 80.28 | 51.70% | 48.30% | 97.77% | 99.62% | 497865528 | 15747491 |
| *B. hybridum* | Bhyb_BG30 | 322739773 | 75.53 | 51.33% | 48.67% | 97.83% | 99.66% | 499489535 | 15761031 |
| *B. hybridum* | Bhyb_Bhyb_127 | 187168279 | 75.7 | 51.21% | 48.79% | 97.98% | 99.58% | 499667165 | 15706073 |
| *B. hybridum* | Bhyb_Bhyb_30 | 194008623 | 50.47 | 51.60% | 48.40% | 97.91% | 99.61% | 497506973 | 15759438 |
| *B. hybridum* | Bhyb_Bhyb118-5 | 278518793 | 114.58 | 55.07% | 44.93% | 96.14% | 97.29% | 455038004 | 15347651 |
| *B. hybridum* | Bhyb_Bhyb118-8 | 166319636 | 70.94 | 50.93% | 49.07% | 97.82% | 99.58% | 493347351 | 15746694 |
| *B. hybridum* | Bhyb_Bhyb123-1 | 211241814 | 87.97 | 51.30% | 48.70% | 97.87% | 99.55% | 494998184 | 15749880 |
| *B. hybridum* | Bhyb_Bhyb17 | 282320443 | 22.68 | 52.01% | 47.99% | 97.85% | 99.46% | 496649771 | 15740322 |
| *B. hybridum* | Bhyb_Bhyb24 | 243540751 | 59.37 | 51.51% | 48.49% | 98.03% | 99.68% | 502451276 | 15664655 |
| *B. hybridum* | Bhyb_Bhyb26_2 | 284287377 | 73.73 | 51.67% | 48.33% | 97.92% | 99.55% | 498482660 | 15753066 |
| *B. hybridum* | Bhyb_Bhyb26_reference | 193513047 | 50.96 | 54.38% | 45.62% | 96.16% | 97.37% | 464476274 | 15336500 |
| *B. hybridum* | Bhyb_Bhyb30-1 | 194008623 | 26.31 | 51.60% | 48.40% | 97.91% | 99.61% | 497506737 | 15759438 |
| *B. hybridum* | Bhyb_Bhyb50 | 439844879 | 98.27 | 51.50% | 48.50% | 97.97% | 99.63% | 500252644 | 15771386 |
| *B. hybridum* | Bhyb_Bhyb51 | 296755131 | 63.86 | 51.74% | 48.26% | 98.02% | 99.65% | 498961198 | 15776961 |
| *B. hybridum* | Bhyb_Bhyb6 | 369688843 | 83.56 | 51.65% | 48.35% | 97.73% | 99.47% | 497804617 | 15733154 |
| *B. hybridum* | Bhyb_Bhyb7 | 269860579 | 59.27 | 51.64% | 48.36% | 97.91% | 99.42% | 497619589 | 15741119 |
| *B. hybridum* | Bhyb_Bhyb8 | 283294347 | 63.88 | 51.50% | 48.50% | 97.71% | 99.42% | 496115684 | 15725189 |
| *B. hybridum* | Bhyb_BhybCA_UCM | 221090173 | 53.99 | 54.20% | 45.80% | 96.15% | 97.44% | 466580670 | 15346058 |
| *B. hybridum* | Bhyb_BhybEM_UCM | 269211042 | 62.99 | 55.43% | 44.57% | 96.18% | 97.40% | 468203083 | 15349244 |
| *B. hybridum* | Bhyb_Bra117 | 158021506 | 42.01 | 51.83% | 48.17% | 97.85% | 99.56% | 496456966 | 15742712 |
| *B. hybridum* | Bhyb_Bra142 | 292968467 | 74.36 | 51.97% | 48.03% | 97.98% | 99.57% | 498104373 | 15768200 |
| *B. hybridum* | Bhyb_Bra143 | 313250724 | 79.21 | 51.22% | 48.78% | 97.95% | 99.63% | 499611137 | 15766607 |
| *B. hybridum* | Bhyb_Bra151 | 331507219 | 87 | 52.14% | 47.86% | 97.93% | 99.60% | 499166415 | 15761828 |
| *B. hybridum* | Bhyb_Bra293 | 349550281 | 89.88 | 51.63% | 48.37% | 97.90% | 99.60% | 499087628 | 15760235 |
| *B. hybridum* | Bhyb_Bra32 | 267682465 | 70.27 | 51.73% | 48.27% | 97.92% | 99.64% | 498804164 | 15764217 |
| *B. hybridum* | Bhyb_Bra9 | 302754058 | 76.31 | 52.01% | 47.99% | 97.95% | 99.65% | 498044819 | 15770589 |
| *B. hybridum* | Bhyb_CIW1_1 | 326885776 | 71.54 | 51.55% | 48.45% | 97.93% | 99.57% | 498820667 | 15759438 |
| *B. hybridum* | Bhyb_DRAGO881 | 396936184 | 105.09 | 51.57% | 48.43% | 97.93% | 99.58% | 498721354 | 15757845 |
| *B. hybridum* | Bhyb_ECI_reference | 133758011 | 121.85 | 51.36% | 48.64% | 95.62% | 97.16% | 510565984 | 12780549 |
| *B. hybridum* | Bhyb_ESPU | 356309557 | 94.12 | 51.49% | 48.51% | 97.90% | 99.54% | 498570961 | 15754659 |
| *B. hybridum* | Bhyb_FARO | 341002821 | 88.86 | 51.30% | 48.70% | 97.94% | 99.56% | 498087643 | 15758642 |
| *B. hybridum* | Bhyb_GROSE | 216077844 | 56.83 | 52.06% | 47.94% | 97.92% | 99.54% | 497668491 | 15750677 |
| *B. hybridum* | Bhyb_IBD_107 | 323458589 | 73.65 | 51.81% | 48.19% | 97.92% | 99.54% | 498872951 | 15755456 |
| *B. hybridum* | Bhyb_IBD_189 | 300865629 | 70.53 | 51.98% | 48.02% | 97.88% | 99.63% | 499614206 | 15758642 |
| *B. hybridum* | Bhyb_Isk_P1 | 288105858 | 65.74 | 51.23% | 48.77% | 97.87% | 99.63% | 498610559 | 15759438 |
| *B. hybridum* | Bhyb_LLIG | 258856731 | 66.96 | 51.96% | 48.04% | 97.85% | 99.58% | 498768736 | 15753066 |
| *B. hybridum* | Bhyb_MT110_4 | 287067040 | 63.62 | 51.68% | 48.32% | 97.83% | 99.64% | 499348733 | 15755456 |
| *B. hybridum* | Bhyb_MT118_1 | 368513120 | 84.43 | 51.65% | 48.35% | 97.96% | 99.67% | 500009499 | 15771386 |
| *B. hybridum* | Bhyb_PC1_17 | 269495930 | 70.95 | 51.69% | 48.31% | 97.97% | 99.59% | 499042265 | 15763421 |
| *B. hybridum* | Bhyb_PC31_17 | 298405054 | 78.03 | 51.79% | 48.21% | 97.88% | 99.59% | 499728745 | 15755456 |
| *B. hybridum* | Bhyb_PC63_17 | 257231426 | 63.65 | 51.62% | 48.38% | 97.98% | 99.54% | 498562227 | 15760235 |
| *B. hybridum* | Bhyb_PC64_17 | 272831267 | 68.91 | 51.69% | 48.31% | 97.88% | 99.58% | 498990190 | 15752270 |
| *B. hybridum* | Bhyb_PC66_17 | 346712182 | 84.42 | 51.74% | 48.26% | 97.91% | 99.59% | 499577693 | 15758642 |
| *B. hybridum* | Bhyb_PC76_17 | 255385450 | 63.4 | 51.30% | 48.70% | 97.93% | 99.49% | 495972151 | 15749880 |
| *B. hybridum* | Bhyb_PC85_17 | 290101785 | 71.39 | 51.44% | 48.56% | 97.92% | 99.53% | 498274476 | 15751473 |
| *B. hybridum* | Bhyb_PC89_17 | 233702270 | 60.8 | 51.05% | 48.95% | 97.94% | 99.58% | 497561763 | 15757845 |
| *B. hybridum* | Bhyb_PC90_17 | 241248829 | 60.79 | 51.24% | 48.76% | 97.95% | 99.60% | 499942660 | 15759438 |
| *B. hybridum* | Bhyb_PC92_17 | 266073568 | 60.56 | 51.15% | 48.85% | 97.89% | 99.56% | 498648729 | 15751473 |
| *B. hybridum* | Bhyb_POBL | 262219481 | 67.57 | 51.61% | 48.39% | 97.86% | 99.53% | 497215888 | 15743508 |
| *B. hybridum* | Bhyb_PRAT | 356557841 | 93.75 | 51.64% | 48.36% | 97.87% | 99.55% | 500559598 | 15748287 |
| *B. hybridum* | Bhyb_SORB | 287513487 | 74.09 | 51.65% | 48.35% | 97.85% | 99.60% | 498334866 | 15753066 |
| *B. hybridum* | Bhyb_TISC | 287387849 | 73.74 | 51.12% | 48.88% | 97.78% | 99.52% | 498208740 | 15737136 |
| *B. hybridum* | Bhyb_TORO | 290281012 | 76.58 | 51.79% | 48.21% | 97.94% | 99.53% | 497970154 | 15754659 |
| *B. hybridum* | Bhyb-7 | 105877303 | 24.53 | 51.40% | 48.60% | 97.68% | 99.40% | 500411362 | 15718817 |
| *B. hybridum* | Bhyb-AS-1 | 135374226 | 35.66 | 51.60% | 48.40% | 97.82% | 99.44% | 501109133 | 15736340 |
| *B. hybridum* | Bhyb-AS-10 | 44067811 | 26.3 | 51.62% | 48.38% | 91.75% | 93.36% | 496704551 | 14737536 |
| *B. hybridum* | Bhyb-AS-11 | 46147928 | 25.27 | 50.88% | 49.12% | 92.98% | 94.63% | 496874008 | 14940642 |
| *B. hybridum* | Bhyb-AS-12 | 42163362 | 33.63 | 51.92% | 48.08% | 91.56% | 93.30% | 496591821 | 14709659 |
| *B. hybridum* | Bhyb-AS-2 | 43260267 | 23.9 | 51.02% | 48.98% | 92.40% | 94.11% | 496711328 | 14845859 |

**Table S3.** Summary of predicted deleterious polymorphic variants in the *Brachypodium distachyon* – *B. stacei* – *B. hybridum* complex species and *B. hybridum* lineages per megabase (Mb). (A) Proportion (%) of deleterious polymorphic variants by functional category based on the full SnpEff annotation, including all mutation types potentially affecting gene function. (B) Number of potential deleterious variants detected using SnpEff and deleterious variants detected using BAD_Mutations (in parenthesis), per (sub)genome for each progenitor species (*B. distachyon*, *B. stacei*) and for the three *B. hybridum* lineages (Ancient: *B. hybridum*_A, Recent West: *B. hybridum*_RW, Recent East: *B. hybridum*_RE), separately for D- and S-type (sub)genomes.

| Type | *B. distachyon* | *B. stacei* | *B. hybridum_A* | *B. hybridum_RW* | *B. hybridum_RE* |
| --- | --- | --- | --- | --- | --- |
| upstream_gene_variant | 31.69% | 29.81% | 29.36% | 31.41% | 28.02% |
| downstream_gene_variant | 28.53% | 26.83% | 26.88% | 28.02% | 27.66% |
| intergenic_region | 23.46% | 28.88% | 29.90% | 25.43% | 20.32% |
| intron_variant | 8.38% | 7.90% | 7.68% | 8.40% | 13.96% |
| missense_variant | 2.83% | 2.11% | 1.92% | 1.89% | 2.63% |
| 3_prime_UTR_variant | 1.73% | 1.64% | 1.51% | 1.77% | 2.36% |
| 5_prime_UTR_variant | 1.45% | 0.88% | 0.91% | 1.02% | 1.36% |
| 5_prime_UTR_premature_start_codon_gain_variant | 0.15% | 0.12% | 0.14% | 0.15% | 0.19% |
| synonymous_variant | 1.48% | 1.52% | 1.42% | 1.61% | 3.00% |
| disruptive_inframe_deletion | 0.00% | 0.00% | 0.00% | 0.00% | 0.00% |
| initiator_codon_variant | 0.00% | 0.00% | 0.00% | 0.00% | 0.00% |
| intragenic_variant | 0.00% | 0.00% | 0.00% | 0.00% | 0.00% |
| splice_acceptor_variant | 0.01% | 0.01% | 0.01% | 0.01% | 0.01% |
| splice_donor_variant | 0.04% | 0.01% | 0.01% | 0.01% | 0.01% |
| splice_region_variant | 0.24% | 0.23% | 0.22% | 0.24% | 0.45% |
| start_lost | 0.01% | 0.00% | 0.00% | 0.00% | 0.00% |
| start_retained_variant | 0.00% | 0.00% | 0.00% | 0.00% | 0.00% |
| stop_gained | 0.03% | 0.04% | 0.04% | 0.03% | 0.03% |
| stop_lost | 0.01% | 0.00% | 0.00% | 0.00% | 0.00% |
| stop_retained_variant | 0.00% | 0.00% | 0.00% | 0.00% | 0.00% |

(B)
|  | *B. distachyon* | *B. stacei* | *B. hybridum_A* | *B. hybridum_RW* | *B. hybridum_RE* |
| --- | --- | --- | --- | --- | --- |
| (Sub)genomeD | 598.87<br>(235.18) | NA | 1,898.32<br>(797.11) | 803.03<br>(328.75) | 736.43<br>(271.37) |
| (Sub)genomeS | NA | 624.86<br>(248.95) | 2,635.30<br>(1,048.58) | 713.09<br>(283.67) | 728.08<br>(291.09) |

**Table S4.**
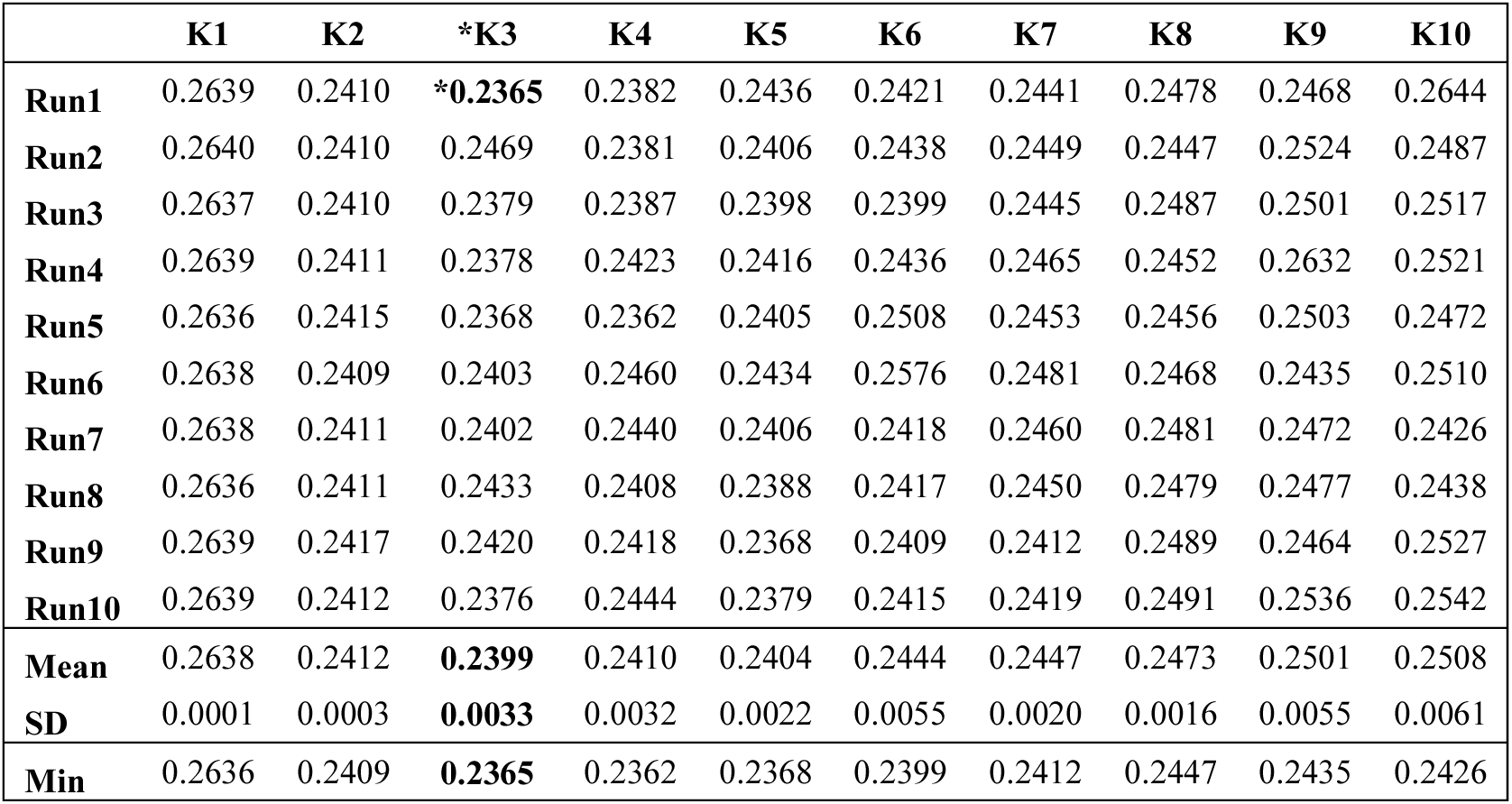
Genomic structure analysis of the studied *Brachypodium hybridum* populations using ADMIXTURE for K=1 to K=10. Cross-validation errors (CVerror) are calculated for each run and each K and mean and standard deviation for all runs of each K. Minimum mean CV error value is highlighted in bold and marked with an asterisk (*).

|  | K1 | K2 | *K3 | K4 | K5 | K6 | K7 | K8 | K9 | K10 |
| --- | --- | --- | --- | --- | --- | --- | --- | --- | --- | --- |
| **Run1** | 0.2639 | 0.2410 | **\*0.2365** | 0.2382 | 0.2436 | 0.2421 | 0.2441 | 0.2478 | 0.2468 | 0.2644 |
| **Run2** | 0.2640 | 0.2410 | 0.2469 | 0.2381 | 0.2406 | 0.2438 | 0.2449 | 0.2447 | 0.2524 | 0.2487 |
| **Run3** | 0.2637 | 0.2410 | 0.2379 | 0.2387 | 0.2398 | 0.2399 | 0.2445 | 0.2487 | 0.2501 | 0.2517 |
| **Run4** | 0.2639 | 0.2411 | 0.2378 | 0.2423 | 0.2416 | 0.2436 | 0.2465 | 0.2452 | 0.2632 | 0.2521 |
| **Run5** | 0.2636 | 0.2415 | 0.2368 | 0.2362 | 0.2405 | 0.2508 | 0.2453 | 0.2456 | 0.2503 | 0.2472 |
| **Run6** | 0.2638 | 0.2409 | 0.2403 | 0.2460 | 0.2434 | 0.2576 | 0.2481 | 0.2468 | 0.2435 | 0.2510 |
| **Run7** | 0.2638 | 0.2411 | 0.2402 | 0.2440 | 0.2406 | 0.2418 | 0.2460 | 0.2481 | 0.2472 | 0.2426 |
| **Run8** | 0.2636 | 0.2411 | 0.2433 | 0.2408 | 0.2388 | 0.2417 | 0.2450 | 0.2479 | 0.2477 | 0.2438 |
| **Run9** | 0.2639 | 0.2417 | 0.2420 | 0.2418 | 0.2368 | 0.2409 | 0.2412 | 0.2489 | 0.2464 | 0.2527 |
| **Run10** | 0.2639 | 0.2412 | 0.2376 | 0.2444 | 0.2379 | 0.2415 | 0.2419 | 0.2491 | 0.2536 | 0.2542 |
| **Mean** | 0.2638 | 0.2412 | **0.2399** | 0.2410 | 0.2404 | 0.2444 | 0.2447 | 0.2473 | 0.2501 | 0.2508 |
| **SD** | 0.0001 | 0.0003 | **0.0033** | 0.0032 | 0.0022 | 0.0055 | 0.0020 | 0.0016 | 0.0055 | 0.0061 |
| **Min** | 0.2636 | 0.2409 | **0.2365** | 0.2362 | 0.2368 | 0.2399 | 0.2412 | 0.2447 | 0.2435 | 0.2426 |

